# Mapping the developing human immune system across organs

**DOI:** 10.1101/2022.01.17.476665

**Authors:** Chenqu Suo, Emma Dann, Issac Goh, Laura Jardine, Vitalii Kleshchevnikov, Jong-Eun Park, Rachel A. Botting, Emily Stephenson, Justin Engelbert, Zewen Kelvin Tuong, Krzysztof Polanski, Nadav Yayon, Chuan Xu, Ondrej Suchanek, Rasa Elmentaite, Cecilia Domínguez Conde, Peng He, Sophie Pritchard, Mohi Miah, Corina Moldovan, Alexander S. Steemers, Martin Prete, John C. Marioni, Menna R. Clatworthy, Muzlifah Haniffa, Sarah A. Teichmann

## Abstract

Recent advances in single cell genomics technologies have facilitated studies on the developing immune system at unprecedented scale and resolution. However, these studies have focused on one or a few organs and were thus limited in understanding the developing immune system as a distributed network across tissues. Here, we profiled prenatal haematopoietic organs, lymphoid organs and non-lymphoid tissues using a combination of single-cell RNA sequencing, paired antigen-receptor sequencing and spatial transcriptomics to reconstruct the developing human immune system. Our analysis revealed the acquisition of immune effector transcriptome profiles in macrophages, mast cells and NK cells from the second trimester, and the transcriptomic changes accompanying the late-stage maturation of developing monocytes and T cells that extended from their organ of origin to peripheral tissues. We uncovered system-wide blood and immune cell development beyond the conventional primary haematopoietic organs. We further identified, extensively characterised and functionally validated the human prenatal B1 cells. Finally, we provide evidence for thymocyte-thymocyte selection origin for αβTCR- expressing unconventional T cells based on TCR gene usage and an *in vitro* artificial thymic organoid culture model. Our comprehensive atlas of the developing human immune system provides both valuable data resources and biological insights that will facilitate cell engineering, regenerative medicine and disease understanding.

**One-Sentence Summary:** By performing a comprehensive single-cell RNA sequencing atlas of human developing immune system together with antigen-receptor sequencing and spatial transcriptomics, we explored the cross-gestation and cross-organ variability in immune cells, discovered system-wide blood and immune cell development, identified, characterised and functionally validated the properties of human prenatal B1 cells and the origin of unconventional T cells.

## Main Text

The human immune system develops across several anatomical sites throughout gestation. Immune cells differentiate initially from extra-embryonic yolk sac progenitors and subsequently from Aorto-Gonad-Mesonephros-derived haematopoietic stem cells (HSCs), before liver and bone marrow take over as the primary sites of haematopoiesis (*1, 2*). Immune cells from these primary haematopoietic sites seed developing lymphoid organs such as the thymus, spleen and lymph nodes, as well as peripheral non-lymphoid organs.

Our understanding of the molecular and cellular basis of immune system development has been based primarily on animal models including rodents. Advances in single cell genomics technologies have enabled developing human organs to be profiled (*3–11*), and revealed some fundamental differences between mice and humans (*1, 7, 9*). However, these studies have focused on one or a few organs, rather than reconstructing the entire immune system as a distributed network across all developing organs. Such a holistic understanding of the developing human immune system would have far-reaching implications for health and disease including cellular engineering, regenerative medicine and a deeper mechanistic understanding of congenital disorders affecting the immune system.

Here we present a cross-tissue single cell and spatial atlas of developing human immune cells across prenatal haematopoietic (yolk sac, liver, bone marrow), lymphoid (thymus, spleen and lymph node) and non-lymphoid peripheral organs (skin, kidney and gut) to provide a detailed characterisation of generic and tissue-specific properties of the developing immune system. We generated new single-cell RNA sequencing (scRNA-seq) data from yolk sac, prenatal spleen and skin, and integrated publicly available cell atlases of 6 more organs, spanning week 4 to week 17 post-conception (*3, 4, 7, 8, 10, 11*). We also generated, for the first time, single-cell γδT cell receptor (γδTCR) sequencing data and additional αβT cell receptors (αβTCR) and B cell receptors (BCR) sequencing data. Furthermore, we integrated the single cell transcriptome profiles with *in situ* tissue location using spatial transcriptomics.

Our study reveals the acquisition of immune effector functions of myeloid and lymphoid lineages from the second trimester, the maturation of developing monocytes and T cells prior to peripheral tissue seeding and system-wide haematopoiesis during human prenatal development. We identify, characterise and functionally validate the properties of human prenatal B1 cells and the origin and selection of unconventional T cells.

### Integrated cross-organ map of prenatal cell states in distinct tissue microenvironments

To systematically assess the heterogeneity of immune cell populations across human prenatal haematopoietic organs, secondary lymphoid organs (thymus, spleen and mesenteric lymph node) and non-lymphoid tissues (skin, kidney and gut), we generated new scRNA-seq data from prenatal spleen, yolk sac and skin which were integrated with a collection of publicly available single-cell datasets from the Human Developmental Cell Atlas initiative (*3, 4, 7, 8, 10, 11*). In total, our dataset comprised samples from 25 embryos/foetuses between 4 to 17 post-conception weeks (pcw) (Fig. 1A), profiled in 221 scRNA-seq libraries. For 61 of these libraries, paired antigen-receptor sequencing data was available for either αβTCR, γδTCR or BCR (Fig. 1B, see Methods). We performed mapping and preprocessing with a unified pipeline, including correction of ambient mRNA expression and filtering of putative contaminant maternal cells (see Methods), yielding a total of 908,178 cells after quality control.

**Fig. 1:**
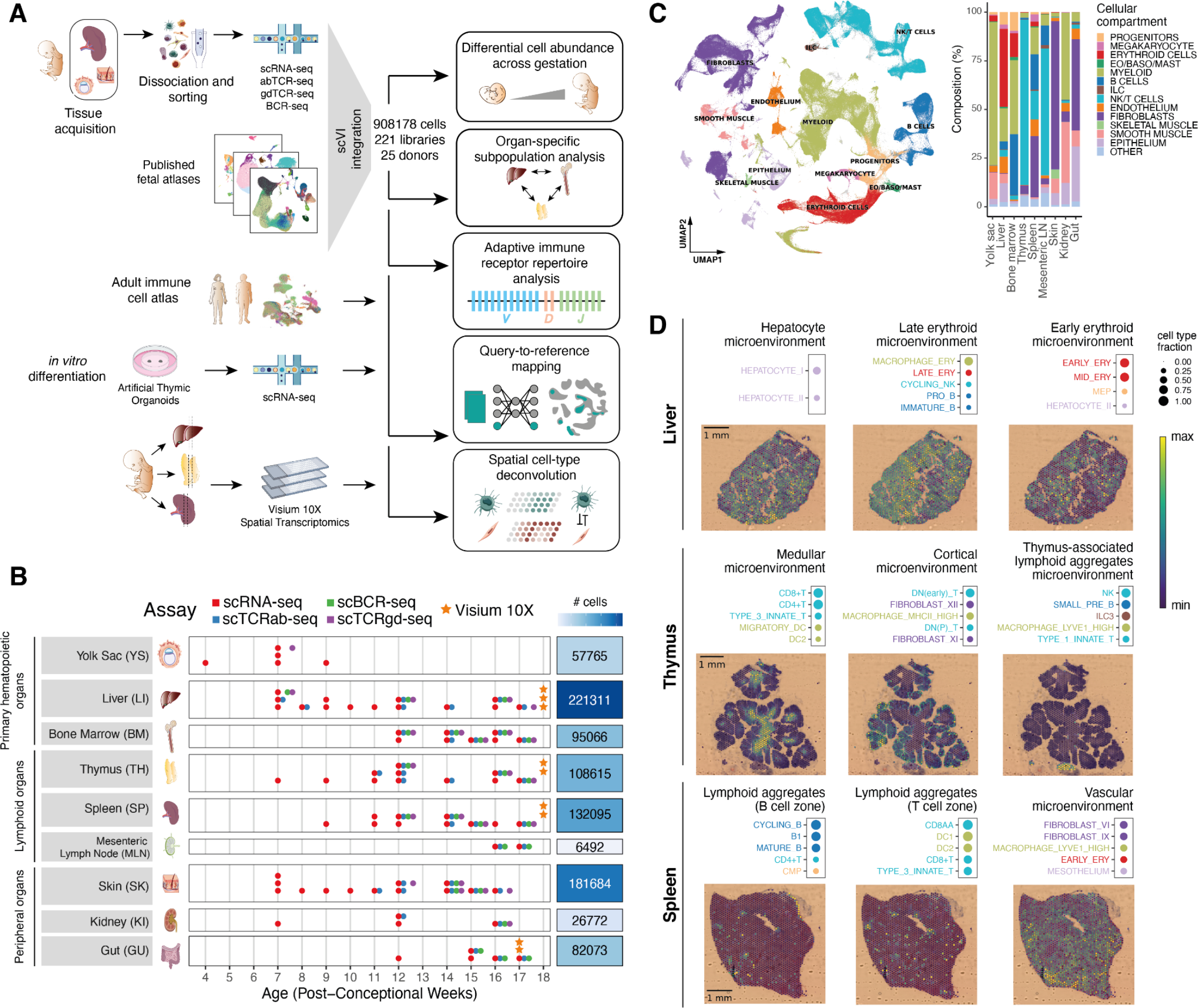
Cross-tissue cellular atlas of the developing human immune system. (A) Overview of study design and analysis pipeline. We generated scRNA-seq and scVDJ-seq data from prenatal spleen, yolk sac and skin which were integrated via scVI with a collection of publicly available single-cell RNA-seq datasets. This cell atlas was used for (i) differential abundance analysis across gestation and organs with Milo, (ii) antigen receptor repertoire analysis with scirpy and dandelion, (iii) comparison with adult immune cells and *in vitro* differentiated cells with scArches and CellTypist, (iv) spatial cell type deconvolution on Visium 10x data of haematopoietic and lymphoid organs using *cell2location*. (B) Summary of analysed samples by gestational stage (x-axis) and organ (y-axis). Colours denote the types of molecular assays performed for each sample. The side bar indicates the total number of cells collected for each organ (after quality control). (C) Left: UMAP embedding of scRNA-seq profiles in prenatal tissues (908178 cells) coloured by broad cellular compartments. Right: barplot of percentage of cells assigned to each broad compartment for each of the profiled organs. Raw cell proportions are adjusted to account for FACS-based CD45 enrichment (see Methods). The category ‘Other’ denotes clusters annotated as low-quality cells. (Eo/Baso/Mast: eosinophils/basophils/mast cells; ILC: Innate Lymphoid Cells; NK: Natural Killer cells). (D) Representative co-localisation patterns identified with non-negative matrix factorisation (NMF) of spatial cell type abundances estimated with cell2location. For each annotated microenvironment, we show (top) a dot plot of relative contribution of cell types to microenvironment (dot size), and (bottom) spatial locations of microenvironments on tissue slides, with the colour representing the weighted contribution of each microenvironment to each spot.

To facilitate the identification of common cell populations and joint analysis of the data, we integrated all libraries using scVI (*12*), to minimise protocol- and embryo-associated variation (<u>fig. S1A</u>) while retaining differences between organs. In keeping with previous single-cell atlases of immune cells of prenatal and adult tissues (*3, 11, 13*), our data captured the emergence of myeloid, lymphoid lineages and closely linked megakaryocytes (MK), erythroid and non-neutrophilic granulocyte lineages (eosinophils, basophils and mast cells) from haematopoietic progenitors (Fig. 1C, left). While haematopoietic and immune cell compartments comprised cells from multiple tissues, variation in cell population representation across organs was notably larger in the non-immune compartment (Fig. 1C, right, <u>fig. S1B</u>, <u>fig. S2</u>). Linking transcriptional phenotypes to paired antigen receptor sequence expression, we have paired αβTCR sequences for 28,739 cells, paired γδTCR for 534 cells, and paired BCR sequences for 14,903 cells (<u>fig. S1C</u>).

We repeated dimensionality reduction and graph-based clustering on subsets of cells from different lineages and used marker gene analysis and comparison with existing cell labels to comprehensively annotate cell types across tissues. In total we defined 127 high quality cell populations (<u>fig. S3</u>, <u>fig. S4</u>, see Methods for detailed description of the annotation workflow). Cross-tissue integration enabled the identification of cell populations that were too rare to be resolved through analysis of datasets from individual tissues, such as *AXL* and *SIGLEC6-* expressing Dendritic Cells (AS DC) (*14*) and Plasma B cells (<u>fig. S4</u>).

To facilitate fast re-use of our atlas for reanalysis of newly collected samples, we make the weights from trained scVI models available to enable mapping of external scRNA-seq datasets using transfer learning with scArches (*15*) (see data and code availability). In this study we demonstrate the capabilities of our atlas as a reference for comparisons between adult and prenatal cells, and prenatal cells with *in vitro-*derived lymphoid cells.

To study the spatial localisations of the cell populations in an early haematopoietic tissue and lymphoid organs critical for T and B cell development, we profiled developing liver, thymus and spleen from 3 donors at 18 pcw with sequencing-based spatial transcriptomics (10x Genomics Visium). Using our multi-organ scRNA-seq dataset as reference, we performed spatial cell type deconvolution with cell2location (*16*) to map cell locations in tissue (<u>fig. S5</u>). We used non-negative matrix factorisation (NMF) of the cell type abundance estimates in tissue spots to identify microenvironments of co-localised cell types in the profiled tissues in an unbiased manner (Fig. 1D, <u>fig. S6</u>, <u>fig. S7</u>, <u>fig. S8</u>).

In the developing liver we recovered expected signatures of tissue-specific parenchymal cells such as hepatocytes. We identified tissue zones corresponding to vasculature with co-localisation of vascular smooth muscle cells, endothelium together with immune cells such as monocytes. In addition, we observed spatial segregation of early and late erythrocytes, suggesting distinctive haematopoietic zones (Fig. 1D, <u>fig. S6</u>).

In the developing thymus we recovered localisation of cell types in known histological structures: developing T cells were largely localised to the thymic cortex, while mature T cells were consistently mapped to the thymic medulla. In addition, in two of the thymic tissue sections, we observed aggregates of lymphoid tissue which we have termed as thymus-associated lymphoid aggregates. Within these, we mapped B cell subsets, innate lymphoid cells and macrophage subtypes (Fig. 1D, <u>fig. S7</u>). In the developing spleen, most of the tissue was highly vascularised. In addition, within splenic lymphoid aggregates, we were able to distinguish B-cell zones and T-cell zones, which were partially co-localising but not completely overlapping (Fig. 1D, <u>fig. S8</u>).

### Heterogeneity of prenatal myeloid cells across organs and gestation

Our cross-tissue embeddings of cellular phenotypes, in-depth cell type annotations, antigen-receptor repertoire data and spatial cell type mappings formed the basis for in-depth comparisons of the cellular compartments across tissues and gestational age. We first examined the main compartments of immune cells in our multi-organ dataset to identify gestation-specific and organ-specific variability within cell populations.

The myeloid compartment captures the development of mononuclear phagocytes, from committed myeloid progenitors to neutrophils, monocytes, macrophages and dendritic cells (<u>fig. S4G-H</u>). DC subsets and stages of neutrophil maturation matched those previously described in human development (*3, 11*). Conversely, our cross-tissue analysis distinguished three distinct subsets of monocytes, which were characterised by differential distribution between prenatal bone marrow and peripheral tissues and by expression of *CXCR4, CCR2* or *IL1B* (*17*). Among macrophages, we identified eight broad macrophage groupings based on transcriptome profile: ‘LYVE1^hi^’ expressing *F13A1, LYVE1* and *SPP1*; ‘Iron-recycling’ expressing the highest levels of ferroportin (*SLC40A1*) and phosphatidylserine receptor *TIMD4* but best characterised by expression of *CD5L*, *VCAM1* and *APOE*; ‘MHC class II^hi’^ expressing the highest levels of *HLA-DRA, HLA-DPA1* and *CLEC7A* amongst macrophages; ‘Kupffer-like’ expressing endothelial transcripts *ENG, KDR* and *CAV*; ‘TREM2’ with expression of microglia-associated transcripts *TREM2* and *P2RY12;* ‘Osteoclasts’ expressing characteristic *MMP9* and *ACP5*; and ‘Proliferating macrophages’ expressing genes associated with cell cycle progression (<u>fig. S4H</u>).

We compared prenatal and adult immune cell populations by mapping a cross-tissue adult dataset of immune cells (*18*) onto our prenatal myeloid reference (<u>fig. S9A-B</u>). We found the transcriptional profiles of DC subsets were conserved between adult and prenatal counterparts (<u>fig. S9C</u>). Adult monocytes were most similar to the IL1B^hi^ and CCR2^hi^ prenatal populations, and we did not observe any CXCR4^hi^ monocytes in non-lymphoid adult tissues (<u>fig. S10</u>). The majority of adult macrophages clustered separately from the prenatal macrophages, with the exception of erythrophagocytic macrophages (<u>fig. S9B-C</u>). This population includes macrophages primarily from the spleen and liver that carry out iron-recycling functions (*18*).

### Proinflammatory macrophages and mast cells in early gestation

We next aimed to quantify changes in cellular composition across gestation. We used differential abundance analysis on cell neighbourhoods with Milo (*19*) to identify regions of the phenotypic manifold where cells become significantly enriched or depleted during gestation and the linear increase in cell abundance with age in different organs (log-fold change, Fig. 2A, <u>fig. S11A</u>).

**Fig. 2:**
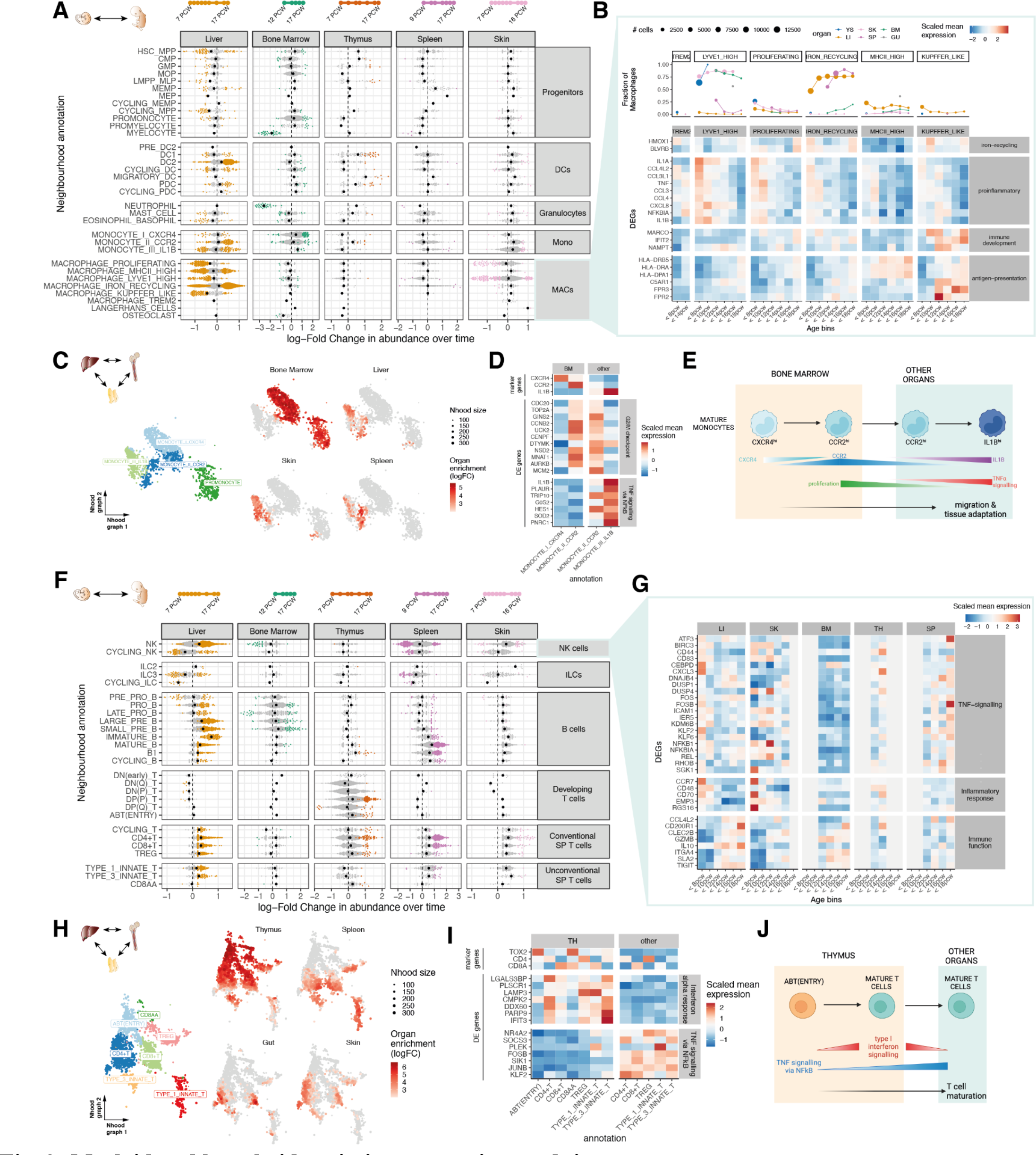
Myeloid and lymphoid variation across time and tissues. (A) Beeswarm plot showing the distribution of log-fold change (x-axis) in cell abundance across gestational stages in neighbourhoods of myeloid cells calculated with Milo. Fold changes of abundance in time of cells from liver (LI), bone marrow (BM), thymus (TH), spleen (SP) and skin (SK) are shown. Neighbourhoods overlapping the same cell population are grouped together (y-axis). Neighbourhoods displaying significant differential abundance at SpatialFDR 10% are coloured. The black point denotes the median log-fold change. The top bar denotes the range of gestational stages of the organ samples analysed. (B) Heat map showing average expression across time of a selection of genes identified as markers of early-specific and late-specific macrophage neighbourhoods. Mean log-normalised expression for each gene is mean-centred and scaled to the standard deviation. Gestational ages are grouped in 5 age bins. Age bins where less than 30 macrophage cells of a given subset were present are not shown. The top panel shows the fraction of all macrophages belonging to the specified macrophage population in each time point and each organ (colour). Point size is proportional to the number of cells in the population. (C) Close-up view of monocytes on Milo neighbourhood embedding of myeloid cells. Each point represents a neighbourhood, the layout of points is determined by the position of the neighbourhood index cell in the UMAP in fig. S4G, the size of points is proportional to the number of cells in the neighbourhood. On the left embedding, neighbourhoods are coloured by the cell population they overlap. On the right embeddings, neighbourhoods are coloured by their log fold change in abundance between the specified organ and all other organs. Only neighbourhoods displaying significant differential abundance (SpatialFDR 10%) are coloured, the rest are grey. (D) Mean expression of a selection of differentially expressed genes between CCR2^hi^ monocytes from bone marrow (BM) and other organs. Log-normalised expression for each gene is mean-centred and scaled to the standard deviation. We show genes upregulated in bone marrow associated with G2/M checkpoint (using MSigDB Hallmark 2020 gene sets) and genes down-regulated in bone marrow associated with TNF signalling. (E) Schematic of the proposed process of monocyte egression from the bone marrow mediated by CXCR4 and CCR2 expression: CXCR4^hi^ monocytes are retained in the bone marrow, until they switch to a proliferative state with increased expression of CCR2, mediating tissue egression. CCR2^hi^ monocytes seed peripheral tissues and then mature further to the periphery-specific IL1B expressing subtype. (F) Beeswarm plot showing the distribution of log-fold change (x-axis) in cell abundance across gestational stages in neighbourhoods of lymphoid cells (as in A). (G) Heat map showing average expression across time of a selection of genes identified as markers of early-specific and late-specific NK neighbourhoods (as in B): NK cells identified in liver and skin before 12 pcw express TNF proinflammatory genes, while expression of immune-effector genes such as cytokines, chemokines and granzyme genes increases after 12 pcw. Age bins where less than 30 NK cells were present in a given organ are greyed out. (H) Close-up view of single-positive T cells on Milo neighbourhood embedding of lymphoid cells. Each point represents a neighbourhood, the layout of points is determined by the position of the neighbourhood index cell in the UMAP in fig. S4I (as in C). (I) Mean expression of a selection of differentially expressed genes between single-positive T cells from thymus (TH) and other organs. We show genes downregulated in thymus associated with TNF signalling (using MSigDB Hallmark 2020 gene sets) and genes upregulated in thymus associated with interferon alpha response. (J) Schematic of the proposed mechanism of thymocyte maturation and egression from thymus mediated by type I interferon signalling and NF-kB signalling.

This analysis reaffirmed well-known compositional shifts that happen during gestation. For example, myeloid progenitor cells decrease in the liver but increase in the bone marrow, recapitulating the transition from liver to bone marrow haematopoiesis with increasing gestation age. DCs increase in proportional abundance across multiple tissues as previously described in the liver and bone marrow (*3*). Notably, for several cell populations, we found some neighbourhoods enriched across gestation and other neighbourhoods depleted across gestation, suggesting evolving transcriptional heterogeneity during development. This was especially evident in the macrophage compartment in the skin and liver (Fig. 2B) with a large fraction of neighbourhoods overlapping the LYVE1^hi^ and proliferating macrophages enriched in early gestation. Differential expression analysis (see Methods) revealed upregulation of a proinflammatory gene signature with chemokines and cytokines specific to early stages in all macrophage subtypes across tissues (Fig. 2B, <u>fig. S11B</u>). TNF and NF-ᴋB have been implicated in lymphoid tissue organogenesis (*20*) and the chemokines noted here have been associated with angiogenesis in the context of cancer (*21–23*) and non-cancerous tissues (*24*). Conversely, a large fraction of neighbourhoods overlapping the iron-recycling and MHCII^hi^ macrophages populations were enriched in later stages of gestation. We found that these subpopulations upregulate genes encoding for scavenger receptors, complement components, antibacterial and antiviral defence components, as well as antigen-presentation genes (Fig. 2B, <u>fig. S11C</u>, table S1). Similar to the early signature, the upregulation of these genes was notable across macrophage subsets. In parallel to macrophages, we observed similar transcriptional heterogeneity during gestation in mast cells (Fig. 2A). Specifically, early mast cells in yolk sac, liver and skin displayed a similar proinflammatory phenotype as early embryonic macrophages, characterised by expression of *TNF*, NF-ᴋB subunits, as well as chemokines associated with endothelial cell recruitment *CXCL3*, *CXCL2* and *CXCL8* (*23*) (<u>fig. S12</u>).

Taken together the differential transcriptional profile during early organ development suggests that early macrophages and mast cells may contribute to angiogenesis, tissue morphogenesis and homeostasis as previously reported (*25–27*), prior to adopting traditional immunological functions. Notably, acquisition of macrophage antigen presentation properties (e.g. MHCII upregulation) between 10 and 12 pcw coincides with expansion of adaptive lymphocytes across organs (<u>fig. S1E</u>) and development of lymphatic vessels and lymph nodes (*28*).

### Dissemination of monocytes from bone marrow to peripheral organs

We then explored organ-specific transcriptional signatures in myeloid cells using differential abundance analysis on cell neighbourhoods, testing for organ-specific enrichment (see Methods, <u>fig. S13A</u>). Within monocytes, CXCR4^hi^ monocytes were enriched amongst bone marrow cells, while IL1B^hi^ were significantly enriched in peripheral organs. Within CCR2^hi^ monocytes, we distinguished bone marrow-specific and peripheral organ-based subpopulations (Fig. 2C). Bone marrow CCR2^hi^ monocytes expressed proliferation genes, while peripheral organ CCR2^hi^ monocytes upregulated IL1B and other TNF-alpha signalling genes characteristic of IL1B^hi^ monocytes, in differential expression analysis (see Methods; Fig. 2D, <u>fig. S13B</u>, table S2). This indicates that a CXCR4^hi^ to CCR2^hi^ transition accompanies monocyte egress from the bone marrow to seed peripheral tissue, and CCR2^hi^ monocytes further mature into IL1B^hi^ monocytes in peripheral tissue (Fig. 2D-E). In mouse bone marrow, interactions between monocyte CXCR4 and stromal cell CXCL12 retain monocytes *in situ* until CCR2/CCL2 interactions predominate, potentially enabling monocyte egress (*17*). In our data, we observed CXCL12 expression in bone marrow fibroblasts and osteoblasts (<u>fig. S13C</u>). In addition, the CXCR4^hi^ to CCR2^hi^ monocytes transition was not observed in the developing liver (<u>fig. S13D</u>), in keeping with reports that alternative mechanisms of monocyte retention and release operate in murine developing liver (*29*).

### Heterogeneity of prenatal lymphoid cells across organs and gestation

The lymphoid compartment captures the developmental process of T and B cells from progenitors to mature cells, together with innate lymphoid cells (ILCs) and natural killer (NK) cell subsets (<u>fig. S4I-L</u>). It is worth noting that the unconventional T cell subtypes contain both T cells with paired αβTCR and those with paired γδTCR (<u>fig. S1C</u>), which we will discuss in detail later.

Mapping adult cells onto our prenatal lymphoid reference, innate lymphoid cells (NK cells, ILC3) displayed high similarity between adult and prenatal counterparts (<u>fig. S14A-B</u>). Amongst adult T cells, naive T cell populations and Tregs closely matched prenatal conventional T cells, while resident and effector memory T cells did not have a developmental equivalent (<u>fig. S14C</u>). Notably, we found none of the adult T cells were transcriptionally matched to prenatal unconventional T cells (Type 1 and Type 3 Innate T). Adult B cell progenitors and plasma B cells matched to the corresponding prenatal subsets. Adult mature B cell populations (naive B cells, memory B cells and germinal centre B cells) were matched to prenatal Mature B cells, although we did not observe expression of memory or germinal centre B cell markers in prenatal Mature B cells (<u>fig. S14D</u>). In contrast, none of the adult B cells were transcriptionally matched to prenatal putative B1 cells (<u>fig. S14C</u>).

### Innate to adaptive lymphocyte transition across gestation

Differential abundance analysis across gestation identified a broad shift from innate to adaptive immune populations, with mature B and T cell populations becoming enriched at later stages of gestation in multiple tissues (Fig. 2G, <u>fig. S15A</u>). While neighbourhoods enriched in early gestation were mostly composed of ILCs and NK cells, these cell populations also included neighbourhoods enriched at later time points. Genes involved in inflammatory response, including TNF-signalling, were overexpressed in <12 pcw liver and skin NK cells, although late splenic NK cells also expressed these genes. Conversely, late NK cells across organs over- expressed genes involved in cytokine signalling, complement genes and granzyme genes (Fig. 2G, <u>fig. S15B-C</u>, table S3). Similarly to the macrophages, these results suggest a progressive development of immune-effector function of NK cells.

### TNF and NF-ᴋB signals accompany post-thymic T cell egress

We next tested for organ-specific cell neighbourhoods in the lymphoid compartment. As expected, we found that B cell progenitors were largely enriched in the liver and bone marrow, while developing double negative (DN) and double positive (DP) T cells were enriched in the thymus. Neighbourhoods specific to peripheral organs were restricted to mature cell states (<u>fig. S16A</u>). Focusing on the mature T cells, while certain populations were exclusively enriched in the thymus (ABT(entry), CD8AA), we found that neighbourhoods of conventional and unconventional T cells could be subdivided into a subset enriched in thymus and other subsets enriched in peripheral organs (specifically, spleen, gut and skin) (Fig. 2H). To further investigate the gene expression signature of thymus-specific mature T cell subpopulations, we performed differential expression analysis between each cell type from thymus and all other organs (see Methods). We found that thymic mature T cells over-expressed genes involved in interferon alpha signalling while mature T cells from other organs had higher expression of genes in TNF and NF-ᴋB signalling (Fig. 2I, <u>fig. S16B</u>, table S4). Both pathways have been implicated in last stages of functional maturation of murine T cells in the thymus, before emigration out of the thymus (*30, 31*). Our data showed that in addition to the increase in type 1 interferon- and NF-ᴋB-signalling accompanying ABT(entry) to thymic mature T cells, expression of NF-ᴋB signalling genes continued to increase when mature T cells migrate out to peripheral tissues (Fig. 2J).

### System-wide blood and immune cell development

While examining the distribution of various cell types across different organ systems, we were surprised to find that lineage-committed haematopoietic progenitors were present in almost all the prenatal organs profiled. We quantified the abundance of B progenitors, megakaryocyte/erythrocyte progenitors, myeloid progenitors and T progenitors, from different donors across all organs. This revealed an abundance of B progenitors in almost all prenatal organs in more than one donor and to a lesser extent megakaryocyte/erythroid progenitors in developing spleen and skin and myeloid progenitors in thymus, spleen, skin and kidney (Fig. 3A). In contrast to the above three lineages, T cell progenitors were highly restricted to the thymus. This may reflect the more stringent requirement for a T cell-nurturing microenvironment that imposes rigorous central tolerance through the expression of diverse peripheral antigens and is not completely unexpected in view of the absence of T cells in children with congenital athymia (*32*). This finding of B, megakaryocyte/erythrocyte and myeloid progenitors across lymphoid and non-lymphoid organs challenges the conventional dogma of haematopoiesis being restricted to developing liver and bone marrow between 7-19 pcw (*33*), and suggest that other organs can also support haematopoiesis during prenatal development.

**Fig. 3.**
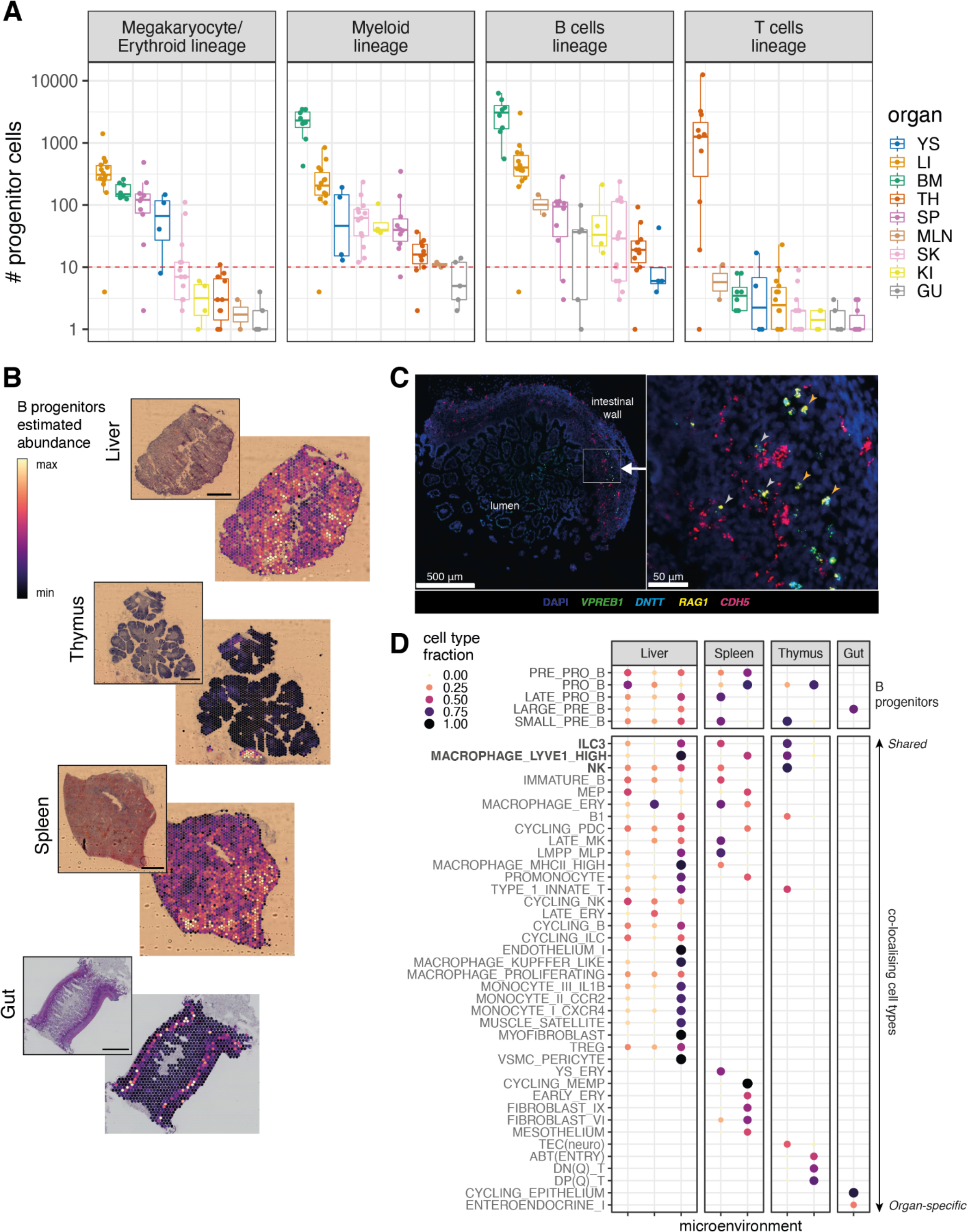
System-wide blood and immune cell development. (A) Boxplots of the number of progenitor cells in all donors across organs. Each point represents a donor, colour- coded by organ (YS: yolk sac; LI: liver; BM: bone marrow; TH: thymus; SP: spleen; MLN: mesenteric lymph node; SK: skin; GU: gut; KI: kidney). Here we treat counts less than 10 (red dash line) as potential technical artefacts. B_prog: B lineage progenitors, including Pre-pro B, Pro B, Late Pro B, Large Pre B and Small Pre B cells; M/E_prog: Megakaryocyte/Erythrocyte progenitors, including MEMP, cycling MEMP and MEP; MYE_prog: myeloid progenitors, including CMP, GMP, MOP, promonocyte, promyelocyte and myelocyte; T_prog: T progenitors, including DN(P), DN(Q), DP(P), DP(Q). Boxes capture the first to third quartile of the cell number and whisks span a further 1.5x interquartile range on each side of the box. (B) Scaled sum of abundances of B progenitor cell types estimated with cell2location, shown on representative slides for each organ. The corresponding H&E staining for each slide is shown on the left. The scale bar represents the length of 1 mm. (C) Multiplex smFISH staining with DAPI, *CDH5* for endothelial cells, and *VPREB1*, *DNTT*, *RAG1* for B progenitors in the human prenatal intestine at 15 pcw. Left panel shows a zoomed-out view with the area of interest boxed in white. Right panel shows a detailed view of the area of interest. Grey arrows point to B progenitors associated with blood vessels and orange arrows point to B progenitors away from blood vessels. (D) Cell type contributions to microenvironments containing B progenitors in different organs identified with non- negative matrix factorisation (NMF) of spatial cell type abundances estimated with cell2location. The colour and the size of the dots represent the relative fraction of cells of a type assigned to the microenvironment.

In addition, we noticed that for all the progenitor lineages observed in the peripheral organs, cells of different developmental stages were simultaneously present (f<u>ig. S1</u>7A-B), further supporting the notion of haematopoiesis beyond fetal liver and bone marrow. For example, we found B cells along the entire developmental path from Pre-pro B cells to Small Pre B cells in thymus, spleen, lymph node, skin, gut and kidney. On the other hand, we only observed Pre-pro B cells in yolk sac, which is consistent with the literature that full B cell lymphopoiesis does not happen until later in embryonic life (*34*).

We then focused on the B lymphopoiesis given its particular widespread nature compared to the other lineages. Using cell2location (*16*) on Visium spatial transcriptomic data, we found that B progenitors were mainly localised in the submucosa of the gut, in the thymus-associated lymphoid aggregates, and proximal to the lymphoid aggregates in the spleen (<u>fig. S18B</u>) in addition to their expected presence in the developing liver (Fig. 3B, <u>fig. S18A</u>). The presence of B progenitors in non-haematopoietic organs was further validated through single-molecule fluorescence *in situ* hybridisation (smFISH) staining. The existence of cells simultaneously expressing *VPREB1*, *RAG1* and/or *DNTT* in the prenatal gut and spleen (Fig. 3C, <u>fig. S18C</u>) confirmed the presence of B progenitors in these samples. Expression of *CDH5* was used to locate the endothelial cells in blood vessels. While some of the B progenitors in the prenatal gut were associated with blood vessels, many could be detected outside the blood vessels (Fig. 3C), supporting the conclusion that B cells develop in prenatal peripheral organs.

The widespread nature of B lymphopoiesis suggests that the cellular environments supporting B cell development are much more widely available than previously thought. To better understand the relevant supporting cell types, we utilised the Visium data to identify cells co-localising with B progenitors across different organs. We identified that ILC3, LYVE1^hi^ Macrophage and NK cells co-localised with B progenitors in different microenvironments across liver, spleen and thymus, whereas other co-localising cell types were organ-specific (Fig. 3D). We then used CellPhoneDB (*35*) to explore the possible cell-cell interactions between B progenitors with ILC3, LYVE1^hi^ Macrophage and NK cells in liver, spleen and thymus (<u>fig. S18D</u>). In addition to the previously described CXCL12-CXCR4 interaction in murine studies (*36, 37*), our analysis identified many additional novel interactions that may inform efforts to generate and engineer B cells *in vitro*. It is worth noting that while many previous studies on B lymphopoiesis-supporting microenvironment have focused on stromal cells (*38*), the co-localisation and predicted interactions between B progenitors and ILC3, macrophages and NK cells highlight that cells from other haematopoietic lineages might also play a role in supporting B cell development.

### Identification of putative prenatal B1 cells

We explored the transcriptional heterogeneity within prenatal non-progenitor B cells, which have productive light chains of B cell receptors (BCR) and low *IL7R* expression compared to B cell progenitors (<u>fig. S19A</u>). Subclustering analysis of non-progenitor B cells enabled the identification of Immature B, Mature B, Cycling B, Plasma B cells and putative B1 cells (Fig. 4A). Immature B cells were characterised by higher expression of *CD19*, *CD24* and *VPREB3*. Mature B, Cycling B, Plasma B and putative B1 cells were functionally mature B cells, all of which expressed MS4A1 except Plasma B (*MS4A1*^lo^ and expressing *CD38*, *SDC1* and *JCHAIN*) (<u>fig. S19B</u>). Cycling B cells were additionally marked with *MKI67* (<u>fig. S19B</u>). The putative B1 cells had the highest expression of *CD27*, *CD5* and *SPN* (CD43) consistent with the surface markers previously reported in mouse and postnatal human putative B1 cells identified by flow cytometry (*39–41*). In addition, we identified *CCR10* as a highly-specific novel potential surface marker expressed in a sub-cluster of B1 cells (Fig. 4A).

**Fig. 4.**
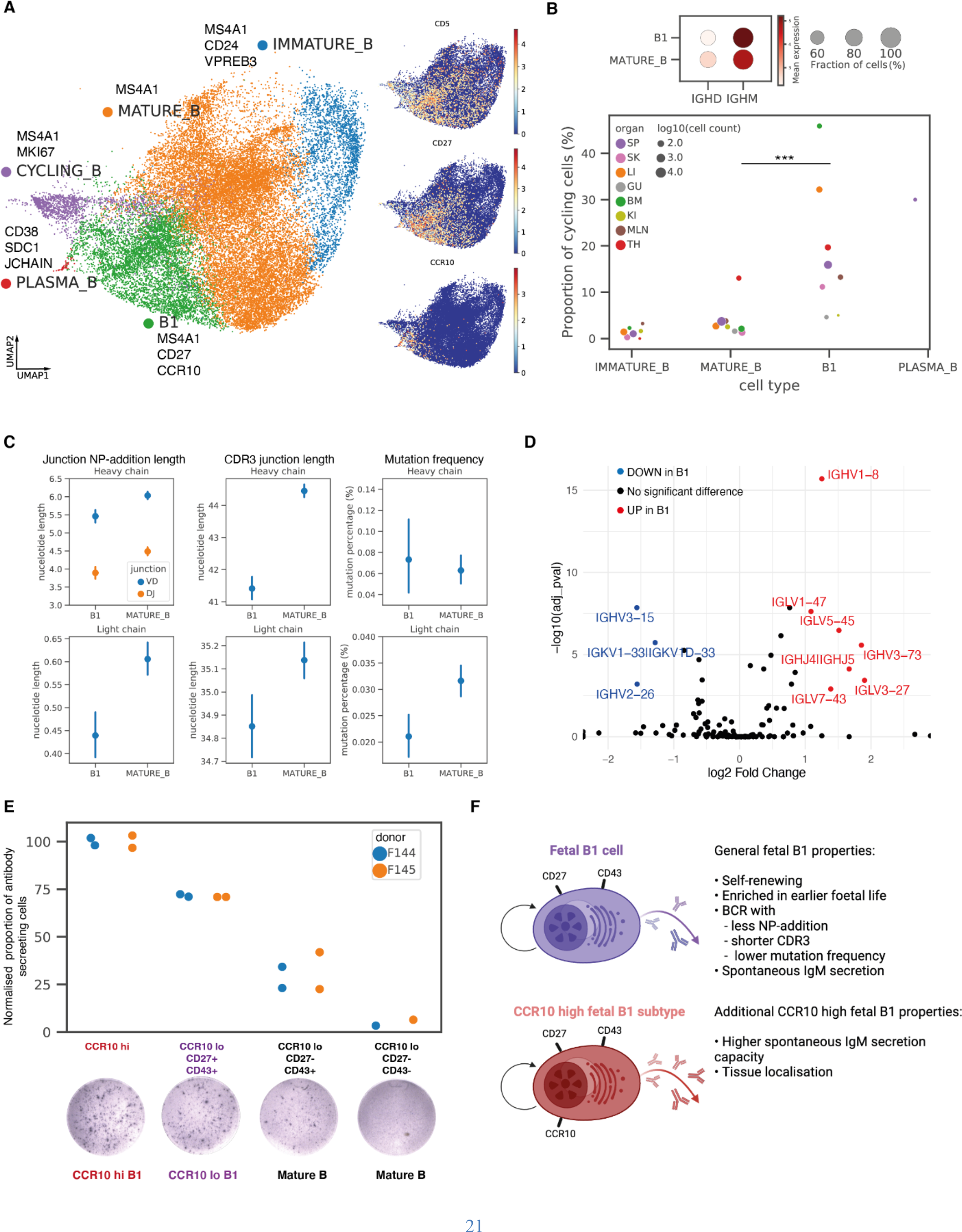
Identification of putative prenatal B1 cells. (A) Left: Close-up view of non-progenitor B cell populations on UMAP embedding of all lymphoid cells (as shown in fig. S4I), with marker genes listed next to the name of each cell type. Right: expression of B1 marker genes overlaid onto the same UMAP. (B) Top: dot plot of *IGHM* and *IGHD* expressions in B1 and Mature B cells, with colour of the dots representing the mean expression and size representing the fraction of cells expressing the gene. Bottom: cycling cell proportions within each B cell subtype, coloured by organs, with dot sizes representing log10(cell count) and only dots with at least 10 cells are shown. A significant difference in cycling proportions was observed between Mature B cells and B1 cells after controlling for organs and gestational age with logistic regression (***p-value < 10^−10^). (C) Point plots of NP-addition length, CDR3 junction length, and mutation frequency in BCR heavy chains or light chains in B1 cells and Mature B cells, with points representing mean expression values and lines representing 95% confidence intervals. Apart from heavy chain mutation frequency comparison, all other characteristics showed statistically significant differences with p-values < 0.001 from t-tests. (D) Volcano plot summarising results of BCR heavy and light chain V, J gene segment usage comparison between B1 and Mature B cells. Y-axis is the -log10(adjusted p-value) and the x-axis is log2(fold change) computed with the chi-squared test. Adjusted p-values were calculated using the Benjamini-Hochberg procedure. (E) Normalised proportions of antibody-secreting cells in different sorted B cell fractions of the ELISpot experiments, coloured by donor. Each point represents a reaction well. The proportions of antibody-secreting cells were normalised against the average proportion in CCR10^hi^ wells for each donor to remove donor-specific effects. A representative well image for each sorted fraction is shown on the bottom. (F) Schematic illustration summarising the features of all human prenatal B1 cells, and additional features specific to CCR10^hi^ prenatal B1 cells.

To further validate the identity of B1 cells, we evaluated the characteristics described for murine B1 cells, including self-renewal (*42, 43*), high IgM and low IgD expressions (*44*), emergence in early development (*45*), low levels of non-templated nucleotide insertions in their BCRs (*46, 47*), and spontaneous antibody secretion (*42*).

We inferred self-renewal capacity of the putative B1 cells by assessing their expression of the cell cycle gene *MKI67*. As our dataset integration was performed via scVI without explicit cell cycle effect regression, all the cycling Immature B, Mature B, B1 and Plasma B cells were clustered together as Cycling B. We used logistic regression to predict the specific cell identity of each of the cells within the ‘Cycling B cells’ group (<u>fig. S20A</u>). This enabled us to calculate the percentage of cycling cells (defined as cells with non-zero *MKI67* expression) within Immature B, Mature B, B1 and Plasma B cells respectively (Fig. 4B, bottom). The proportion of cycling B1 cells was significantly higher than cycling Mature B cells (p-value < 10^−10^) after controlling for organs and gestational age in a logistic regression model, supporting that our putative B1 cells have the capacity for self-renewal.

B1 cells had lower RNA expression of *IGHD* and higher expression of *IGHM* compared to Mature B cells (Fig. 4B, top), another feature present in mouse B1 cells. In addition, we observed the highest frequency of B1 cells in early embryonic stages, and these were gradually replaced by other subsets of non-progenitor B cells over time. The ratio of B1 to Mature B cells showed a general decrease from first to second trimester across most organs except the thymus (<u>fig. S20</u>B) where B1 cells persisted, consistent with a previous report of shared phenotype between thymic B cells and B1 cells (*48*).

Next, we analysed non-templated nucleotide insertions in the BCR of our putative B1 cells. Both N/P-additions and CDR3 junctions in heavy and light chains were shorter in B1 cells compared to Mature B cells (Fig. 4C). Moreover, a much lower mutation frequency was observed in the light chains of the B1 cells compared to that in Mature B cells, although this was not statistically significant for the heavy chain. It is worth noting that the average mutation frequencies observed in prenatal B1 cells were much lower than that in adult B cells (*18, 49*). We next examined the V(D)J usage within different B cell subtypes along the development path (<u>fig. S20C</u>). Both prenatal B1 and Mature B cells exhibited a varied BCR repertoire with minimal clonal expansion (<u>fig. S20D</u>). Comparing the V(D)J usage between B1 and Mature B cells, we found preferential usage of *IGHV1-8, IGHV3-73*, *IGLV1-47, IGLV5-45, IGLV3-27* and *IGLV7-43* in B1 cells in contrast to *IGHV3-15, IGHV2-26* and *IGKV1-33|IGKV1D-33* in Mature B cells (Fig. 4D).

One of the defining characteristics of B1 cells is the capacity for spontaneous antibody secretion capacity in the absence of external stimulation (*42*). We sorted B cell subsets, rigorously excluding plasma cells by gating on CD20^hi^/CD38^lo-int^ cells (see Methods, with sorting strategy shown in <u>fig. S20E</u>), and observed spontaneous IgM secretion from FACS isolated splenic B1 cells using an Enzyme-Linked Immune absorbent Spot (ELISpot) assay. The normalised antibody-secreting spot counts were higher in the two sorted B1 fractions than the two sorted Mature B fractions, with the CCR10^hi^ B1 fraction showing the highest spot counts (Fig. 4E). We further explored the potential role of *CCR10* in prenatal B1 cells and observed the expression of one of its ligands *CCL28* in bone marrow stroma (chondrocyte and osteoblast), gut epithelium, and keratinocytes and melanocytes in skin (<u>fig. S20F</u>). This supports a role for *CCR10* in tissue localisation of prenatal B1 cells.

Overall, our scRNA-seq, paired V(D)J sequencing data as well as the functional assay we have performed provide the first extended characterisation of human prenatal B1 cells (Fig. 4F).

### Human unconventional T cells are trained by thymocyte-thymocyte selection

The mature T cell compartment consisted of conventional T cells (CD4+T, CD8+T and regulatory T (Treg) cells) and unconventional T cells. The origin of the latter in humans is poorly understood. Unconventional T cells expressed the key innate marker *ZBTB16* (PLZF) (*50*) (<u>fig. S21A</u>), and could be further separated into three different subtypes, *RORC* and *CCR6* expressing Type 3 Innate T cells, *EOMES* and *TBX21* expressing Type 1 Innate T cells and *PDCD1* expressing CD8AA cells (<u>fig. S4L</u>), corresponding respectively to TH17-like cells, NKT-like cells and CD8αα^+^ T cells previously described in Park et al. 2020 (*7*).

Type 1 and 3 Innate T cells were widespread across all organs, while CD8AA cells were more restricted to thymus (<u>fig. S2</u>). The proportions of unconventional T cells among all mature T cells exhibited a decreasing trend from 7-9 pcw to 10-12 pcw across most of the organs surveyed here (<u>fig. S21B</u>). Notably, the thymus at the earliest stage (7-9 pcw) produced almost exclusively unconventional T cells. The proportions of Type 1 and 3 Innate T cells were almost negligible in postnatal thymus (using dataset from (*7*)), while CD8AA T cell abundance rebounded in paediatric age groups before a further decline in adulthood (<u>fig. S21B</u>). This suggests that Type 1 and 3 Innate T cells, but not CD8AA, are developmental-specific unconventional T cells, and we could not identify an adult counterpart to these cell states in the cross-tissue atlas from (*18*) (<u>fig. S14C</u>).

We explored the spatial locations of mature T cell subtypes within the developing thymus to interrogate the mechanisms regulating their negative selection. Using factor analysis on cell type abundances deconvolved from Visium spatial data, we found that mature T cells segregated in two microenvironments in the thymic medulla (<u>fig. S21C</u>). Conventional CD4+T and CD8+T cells co-localised with mTEC(II) and mTEC(III) close to the inner medulla, while CD8AA and Type 1 Innate T cells co-localised with DC1 closer to the cortico-medullary junction (<u>fig. S21D- E</u>). Treg and Type 3 Innate T cells were located within both microenvironments. The co-localisation of CD8AA and DC1 observed with Visium here is also in agreement with what we showed previously with smFISH (*7*). These findings suggest that, in contrast to conventional T cells, CD8AA and Type I Innate T cells likely undergo distinct negative selection processes mediated by DCs rather than mTECs and they might also be involved in DC activation as previously suggested (*7*).

Single cell sequencing of γδTCR and αβTCR was performed on a subset of samples to characterise antigen-receptor repertoires in unconventional T cells (Fig. 1B). The vast majority of unconventional T cells expressed paired αβTCR but some of these cells expressed paired γδTCR (Fig. 5A). The majority of the γδT cells expressed *TRGV9* and *TRDV2*, consistent with previous reports (*51, 52*) (Fig. 5B). However, there was also a large proportion of γδT cells, particularly those of CD8AA and Type 3 Innate T cell subtypes, expressing *TRGV8* and *TRGV10* instead of *TRGV9*. The full repertoire of γδT cells within the three unconventional subtypes is shown in Fig. 5B with asterisk marks highlighting the gene segments showing differential usage across the three cell subtypes. Overall, the γδTCR showed a relatively restricted repertoire and substantial clonal expansion, with the largest clonotype containing 27 cells from all unconventional T cell subtypes and shared across multiple organs and donors (<u>fig. S22B</u>).

**Fig. 5.**
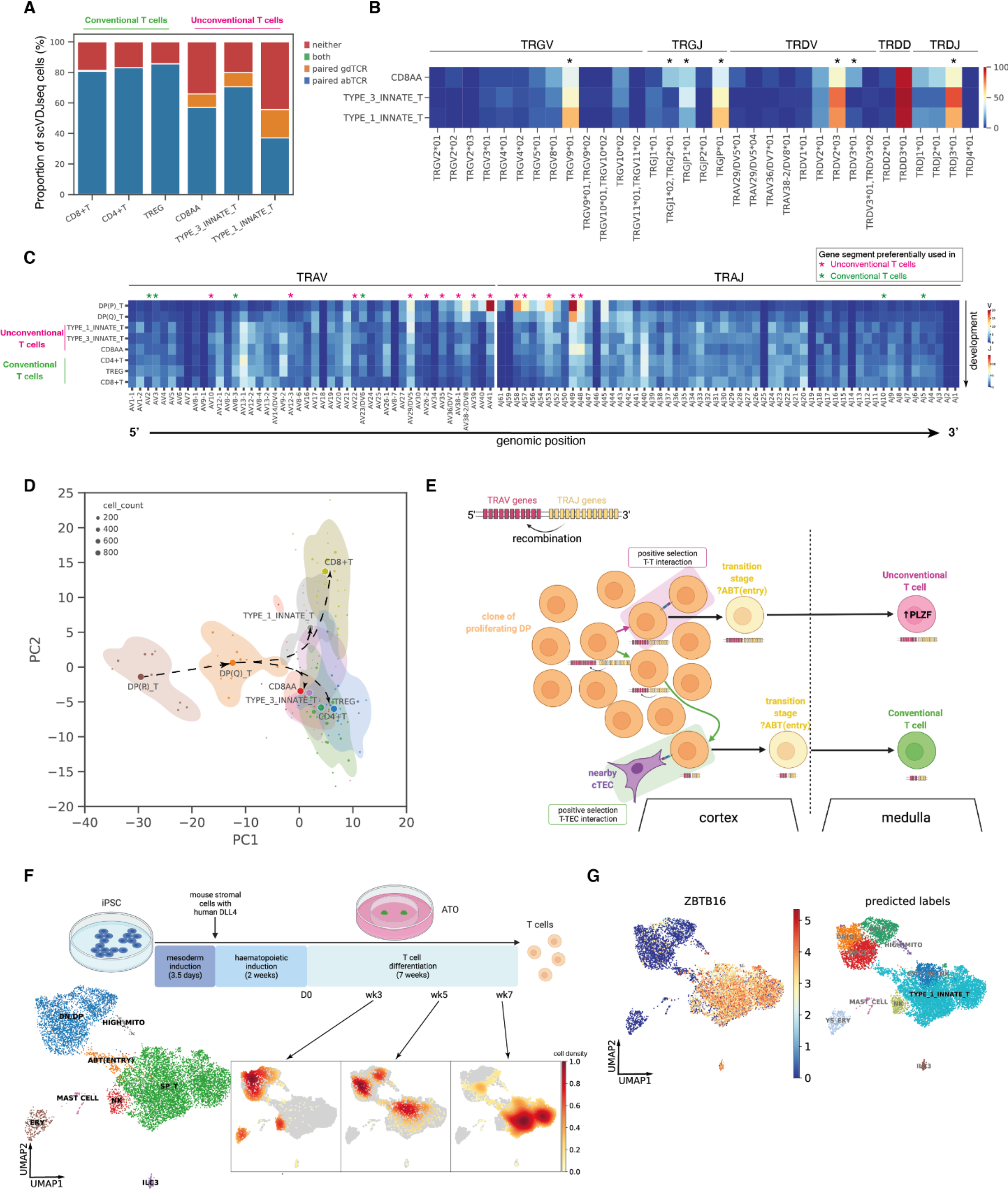
Deep characterisation of human unconventional T cells. (A) Proportions of cells expressing paired γδTCR, paired αβTCR, or both or neither. The proportions were calculated over cells that have had both single cell αβTCR and γδTCR sequencing. Only 3 cells expressed both paired αβTCR and γδTCR (two from Type 1 Innate T cells and one from CD8+T), potentially due to a technical artefact. The cells that expressed neither paired αβTCR nor paired γδTCR could be due to dropouts in single cell TCR sequencing as over 50% of these contained orphan VDJ or VJ chains of αβTCR or γδTCR (fig. S22A). (B) Heat map showing the percentage of each TRGV, TRGJ, TRDV, TRDD and TRDJ gene segment present in different T cell subtypes. Differential usage between cell subtypes was computed using the chi-squared test and gene segments with adjusted p-values < 0.05 are highlighted with *. Adjusted p-values were calculated using the Benjamini-Hochberg procedure. (C) Heat map showing the proportion of each TRAV and TRAJ gene segment present in different T cell subtypes. The usage of each gene segment in unconventional T cells was compared with that in conventional T cells using the chi-squared test. Adjusted p-values were calculated using the Benjamini-Hochberg procedure. Gene segments with adjusted p-value < 0.05 are marked with *, with magenta * indicating preferential usage in unconventional T cells and green * indicating preferential usage in conventional T cells. (D) Principal components analysis plot summarising TRAV, TRAJ, TRBV, TRBJ gene segment usage proportion in different T cell subtypes. Each dot represents a sample, with the size of the dot representing its cell count. Samples with less than 20 cells of a given cell type were excluded. The centroid of each cell type is shown as a filled circle with a white ring, and 80% confidence contours are shown around the centroids. Arrows represent the proposed developmental trajectories. (E) Schematic illustration showing the T-T training origin of unconventional T cells in contrast to the T-TEC training origin of conventional T cells. (F) Top: schematic diagram showing the experimental set-up of T cell differentiation from human iPSCs in Artificial Thymic Organoids (ATO). Cells were harvested at week 3, 5 and 7 of T cell differentiation phase. Bottom left: UMAP visualisation of different cell types in the ATO. Bottom right: density plots of cells from each time point were overlaid on the same UMAP plot. (G) Left: predicted annotations from logistic regression overlaid on the same UMAP plot as in Fig. 5F; right: *ZBTB16* expression pattern overlaid onto the same UMAP plot.

Prenatal unconventional T cells expressed a varied αβTCR repertoire (Fig. 5C for TCRα, <u>fig. S22C</u> for TCRβ) with minimal clonal expansion (<u>fig. S22D</u>), unlike the well-described unconventional T cells (e.g. type I NKT and MAIT cells) found in adults (*53*). V-J gene usage in TCRα was previously observed to have a strong association with T cell developmental timing in both mice (*54*) and humans (*7*). Specifically, double positive (DP) T cells tend to use proximal TRAV, TRAJ gene segments, whereas mature T cells tend to use more distal pairs, which is explained by the processive depletion of proximal segments in V-J gene recombination (*54*). Our αβTCR results revealed for the first time that the V-J gene usage of abTCR-expressing unconventional T cells lies in between that of DP cells and conventional T cells (Fig. 5C). We found that the genes preferentially used in unconventional T cells were proximal on the genome while genes preferentially used in conventional T cells were located in the distal ends of the genome (Fig. 5C). Principal component analysis of TCR repertoire of different cell types placed unconventional T cells closer to quiescent double positive (DP(Q)) T cells compared to conventional T cells, with Type 1 Innate T cells lying between DP(Q) to CD8+T cells, and Type 3 Innate T and CD8AA lying in between DP(Q) and CD4+T and Treg cells (Fig. 5D). Such a TCR V(D)J-gene usage pattern suggests that unconventional T cells are developmentally closer to DP cells (Fig. 5E) and undergo fewer recombinations before positive selection. Our findings on the developmental proximity of unconventional αβT cells to DP cells support thymocyte-thymocyte (T-T) training as a mechanism accounting for the origin of unconventional T cells.

Previous studies have suggested that these PLZF-expressing unconventional T cells might have originated from positive selection on neighbouring T cells (*50, 55–57*), in contrast to conventional T cells arising from positive selection on cortical thymic epithelial cells (cTEC). Post β-selection, DP T cells undergo proliferation prior to recombination of TCRα (*58, 59, 7*). Each DP cell is hence surrounded by many neighbouring DP cells from the same clone. It is therefore theoretically conceivable that it requires less physical migration and hence is quicker for a DP T cell to receive positive signalling from a neighbouring DP T cell rather than having to migrate to meet a nearby cTEC. Therefore, our finding of the TCR usage in unconventional T cells being closer to DP cells is in agreement with the T-T origin hypothesis.

To test our hypothesis for T-T rather that cTEC-mediated selection of unconventional T cells during development, we differentiated induced pluripotent stem cells (iPSCs) into mature T cells using the artificial thymic organoid (ATO) protocol described in Montel-Hagen et al. 2019 (*60*). Importantly, there were no human thymic epithelial cells present in the ATO system, with NOTCH signalling provided by hDLL4-transduced mouse stromal cells. scRNA-seq analysis of differentiated cells harvested at week 3, 5 and 7 from two iPSC lines (see Methods) confirmed that the *in vitro* culture system recapitulated T cell development from DN/DP, to ABT(ENTRY) then to single positive mature T cells (SP_T) (Fig. 5F, <u>fig. S23A-B</u>). Interestingly, the single positive mature T cells differentiated *in vitro* using the ATO protocol were dominated by *ZBTB16*-expressing unconventional T cells (Fig. 5G). Both label transfer with logistic regression (Fig. 5G) and distance-based similarity scores computed on merged embeddings (<u>fig. S23D</u>) showed that the *in vitro* SP_T were most similar to *in vivo* Type 1 Innate T cells. Hence, our *in vitro* experiments support the T-T origin hypothesis of unconventional T cells, given the absence of human thymic epithelial cells in the ATO system.

## Discussion

Our study provides the most comprehensive single-cell dataset of the developing human immune system, spanning over 900k single cell profiles from 9 tissues, encompassing over 100 cell states. Compared to previous multi-organ developmental atlases (*9*) we increased coverage of developmental organs and gestation stages, sequencing depth, and generated paired BCR, αβTCR, and γδTCR datasets. Furthermore, we demonstrate the utility of scRNA-seq reference from dissociated cells to delineate tissue organisation and cellular communication in spatial transcriptomics data, providing the first proof-of-concept study of localisations of immune cells across prenatal tissues. We provide the research community with preprocessed data and pre-trained models (scVI and CellTypist models) to facilitate alignment of new data to our dataset, which will streamline future expansion and analysis of human developmental atlases.

Our cross-organ analysis unravelled novel biological discoveries with important implications. Firstly, we identify for the first time in humans that the transcriptome of macrophages, mast cells and NK cells suggests that they acquire immune effector functions between 10 and 12 pcw. Their transcriptomic signatures prior to this time point suggest a role in tissue morphogenesis, for example expansion of the vascular network both to supply the metabolic demands of growing organs and to establish tissue-based immune networks from circulating cells. Such roles are in line with what has been previously suggested for murine macrophages (*61*), and might explain why these cells are evident in early development. It is unclear what molecular signals initiate the transcriptional switch in these immune cells. However, the coincidental development of lymphatic vessels and lymph nodes around 12 pcw (*28*) raises the possibility of a potential role for the lymphatic system in this switch. Secondly, we observed conserved processes of proliferation and maturation for monocytes and T cells prior to their migration from the bone marrow and thymus respectively into peripheral tissues.

Thirdly, in contrast to the previous dogma of haematopoiesis being restricted to the yolk sac, liver and bone marrow during human development, we show system-wide haematopoiesis, in particular B lymphopoiesis, across all sampled peripheral organs. It is possible that haematopoiesis is supported to varying levels in prenatal organs, including the adrenal gland (*9*), prior to the onset of functional organ maturation as exemplified by the fetal liver, which transitions from a haematopoietic to a metabolic organ during development. The potential for other lymphoid and non-lymphoid organs to support haematopoiesis is evidenced by the re-emergence of extramedullary haematopoiesis in adults, primarily in pathological settings (*62*–*65*), as well as the recent description of B lymphopoiesis in murine and non-human primate meninges (*66–68*). The quantitative and qualitative contribution of immune cells developed in peripheral organs to prenatal immunity will require further study.

Finally, we identified and functionally validated the properties of human prenatal innate-like B and T cells. To our knowledge, our study provides the most extensive characterisation of human B1 cells. Our *in vivo* αβTCR V(D)J usage patterns and *in vitro* T cell differentiation data proposes T-T selection underpinning unconventional T cell development. It is possible that during early development, cTECs, especially in the primordial thymus, are functionally immature and provide inefficient T-TEC interactions. As such, T-T selection predominates and is supported by our observation of abundant unconventional T cells compared to conventional T cells at this gestation stage. Further studies are required to establish if B1 cells arise from different progenitors (lineage model) (*69–71*) or from the same progenitors but with different signalling (selection model) (*72, 73*), similar to the conventional and unconventional T cell model proposed here. Notably, both innate-like B and T cells were abundant during early development and their precise role at this developmental time-point warrants further investigations. B1 cells are reported as weakly self-reactive, and involved in removal of cell debris (*42, 41*). Both innate-like B and T cells are also reported to react more quickly to antigens (*41, 42, 53*), while B1 cells have also been hypothesised to have an immune regulatory function in suppressing potential self-reactive T cell responses (*42*). Their debris removal capability, antigen reactivity as well as regulatory capacity may confer these prenatal innate-like B and T cells tissue homeostatic and important immunological functions.

In summary, our comprehensive atlas of the developing human immune system provides valuable resources and biological insights to facilitate *in vitro* cell engineering, regenerative medicine and enhance our understanding of congenital disorders affecting the immune system.

## Acknowledgements

We gratefully acknowledge the Sanger Flow Cytometry Facility, Newcastle University Flow Cytometry Core Facility, Sanger Cellular Generation and Phenotyping (CGaP) Core Facility, and Sanger Core Sequencing pipeline for support with sample processing and sequencing library preparation. We thank Gay Cooks lab (A. Montel-Hagen, S. Lopez and G. Cooks) for their kind help in setting up the ATO experiments; and R. Lindeboom, C. Talavera-Lopez and K. Kanemaru for helpful discussions; J. Eliasova and BioRender.com for graphical illustrations. The human embryonic and fetal material was provided by the MRC–Wellcome Trust-funded Human Developmental Biology Resource (HDBR; http://www.hdbr.org). We are grateful to the donors and donor families for granting access to the tissue samples. This publication is part of the Human Cell Atlas: www.humancellatlas.org/publications. We acknowledge Wellcome Trust Sanger Institute as the source of HPSI0114i-kolf_2 and HPSI0514i-fiaj_1 human induced pluripotent cell lines which were generated under the Human Induced Pluripotent Stem Cell Initiative funded by a grant from the Wellcome Trust and Medical Research Council, supported by the Wellcome Trust (WT098051) and the NIHR/Wellcome Trust Clinical Research Facility, and acknowledge Life Science Technologies Corporation as the provider of Cytotune.

## Funding

We acknowledge funding from the Wellcome Human Cell Atlas Strategic Science Support (WT211276/Z/18/Z), CZI Seed Networks for the Human Cell Atlas (Thymus award), MRC Human Cell Atlas award and Wellcome Human Developmental Biology Initiative. M.H. is funded by Wellcome (WT107931/Z/15/Z), The Lister Institute for Preventive Medicine and NIHR and Newcastle Biomedical Research Centre. S.A.T. is funded by Wellcome (WT206194), the ERC Consolidator Grant ThDEFINE. C.S. is supported by a Wellcome Trust Ph.D. Fellowship for Clinicians. Z.K.T and M.R.C are supported by a Medical Research Council Research Project Grant (MR/S035842/1). M.R.C. was supported by an NIHR Research Professorship (RP-2017-08-ST2- 002) and a Wellcome Investigator Award (220268/Z/20/Z).

## Author contributions

Conceptualisation: SAT, MH, MRC, CS, ED. Data curation: CS, ED, IG. Formal Analysis: ED, CS, IG, LJ, JEP, VK, ZKT, KP, CX, NY, RE, CDC, PH, CM, JCM. Funding acquisition: SAT, MH. Methodology: CS, IG, RAB, ES, JE, MM, ASS. Project administration: SAT, MH, CS, ED, IG. Software: ED, KP, ZKT, CX, MP. Supervision: SAT, MH, MRC. Validation: CS, SP, NY, OS. Visualisation: CS, ED, NY. Writing - original draft: CS, ED, MH, LJ, IG, VK, NY, KP, ZKT, SP. Writing - review and editing: all authors.

## Competing interests

In the past 3 years, S.A.T. has consulted for Genentech and Roche and sits on Scientific Advisory Boards for Qiagen, Foresite Labs, Biogen, and GlaxoSmithKline and is a co-founder and equity holder of Transition Bio. The remaining authors declare no competing interests.

## Materials and Methods

### Tissue acquisition and processing

All human developmental tissue samples used for this study were obtained from the MRC– Wellcome Trust-funded Human Developmental Biology Resource (HDBR; http://www.hdbr.org) with written consent and approval from the Newcastle and North Tyneside NHS Health Authority Joint Ethics Committee (08/H0906/21+5).

All tissues were processed into single cell suspensions immediately upon receipt. Tissue was first minced in a tissue culture dish using scalpel. It was then digested with type IV collagenase (final concentration of 1.6 mg/ml; Worthington) in RPMI (Sigma-Aldrich) supplemented with 10% fetal bovine serum (FBS; Gibco), at 37°C for 30 min with intermittent agitation. Digested tissue was then passed through a 100 µm cell strainer and cells were pelleted by centrifugation at 500 *g* for 5 min at 4°C. Cells were then resuspended in 5 ml red blood cell lysis buffer (eBioscience) and left for 5-10 min at room temperature. It was then topped up with flow buffer (PBS containing 2% (v/v) FBS and 2 mM EDTA) to 45 ml prior to cell counting and antibody staining. Single cell suspensions were generated from 65 samples across yolk sac, liver, bone marrow, spleen, thymus, kidney and skin of 21 donors. The ages of the donors spanned from 4 pcw (post conception weeks) to 17 pcw.

### Single-cell RNA sequencing experiment

Dissociated cells were stained with anti-CD45 antibody and DAPI prior to sorting. Sorting by flow cytometry was performed with BD FACSAria Fusion Flow Cytometer. CD45 positive fraction was sorted from DAPI^−^CD45^+^ gate, and CD45 negative fraction was sorted from DAPI^−^ CD45^−^ gate. CD45 gating was contiguous so that no live cells were lost in sorting.

For scRNA-seq experiments, either Chromium single cell 3’ reagent kit or Chromium single cell V(D)J kits from 10x Genomics were used. Unsorted, or DAPI^−^CD45^+^ or DAPI^−^CD45^−^ FACS-isolated cells were loaded onto each channel of the Chromium chip following the manufacturer’s instructions before droplet encapsulation on the Chromium controller. Single-cell cDNA synthesis, amplification, gene expression (GEX) and targeted B cell receptor (BCR) and T cell receptor (TCR) libraries were generated. Targeted enrichment for γδTCR was done following the TCR enrichment protocol from 10x with customised primers binding to the constant region of the TRD and TRG genes as described in Mimitou et al. 2019 (*75*). Primers are listed below:

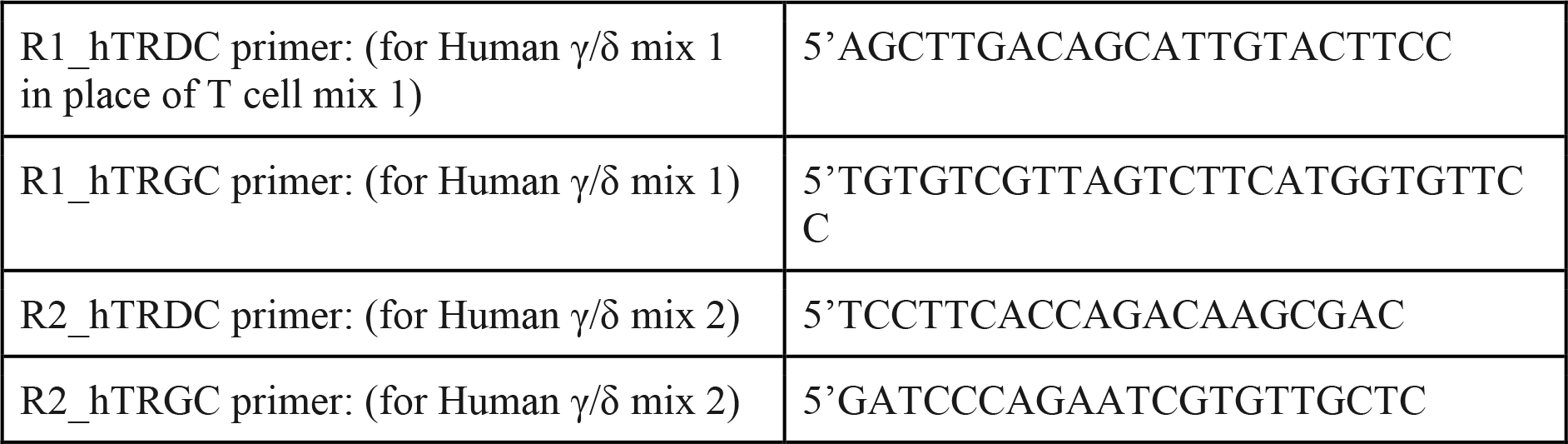

Sequencing was performed on the Illumina Novaseq 6000 system. The gene expression libraries were sequenced at a target depth of 50,000 reads per cell using the following parameters: Read1: 26 cycles, i7: 8 cycles, i5: 0 cycles; Read2: 91 cycles to generate 75-bp paired-end reads. BCR and TCR libraries were sequenced at a target depth of 5000 reads per cell.

### Cell cultures for artificial thymic organoid (ATO)

MS5 line transduced with human DLL4 was obtained from Gay Crooks’ lab (UCLA) as a gift. The MS5-hDLL4 cells were cultured in DMEM (Gibco) with 10% FBS. Two iPSC lines were used in this study. Cell lines HPSI0114i-kolf_2 (Kolf) and HPSI0514i-fiaj_1 (Fiaj) were obtained from the Human Induced Pluripotent Stem Cell initiative (HipSci: www.hipsci.org) collection. All iPSC lines were cultured on vitronectin (diluted 1:25 in PBS; Gibco) coated plates, in TeSR-E8 media (Stemcell Technologies).

We followed the PSC-ATO protocol as described in Montel-Hagen et al. 2019 (*60*). iPSC cells were harvested as single cell suspension and seeded (3 million cells per well) in GFR reduced Matrigel (Corning) -coated 6-well plates in X-VIVO 15 media (Lonza), supplemented with rhActivin A, rhBMP4, rhVEGF, rhFGF (all from R&D Systems), and ROCK inhibitor (Y27632; LKT Labs) on Day -17, and only rhBMP4, rhVEGF and rhFGF on Day -16 and Day -15. Cells were harvested 3.5 days later, and sorted by flow cytometry for CD326^−^CD56^+^ human embryonic mesodermal progenitors (hEMPs).

Isolated hEMPs were combined with MS5-hDLL4 at a ratio of 1:50. Two or three cell-dense droplets (5 x 10^5^ cells in 6 µl haematopoietic induction medium) were deposited on top of an insert in each well of a 6-well plate. Haematopoietic induction medium composed of EGM2 (Lonza) supplemented with ROCK inhibitor and SB blocker (TGF-β receptor kinase inhibitor SB-431542; Abcam) was added into the wells outside the inserts so that the cells sat at the air-liquid interface. The organoids were then cultured in EGM2 with SB blocker for 7 days (Day -14 to Day -7), before the addition of cytokines rhSCF, rhFLT3L, rhTPO (all from Peprotech) between Day -6 to Day 0. These two weeks formed the haematopoietic induction phase. On Day 1, media was changed again to RB27 (RPMI supplemented with B27 (Gibco), ascorbic acid (Sigma-Aldrich), penicillin/streptomycin (Sigma-Aldrich) and glutamax (Thermo Fisher Scientific)) with rhSCF, rhFLT3L and rhIL7. The organoids can be maintained in culture for 7 more weeks in this medium.

For dissociation and checking of ATO, a cell scraper was used to detach ATOs from cell culture insert membranes and detached ATOs were then submerged in cold flow buffer. Culture inserts were washed and detached ATOs were pipetted up and down to form single cell suspension before passing through a 50 µm strainer. Cells were then stained with designed panels of antibodies and analysed by flow cytometry. FACS sorting was done at the same time and live human (DAPI^−^anti-mouse CD29^−^) cells were sorted for week 3 ATO cells, and live (DAPI^−^) cells were sorted for week 5 and week 7 ATO cells before loading onto each channel of the Chromium chip from Chromium single cell V(D)J kit (10x Genomics).

### Visium

OCT embedded fresh frozen samples were used for 10x Genomics Visium, and samples were processed following manufacturer’s instructions. All tissues were sectioned with a thickness of 15 µm. Tissue optimisation was then performed with 18-min permeabilisation time for fetal spleen and liver, and 24-min for fetal thymus. The spatial gene expression library was then generated following the manufacturer’s protocol. All images for this process were scanned using a Zeiss AxioImager system.

### Single molecule fluorescence *in situ* hybridisation (smFISH)

The smFISH technique RNAscope was performed on spleen and gut sections, using the RNAScope 2.5 LS multiplex fluorescent assay (ACD, Bio-Techne) on the automated Leica BOND RX system (Leica). Cryosections were 10 µm-thick and placed on superfrost plus slides. Prior to running RNAscope probes of interest, positive and negative control probes were used for optimisation of these tissues. Following optimisation, OCT embedded fresh frozen fetal gut and spleen sections were pretreated offline for 15 minutes with chilled 4% paraformaldehyde and dehydrated through an ethanol series (50%, 70%, 100%, 100% ethanol), before processing on the Leica BOND RX with protease IV for 30 minutes at room temperature. Slides were stained for DAPI (nuclei) and three or four probes of interest, with fluorophores opal 520, opal 570, opal 650 and atto 425. DAPI was used at 1:50,000 concentration; opals at 1:1000 and atto 425 at 1:400 concentration. The fetal gut sections were then imaged on a Perkin Elmer Opera Phenix High Content Screening System with water immersion at 20x magnification.

Due to high levels of endogenous autofluorescence, we imaged the spleen sections with a confocal microscope (Leica SP8) with 40X 1.3NA oil immersion objective. Emission spectral filters were set for DAPI, opal 520 (*VPREB1*), opal 570 (*RAG1*), opal 650 (*CDH5*). Images were processed with Fiji as follows. All channels were subjected to 2D Gaussian filtering (sigma = 0.5 pixels). The DAPI channel was flat field corrected (biovoxxel – pseudo flat field correction plugin) with a rolling ball radius of 200 pixels. The large image was cropped around the tissue area to remove flat field corrections edges.

### scRNA-seq analysis

#### Preprocessing

The gene expression data was mapped with cellranger 3.0.2 to an Ensembl 93 based GRCh38 reference (10x-distributed 3.0.0 version). Ambient RNA was removed with cellbender v0.2.0 (*76*). Low quality cells were filtered out (minimum number of reads = 2000, minimum number of genes = 500, Scrublet (v0.2.3) (*77*) doublet detection score < 0.4).

In order to identify possible maternal contamination, the samples were pooled on a per-donor basis and processed with souporcell (v.2.4.0) (*78*). The common GRCh38 variants file (SNPs with ≤ 2% frequency from 1k genomes) provided by souporcell authors was used. The pipeline was run twice, setting the number of genotype clusters to 1 and 2 to obtain models for no maternal contamination and possible maternal contamination. The better of these models was identified via BIC (Bayesian Information Criterion), calculated using the formula below:

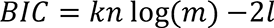

Whereby *k* is the number of genotype clusters set for each souporcell run, *n* denotes the number of loci used for genotype deconvolution, *m* is the cell count for a given donor, and *l* is the log likelihood obtained after running the pipeline with each *k*. In two donors (F19 and F37), the BIC was smaller when *k* = 2. The cells with the minor genotype were identified as possible maternal contaminants, which mainly consisted of NK cells, monocytes, mature B and T cells. The cells of minor genotype from the remaining donors were further screened for similar cell compositions, and a further donor (F33) was identified with possible maternal contamination.

For these three donors, cells from the minor genotype were excluded from the downstream analysis.

#### Data integration and annotation

Data normalisation and preprocessing were performed using the Scanpy workflow (*79*). We normalised raw gene read counts by sequencing depth in each cell (*scanpy.pp.normalize_per_cell,* with parameters *counts_per_cell_after=10e4*) and performed *ln(x)+1* transformation. Expression levels reported in this manuscript refer to normalised and log-transformed gene read counts. We then selected highly variable genes (HVG) for joint embedding by dispersion (*scanpy.pp.highly_variable_genes* with parameters *min_mean = 0.001*, *max_mean = 10*). We considered the 10x chemistry (5’ and 3’) and the donor ID for each cell as the technical covariates to correct for. We performed dimensionality reduction and batch correction using the scVI model (*12*) as implemented in scvi-tools (*80*). For model specification and training we used the recommended parameters to enable scArches mapping (*dropout_rate = 0.2*, *n_layers = 2*). To verify conservation of biological variation after integration, we collected and harmonised the available cell type labels from the published datasets (66% of cells) and quantified the agreement between labels across different datasets in the cell clusters identified post-integration, using the Normalized Mutual Information (NMI) score, as implemented in *scikit-learn* (*81*). The model was trained on raw counts of the 7500 most highly variable genes, excluding cell cycle genes and TCR/BCR genes (as defined by Park et al. (*7*)) with 20 latent dimensions. These parameters (number of HVGs, number of latent dimensions, exclusion/inclusion of cell cycle and TCR/BCR genes) were picked through a parameter sweep, focused on maximising the NMI between clusters after embedding and pre-existing cell type label annotation (data not shown). Unless otherwise specified, cell clustering was performed using the Leiden algorithm (*82*) with resolution = 1.5 on a *k*-nearest neighbour graph with *k* = 30. To verify that our cell type clusters were robust to the choice of integration method, we performed in parallel integration on the full dataset using BBKNN (*83*) as previously described (*7*) (<u>fig. S24A</u>). We found that clustering post-integration both with scVI and BBKNN was consistent with previous annotations (<u>fig. S24B</u>).

To annotate fine cell populations across tissues, we clustered cells in the scVI latent space and preliminarily assigned cells to broad lineages examining expression of marker genes and assigning putative cell labels based on previous annotations (we propagated existing cell type labels to unannotated cells by taking the most abundant label in the *k*-nearest neighbours for each unannotated cell). For each broad lineage we repeated scVI integration and clustering as described above and defined further subsets (see hierarchy in <u>fig. S3</u>). Leiden clusters for the highest resolution subsets (Stroma, megakaryocyte/erythroid, progenitors, lymphoid, myeloid) were annotated manually, using marker panels shown in <u>fig. S4</u>. A common subset of progenitor cells was included in scVI embeddings for all haematopoietic-derived cell subsets (megakaryocyte/erythroid, lymphoid, myeloid, NK/T), to allow feature selection and dimensionality reduction to capture the differentiation process of different lineages. A distinct embedding of progenitor cells was then used to finely annotate these cell populations (<u>fig. S4E- F</u>). We verified that refined annotations were highly consistent with unsupervised clustering post-integration on the full dataset both with scVI and BBKNN (<u>fig. S24C</u>).

After full annotation 23,156 cells (2.5% of total) were assigned to low quality clusters. These comprised doublet clusters, maternal contaminants clusters and clusters displaying a high percentage of reads from mitochondrial genes.

#### Differential abundance analysis

We tested for differences in cell abundances associated with gestational age or organ using the Milo framework for differential abundance testing (*19*), with the python implementation milopy (https://github.com/emdann/milopy). Briefly, we subsetted the dataset subset to cells from libraries obtained with CD45^+^ FACS sorting, CD45^−^ FACS sorting or no FACS sorting. In addition, we excluded FACS sorted samples for which we weren’t able to recover the true sorting fraction quantification. In total we retained 228,731 lymphoid cells and 214,874 myeloid cells. To further minimise the differences in cell numbers driven by differences in FACS sorting efficiency, we calculated a FACS sorting correction factor for each tissue sample *s* sorted with gate *i* (where *i* is either CD45^+^ or CD45^−^):

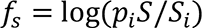

where *p*_*i*_ represents the true proportion of cells from gate *i* in the tissue samples from the same organ and donor, *S* represents the total number of cells recovered from both CD45^+^ and CD45^−^ gates for this organ and donor and *S*_*i*_ represents the number of cells recovered in gate *i*. For the unsorted samples we set *f*_*s*_ = 0. Encoding the true proportions of CD45^+/–^ cells with *f*_*s*_ reduced the proportion of false positives that were found without regressing out the effect of FACS sorting, or when encoding the effect of sorting as a label rather than a proportion (where CD45^+^ = 1, CD45^−^ = -1, unsorted = 0) (<u>fig. S26A</u>). To validate this approach, we confirmed agreement between estimated fold-changes testing on unsorted samples with the fold-changes estimated accounting for FACS sorting on sorted samples from the same organ (<u>fig. S26B</u>).

We constructed a KNN graph of remaining cells using similarity in the scVI embedding (k = 30 for test across gestation, k = 100 for test across tissues). We assigned cells to neighbourhoods on the KNN graph using the function *milopy.core.make_nhoods* (parameters: *prop = 0.05*). We then counted the number of cells belonging to each sample in each neighbourhood, creating a cell count matrix with rows representing neighbourhoods and columns representing samples (using the function *milopy.core.count_cells*). We assigned each neighbourhood a cell type label based on majority voting of the cells belonging to that neighbourhood. We assigned a ‘Mixed’ label if the most abundant label is present in less than 50% of cells within that neighbourhood.

#### Differential abundance across time

To test for differences in cell numbers across gestational age, we divided the sample ages into 6 equally sized bins (bin size = 2 pcw) and excluded from the cell count matrix samples from organs where less than 3 consecutive age bins were profiled (yolk sac, mesenteric lymphnode, kidney, gut). For the matrix of cell counts from samples in each organ, we modeled the cell count *c*_*n,s*_of cells from sample *s* in neighbourhood *n* as a negative binomial generalised linear model (NB-GLM):

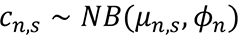

where *μ*_*n,s*_is the mean number of cells from sample *s* in neighbourhood *n* and *ϕ*_*n*_ is the dispersion parameter. We used a log-linear model to model the effect of age on cell counts:

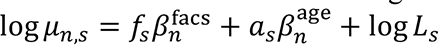

here:

– *L*_*s*_ is the sum of counts of cells of sample *s* over all the neighbourhoods.
– α_*s*_ is the age bin associated to sample *s*.
– β^age^_*n*_ is the regression coefficient encoding the effect of age on the number of cells in neighbourhood *n*, that represents the log fold-change (logFC) that can be interpreted as the per-bin linear change in neighbourhood cell abundance.
– *f*_*s*_ is the FACS sorting correction factor associated to sample *s*.
– β^facs^_*n*_ is the regression coefficient encoding the effect of CD45 enrichment on the number of cells in neighbourhood *n*.

To control for multiple testing we use the weighted BH correction as implemented by Dann et al. (*19*). In addition, we test in parallel for differential abundance associated with the library prep protocol (instead of gestational age) and exclude neighbourhoods where we detected significant differential abundance associated with library prep protocol (SpatialFDR < 0.1). We applied this stringent filtering step instead of including the library prep protocol as a covariate in the model (as described below for the test on organ enrichment) to exclude from downstream analysis false positive neighbourhoods identified in a small number of thymus samples, where we observed strong confounding between age bins and library prep protocol.

To detect markers of early-specific neighbourhoods (SpatialFDR < 0.1, logFC < 0) and/or late-specific neighbourhoods (SpatialFDR < 0.1, logFC > 0) in cell type *c* and organ *o*, we tested for differential expression between cells from organ *o* assigned to the significant neighbourhoods labelled as cell type *c* and cells belonging to all other neighbourhoods labelled as cell type *c*. We used the t-test implementation in scanpy (*scanpy.tl.rank_genes_groups*, *method = “t- test_overestim_var”*). Genes expressed in > 70% of tested cells were excluded. We considered genes as significantly over-expressed (i.e. markers) if the differential expression logFC > 1 and FDR < 0.1%. Gene set enrichment analysis was performed using the implementation of the EnrichR workflow (*84*) in the python package gseapy (https://gseapy.readthedocs.io/). The list of significantly over-expressed genes for all organs and cell types where differential expression testing was carried out can be found in Table S1 and S3.

Differential abundance between organs: We modeled the cell counts *y*_*n,s*_ for each experimental sample *s* in neighbourhood *n* by a Negative Binomial distribution:

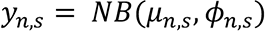

Where the expected count value *μ*_*n,s*_is given by the following log-linear model

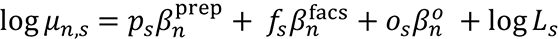

here:

– *L*_*s*_ is the sum of counts of cells of sample *s* over all the neighbourhoods.
– α_*s*_ is a binary factor indicating whether sample *s* is derived from organ *o*.
– β^*o*^_*n*_ is the regression coefficient encoding the effect of the organ on the number of cells in neighbourhood *n*, that represents the log fold-change in abundance of cells from organ *o* compared to the cells from other organs.
– *f*_*s*_ is the FACS sorting correction factor associated to sample *s*.
– β^facs^_*n*_ is the regression coefficient encoding the effect of CD45 enrichment on the number of cells in neighbourhood *n*.
– *p*_*s*_ is the binary design matrix associating sample *s* to a library prep protocol.
– β^prep^_*n*_ is the regression coefficient encoding the effect of the library prep protocol on the number of cells in neighbourhood *n*.

We estimated ²^o^_*n*_ for each *n* and *o* by fitting the NB-GLM to the count data for each neighbourhood, i.e. by estimating the dispersion *ϕ*_*n,s*_ that models the variability of cell counts in replicate samples for each neighbourhood. To control for multiple testing we use the weighted BH correction as implemented by Dann et al. (*19*).

We considered the neighbourhoods where β^o^_*n*_ > 0 and SpatialFDR < 0.01 as cell subpopulations that show organ-specific transcriptional signatures.

Having identified a subset of neighbourhoods overlapping a cell type or a subset of transcriptionally related cell types *ĉ* that were enriched in an organ *ô*, we performed differential expression (DE) analysis between these cells and cells from cell type *c* in other organs. Let *x*^*g,c,s*^_i_ be the raw gene expression counts of gene g in the *i*th cell from sample *s* and of cell type *c*. We first aggregated single-cell expression profiles into pseudo-bulk expression profiles *x̂* for each (c,s) (as recommended by (*85, 86*)):

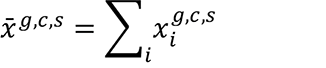

We next defined a subset of cell types and samples where we will fit the model to test for differentially expressed genes in organ *ô*. Firstly, we subsetted to the samples from donors where organ *ô* and at least 3 other organs were profiled. We then identified 3 cell types *c*^ctrl^_*j*_ ≠ *ĉ* where at least 2 pseudobulks aggregated from at least 50 cells are profiled in the selected donors, for organ *ô* and at least 3 other organs. These cell types represent populations where we don’t expect to see biological differences in expression in organ *ô*.

After sample selection, we subsetted the number of genes for DE testing selecting the top 7500 highly variable genes in *x̄*^*ĉ,s*^ using the method implemented in the R package scran. We further excluded genes where the sum of expression values across pseudobulks from either *c*^ctrl^_*j*_ or *ĉ* is equal to 0.

These steps yielded a *P*-by-*G* data matrix *X̄*, where *P* is the number of selected pseudobulks and *G* is the number of selected genes.

We modeled the mRNA counts of gene *g* in pseudobulk *p* by a NB-GLM:

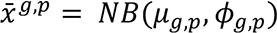

Where the expected count value *μ*_*n,s*_ is given by the following log-linear model

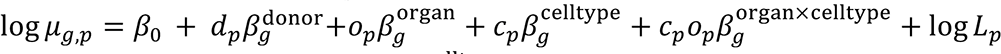

We estimated the log-fold change β^organ×celltype^_*g*_ in expression in a given cell type for organ *ô* using the quasi-likelihood method (*87*) implemented in the R package glmGamPoi (*85*).

We used the estimated logFC from the test on the control cell types to filter out genes where differential expression is driven by technical differences in tissue processing. In particular, we considered a gene to be significantly over-expressed in cell types *ĉ* and organ *ô* if it is significant in the test on *ĉ* (FDR < 10% and logFC > 1) but not in the test on control cell types (FDR > 10% and logFC > 0). We provide the full results for the differential expression analysis between organs in mature T cells and monocytes in table S2 and S4.

#### TCR analysis

Single-cell αβTCR sequencing data was mapped with cellranger-vdj (v.6.0.0). The output file *filtered_contig_annotations.csv* was used and analysed with scirpy (v.0.6.0) (*88*).

Single-cell γδTCR sequencing data was mapped with cellranger-vdj (v.4.0.0). All contigs deemed high quality were selected, and re-annotated with igblastn (v.1.17.1) against IMGT reference sequences (last downloaded: 01/08/2021) with change-o scripts (*89*), via a workflow provided in dandelion (v.0.1.10) (*90*) (https://github.com/zktuong/dandelion). The output file *hiconf_contig_igblast_db-pass.tsv* was used and analysed with scirpy (v0.6.0).

We determined productive TCR chain pairing status with *scirpy.tl.chain_qc()* function. For TCR usage principal component analysis (PCA) and clonotype analysis, cells with orphan VDJ or orphan VJ were filtered out so that each cell has at least one paired TCR. For clonotype analysis, only mature T cells were included to look at clonotype sharing. Clonotypes were determined using *scirpy.pp.ir_neighbors()* and *scirpy.tl.define_clonotypes()* functions with the CDR3 nucleotide sequence identity from both TCR chains as a metric.

Two samples from F67, F67_TH_CD137_FCAImmP7851896 and F67_TH_MAIT_FCAImmP7851897 were excluded from all downstream TCR analysis as they were sorted for specific T cell subpopulations, instead of the CD45 sorting in all other donor samples, and inclusion might result in biased TCR sampling within this donor.

#### BCR analysis

Single-cell BCR data was initially processed with cellranger-vdj (v.6.0.0). BCR contigs contained in *all_contigs.fasta* and *all_contig_annotations.csv* were then processed as follows: i) re-annotated with *igblastn* as per above; ii) re-annotated heavy chain constant region calls using *blastn* (v.2.12.0+) against curated sequences from CH1 regions of respective isotype class; and iii) heavy chain v-gene allele correction using *tigger (v1.0.0)* (*91*). Contigs were then filtered for basic quality control as described previously (*90*). Briefly, the following would lead to removal of contigs from further analysis: i) contigs were annotated with mismatched V, D, J or constant gene calls not from the same locus; ii) multiple heavy chain contigs. Exceptions to this would be when a) contigs were assessed to have identical V(D)J sequences but assigned as a different contig by cellranger-vdj (due to difference in non-V(D)J elements), b) when UMI count differences were large in which case the contig with the highest UMI count is retained, and c) if only IgM and IgD were both assigned to a cell; iii) only light chain contigs in a cell; iv) multiple light chain contigs in a cell. These were performed using dandelion (*90*) singularity container (v.0.1.10). BCR mutation frequencies were obtained using the *observedMutations* function in *shazam* (v.1.0.2) (*89*) with default settings (mutation counts for the different regions and mutation types were combined and returned as one frequency value per contig). Mutation rates per cell were averaged across contigs if multiple combinations of productive BCRs pairings were found in a single cell.

BCR clonotypes were determined with *dandelion.tl.find_clones()* function, based on the following criteria for both heavy-chain and light-chain contigs: (1) identical V and J gene usage, (2) identical junctional CDR3 amino acid length, and (3) at least 85% amino acid sequence similarity at the CDR3 junction (based on hamming distance). This strategy was chosen instead of using exact CDR3 nucleotide sequence identity to account for possible somatic hypermutations that happen within the same B cell clone.

#### Cell-cell interaction analysis

We used the CellPhoneDB v.3.0 Python package (*35*) to infer cell-cell interactions. The scRNA-seq dataset was split by organ and cell types with less than 20 cells in a given organ were filtered out. CellPhoneDB was run separately to infer cell-cell interactions in each organ, using default parameters. We used p-values from the permutation test (*pvalues.txt* output from CellPhoneDB), as well as the average expressions (‘means’) of the ligand and receptor within their corresponding cell types (*means.txt* output from CellPhoneDB). To explore cell-cell interactions between B progenitors and co-localising cell types (<u>fig. S18D</u>), we aggregated the interactions predicted between each co-localising cell type e.g. ILC3, and different subtypes of B progenitors (Pre-pro B, Pro B, Late Pro B, Large Pre B and Small Pre B cells), by averaging the means and using the minimum of the p-values. We then filtered for the ligand-receptor pairs that were significant (p-value < 0.05) across all three organs of liver, spleen and thymus, and ranked by the maximum aggregated means. Only the top 30 ligand-receptor pairs were shown in S18D.

#### Query-to-reference mapping

We mapped query data to our prenatal data embeddings using online update of the scVI models following the scArches method (*15*), as implemented in the *scvi-tools* package (*80*). The model was trained for 200 epochs and setting *weight_decay = 0*, to ensure that the latent representation of the reference cells remained exactly the same. Reference genes missing in the query were set to 0, as recommended in (*15*). To generate a joint embedding of query and reference cells, we concatenated the latent dimensions learnt for query cells to the latent dimensions used for the reference embedding and computed the KNN graph and UMAP as described above. To assess that the mapping to the developmental reference conserves biological variation while minimising technical variation in the query data, we compared query cell type labels and batch labels with clusters obtained from Leiden clustering on the learnt latent dimensions, using the Normalized Mutual Information score (see <u>fig. S25</u> for mapping of adult query data).

#### Annotation prediction using CellTypist

We used CellTypist v.0.1.5 Python package (*18*) to perform annotation prediction with logistic regression models. For prediction on cycling B cells, the rest of the mature B cells, including Immature B, Mature B, B1 and Plasma B cells were used as training dataset. Default parameters were used for model building and prediction was made without majority voting for accurate enumeration of predicted B cell subtypes within cycling B cells.

#### Comparison with human adult immune cells

Single-cell RNA-seq data from adult immune cells was generated and preprocessed as described by Conde et al. (*92*). Cell type annotations were provided by the authors. We mapped 33,694 adult lymphoid cells to the lymphoid embeddings of our developmental dataset and 13,062 adult myeloid cells to our myeloid embeddings.

In order to use cell annotations in our developmental dataset to predict adult cell types in the joint embedding, for each adult cell *c* we identified its *k* nearest prenatal cell neighbours (*N*_T_) (k = 50), and calculated the probability of assigning a label *y* to adult cell *c* as

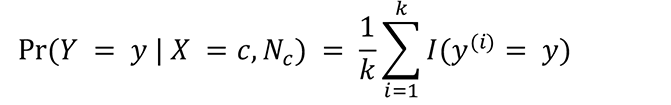

where *y*(^*i*^) is the label of the *i*th nearest neighbour and *I* is the binary indicator function. To label each cell we calculate *Ŷ*_*c*_ as follows

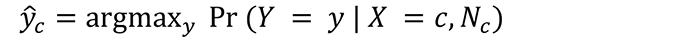

and label *c* as *Ŷ*_*c*_ if Pr(*Y* = *y* | *X* = *c*, *N*_*C*_) > 0.5, otherwise *c* is labelled as ‘low confidence’.

To quantify similarity of adult cells to prenatal cells (<u>fig. S</u>9C, <u>fig. S1</u>4C), for each adult cell *c* we calculated its similarity to prenatal cells labelled as *ŷ*_*c*_ taking the Euclidean distance in the joint embedding, weighted by a gaussian kernel following the approach described in (*15*).

#### Artificial thymic organoids scRNA-seq data analysis

Raw scRNA-seq reads were mapped with cellranger 3.0.2 with combined human reference of GRCh38.93 and mouse reference of mm10-3.1.0. Low quality cells were filtered out (minimum number of reads = 2000, minimum number of genes = 500, minimum Scrublet (*77*) doublet detection score < 0.4). Cells where the percentage of counts from human genes was < 90% were considered as mouse cells and excluded from downstream analysis. Cells were assigned to different cell lines (Kolf, Fiaj) using genotype prediction with souporcell (v.2.4.0) (*78*). We performed batch correction to minimise the differences between cells from different cell lines using scVI and clustered cells using the Leiden algorithm on the latent embedding as described above. We used CellTypist v.0.1.5 Python package (*18*) to perform annotation prediction with logistic regression prediction using the whole *in vivo* scRNA-seq developmental dataset for training. Stochastic gradient descent was used (setting *use_SGD = True*), and maximum iterations were set to 1000 in model building to reduce the run time. Predicted annotations were then aggregated using a majority voting scheme with *majority_voting = True*, *over_clustering = leiden* in CellTypist prediction to refine cell identities within Leiden clusters. For the *in vivo*-to- *in vitro* similarity analysis in <u>fig. S23E</u>, we mapped *in vitro* cells to the scVI model of lymphoid cells as described above. For each cell in the *in vitro* dataset we calculated the Euclidean distance (weighted by a gaussian kernel as described above) to the closest *in vivo* cell from each *in vivo* cell population.

### Spatial data analysis

Spatial transcriptomics data was mapped using spaceranger v.1.2.1. In parallel, we used a custom image-processing script to identify regions overlapping tissues (see data and materials availability) and retained for analysis the intersection of the tissue spots identified by this pipeline and by tissue calling by spaceranger. In two spleen slides (WSSS_F_IMMsp10864182 and WSSS_F_IMMsp10864181), we additionally excluded spots with total RNA counts < 3500, which we marked as low quality after initial QC, possibly due to a border effect. To map cell types identified by scRNA-seq in the profiled spatial transcriptomics slides, we used the cell2location method (*16*). Briefly, this consists of two steps. Firstly, for each of the profiled organs we trained a negative binomial regression model to estimate reference transcriptomic profiles for all the cell types profiled with scRNA-seq in the organ. Here we excluded very lowly expressed genes using the filtering strategy recommended by (*16*). Cell types where less than 20 cells were profiled in the organ of interest and cell types labelled as low-quality cells were excluded from the reference. For the analysis of unconventional T cell localisation in thymus (<u>fig. S21C</u>) we trained a reference adding all the prenatal Thymic Epithelial Cells profiled by Park et al. 2020 (data was downloaded from Zenodo (*93*)). Next, we estimated the abundance of cell types in the spatial transcriptomics slides using reference transcriptomic profiles of different cell types. All slides representing a given organ were analysed jointly. Cell2location requires the choice of two hyperparameters: (1) expected cell abundance (*N_cells_per_location = 30*) which was determined by counting average number of nuclei in the histology images corresponding to Visium spots; (2) regularisation strength of detection efficiency effect (*detection_alpha = 20*) was used at the low setting to account for variations in RNA detection sensitivity across different spots of Visium slides. The training was stopped after the cell2location model converged, the number of training iterations was 50,000 for thymus, liver, spleen and 30,000 for gut. All other parameters were used at default settings. Cell2location estimates the posterior distribution of cell abundance of every cell type in every spot. Posterior distribution was summarised as 5% quantile, representing the value of cell abundance that the model has high confidence in, and thus incorporating the uncertainty in the estimate into values reported in the paper and used for downstream co-localisation analysis.

To identify microenvironments of co-localising cell types, we used non-negative matrix factorisation (NMF) on the matrix of estimated cell type abundances *X* of dimensions *n* × *c*, where *n* is the total number of spots in the Visium slides and *c* is the number of cell types in the reference. We decomposed the estimated cell type abundances *X* as *X* = *WZ*^T^, where *Z* is a *n* × *d* matrix of latent factor values for each spot and *W* is a *d* × *c* matrix representing the fraction of abundance of each cell type attributed to each latent factor. Here latent factors correspond to tissue microenvironments defined by a set of co-localised cell types. We use the NMF implementation in scikit-learn (*81*), with the wrapper in the cell2location package, setting the number of factors *d =* 10. For downstream analysis, we excluded cell types where the 99% quantile of cell abundance across locations in every slide from the same organ was always below the detection threshold of 0.15. Unless otherwise specified, we consider a cell type to be part of a microenvironment if the cell type fraction is over 0.2.

For analysis of mature T cell localisation in the thymic medulla (<u>fig. S21D-E</u>), we retained factors where the sum of the cell type fractions for mature T cells (CD4+T, CD8+T, Treg, Type I Innate T, Type 3 Innate T, CD8AA) was above 0.8. We assigned spots to the inner medulla or cortico-medullary microenvironment if the factor value in the spot was above the 90% quantile of all values in the slide. To annotate histological regions in the thymus, we extracted image features from the high resolution images of H&E staining using the python package *squidpy* (*94*) (running *sq.im.calculate_image_features*, with parameters *features = “histogram”*, *spot_scale = 1*, *mask_circle = True*). We scaled and mean centered image feature matrix, and performed leiden clustering on the first 10 principal components. We manually annotated spot clusters overlapping the thymic cortex and medulla. To define the cortico-medullary junction (CMJ), we detected the spatial neighbours of each spot in medulla or cortex (using the function *squidpy.gr.spatial_neighbours*, with parameters *n_rings = 1*, *coord_type = “grid”*, *n_neighs = 6*). We then labelled a spot as CMJ if it had at least 5 neighbours (to exclude tissue borders), and if the neighbours included spots from both medulla and cortex regions. For each spot we calculated the distance to the CMJ as the Euclidean distance between the spatial coordinates of the spot and the closest spot annotated as CMJ.

### B1 functional validation experiment

Spleen was isolated from two donors, F144 (17 pcw) and F145 (15 pcw). Single cell suspension was obtained following the protocol described in the *Tissue acquisition and processing* section. Cells were then cryopreserved with 90% FBS and 10% DMSO (Sigma-Aldrich). On the day of the ELISpot experiment, cells were thawed and stained with anti-CD3, anti-CD20, anti-CD43, anti-CD27, anti-CD38, anti-CCR10 antibodies and DAPI together with control peripheral blood mononuclear cells (PBMC; Stemcell Technologies). Mature B cells were gated from single cells, then DAPI^−^CD3^−^CD20^+^. The top 1% of cells expressing the highest level of CD38 were gated out to exclude possible plasma cell contamination (which should also be CD20 low and therefore not gated in). The rest of the mature B cells were then sorted into 4 fractions — CCR10^hi^, CCR10^lo^CD27^+^CD43^+^, CCR10^lo^CD27^−^CD43^+^, CCR10^lo^CD27^−^CD43^−^. CD27 and CD43 gatings were chosen based on fluorescence minus one (FMO) controls. The cells were sorted into RPMI supplemented with 10% FBS, penicillin/streptomycin (Gibco) and glutamax (Thermo Fisher Scientific).

ELISpot experiment was performed with Human IgM ELISpot^BASIC^ kit (ALP) from Mabtech AB. 7000-8000 of sorted cells were added into ELISpot plate pre-coated with anti-IgM antibody following manufacturer’s instructions and incubated in a 37°C humidified incubator with 5% CO2 for 22 hours. The plate was then washed and incubated with biotinylated anti-IgM for 2 hours at room temperature, followed by 1-hour incubation of streptavidin-ALP. The coloured spots were developed with 15-min incubation of BCIP/NBT substrate solution (Thermo Fisher Scientific). Five rounds of washing were performed between each step of incubation as per manufacturer’s instructions. After the coloured spots appeared clearly, the reaction was then stopped by rinsing under running tap water for 5 min. Spots were counted with the AID ELISpot reader and iSpot software version 4.

### Data and code availability

Raw sequencing data for newly generated sequencing libraries have been deposited in ArrayExpress (scRNA-seq libraries: accession number E-MTAB-11343; scVDJ-seq libraries: accession number TBD; Visium 10X libraries: accession number E-MTAB-11341). Processed data objects are available for online visualisation and download in AnnData format (*74*) at https://developmentcellatlas.cellgeni.sanger.ac.uk/fetal-immune, as well as trained scVI models for query to reference mapping and trained Celltypist models for cell annotation. All code scripts and notebooks for analysis presented in the manuscript are available at https://github.com/Teichlab/Pan_fetal_immune.

## Notes

https://developmentcellatlas.cellgeni.sanger.ac.uk/fetal-immune

### Competing Interest Statement

In the past 3 years, Sarah A. Teichmann has consulted for Genentech and Roche and sits on Scientific Advisory Boards for Qiagen, Foresite Labs, Biogen, and GlaxoSmithKline and is a co-founder and equity holder of Transition Bio.

