## Supplementary Figures for "Mapping the developing human immune system across organs"

**This PDF file includes:**

Figs. S1 to S26
Tables S1 to S4

### Supplementary Figures

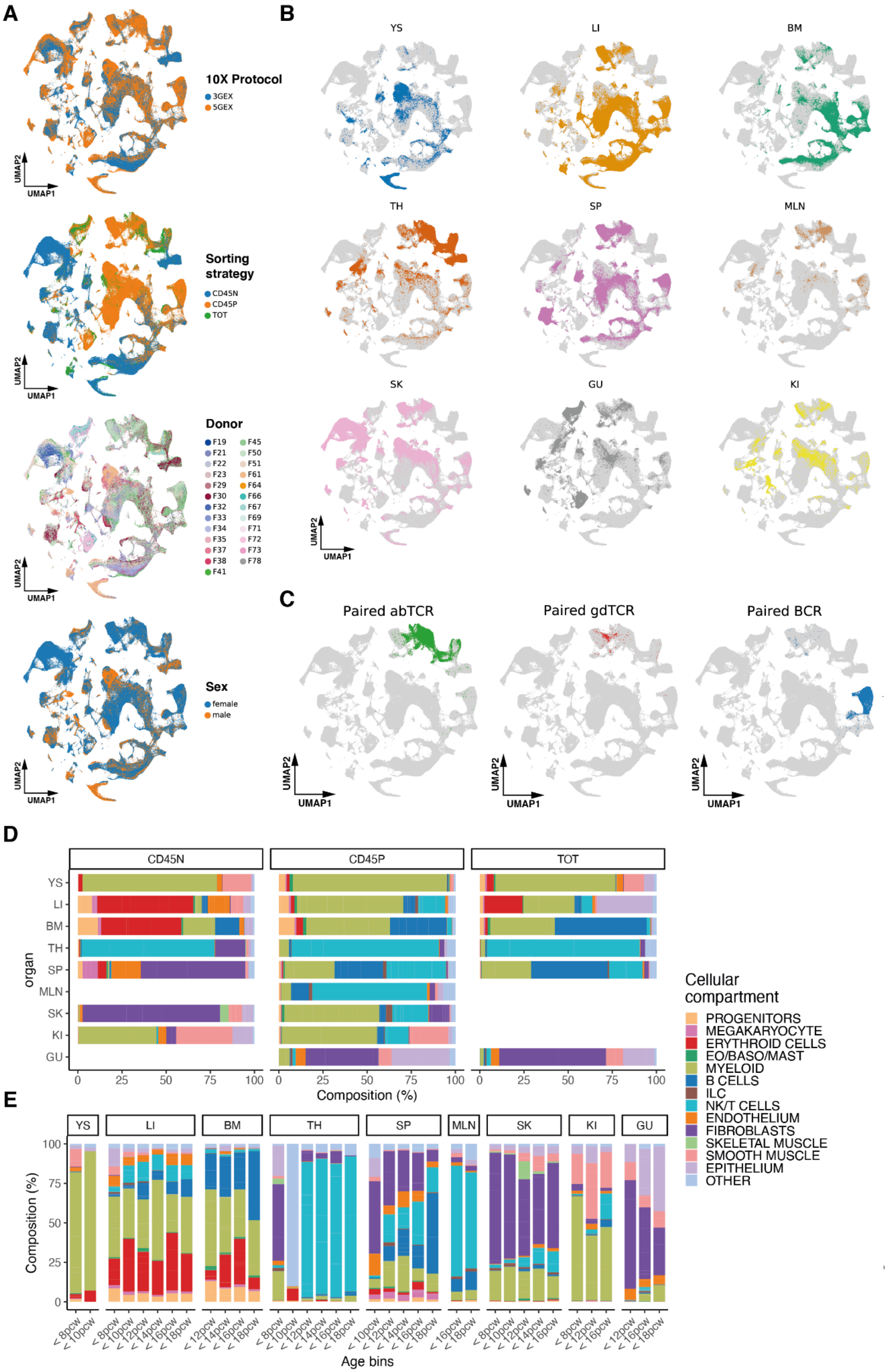

A
10X Protocol
3GEX
5GEX
Sorting strategy
CD45N
CD45P
TOT
Donor
F19 F21 F22 F23 F29 F30 F32 F33 F34 F35 F37 F38 F41
F45 F50 F51 F61 F64 F66 F67 F69 F71 F72 F73 F78
Sex
female
male
UMAP1
UMAP2
B
YS LI BM TH SP MLN SK GU KI
C
Paired abTCR
Paired gdTCR
Paired BCR
D
CD45N
CD45P
TOT
organ
YS LI BM TH SP MLN SK KI GU
Composition (%)
Cellular compartment
PROGENITORS
MEGAKARYOCYTE
ERYTHROID CELLS
EO/BASO/MAST
MYELOID
B CELLS
ILC
NK/T CELLS
ENDOTHELIUM
FIBROBLASTS
SKELETAL MUSCLE
SMOOTH MUSCLE
EPITHELIUM
OTHER
E
Composition (%)
Age bins

**fig. S1: Characterisation of cross-organ developmental scRNA-seq atlas.** (**A**) Sample characteristics in integrated atlas. UMAP embeddings (as in Fig. 1C) of scRNA-seq profiles coloured by (top to bottom): 10x chemistry protocol, FACS sorting protocol (CD45P: $CD45^+$; CD45N: $CD45^-$; TOT: unsorted), donor ID, sex of donor. (**B**) Distribution of cells from different organs in integrated atlas. UMAP embeddings (as in Fig. 1C) of scRNA-seq profiles, highlighting cells from each organ. (**C**) UMAP embeddings (as in Fig. 1C) of scRNA-seq profiles, highlighting cells for which paired αβTCR, γδTCR or BCR sequences were detected. (**D**) Percentage of cells of each broad type in each organ, stratified by FACS sorting protocol. (**E**) Percentage of cells of each broad type in each gestational age group, stratified by organ. (YS: yolk sac; LI: liver; BM: bone marrow; TH: thymus; SP: spleen; SK: skin; GU: gut; KI: kidney).

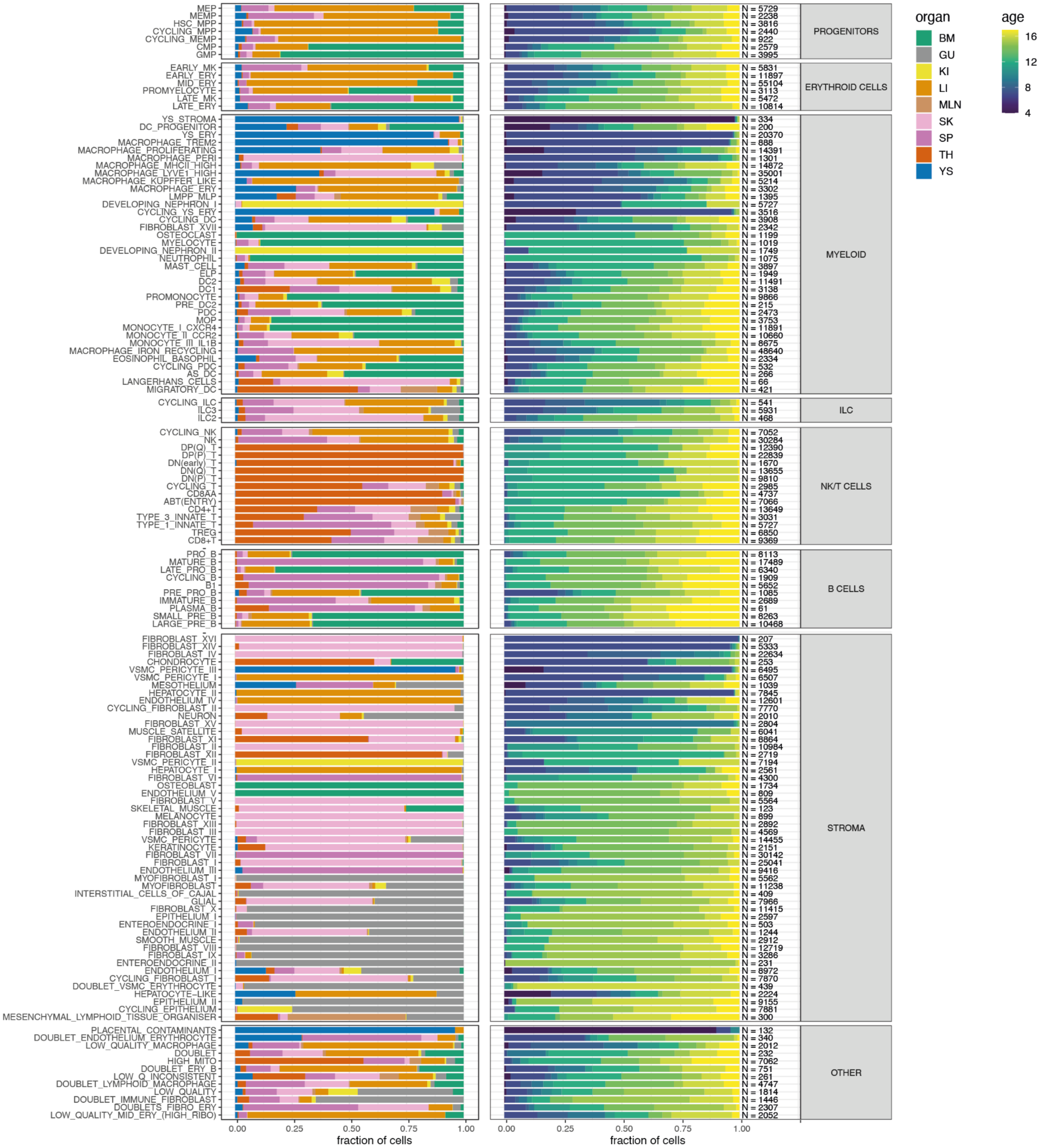


**fig. S2: Distribution across organs (left) and gestational age (pcw, right) of annotated cell populations.** Cell populations are grouped according to broad population annotations. The category ‘Other’ denotes clusters annotated as low-quality cells. N indicates the total number of cells across the dataset for each annotation.

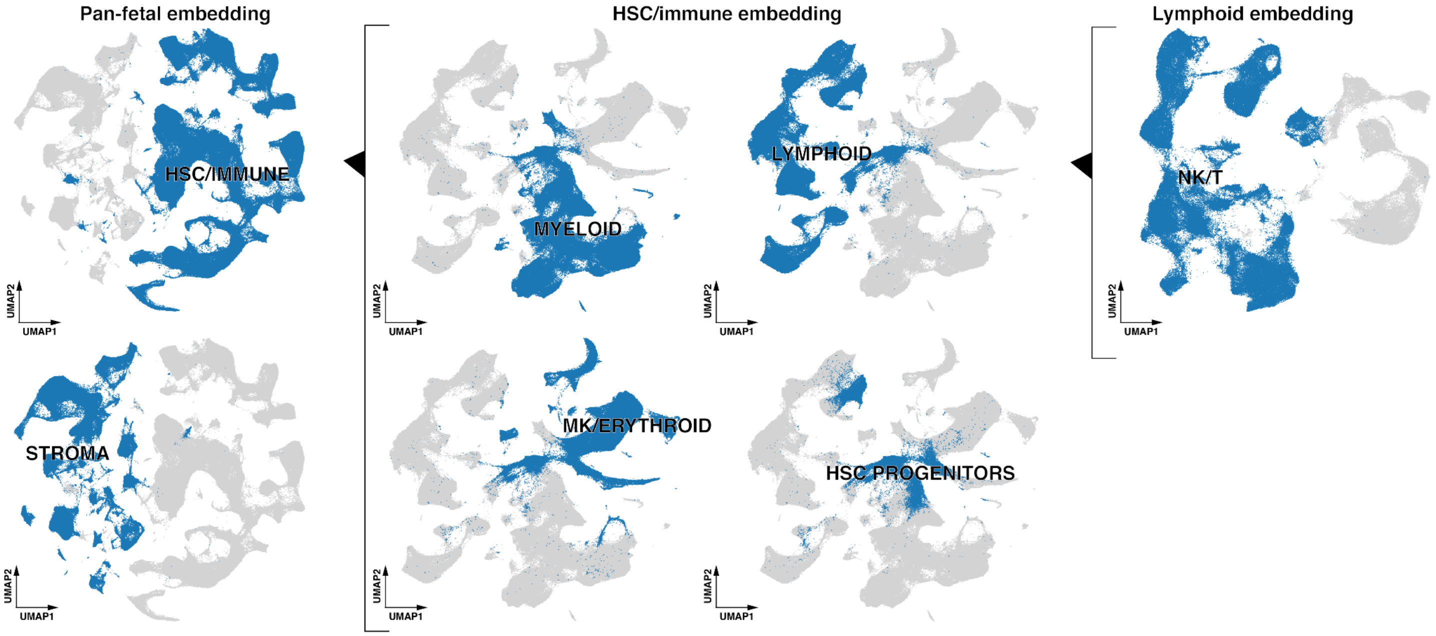


**fig. S3: Overview of the hierarchical subsetting strategy used for annotation of fine immune subtypes.** In each embedding, the cells that make up the data views in fig. S4 are highlighted in blue.

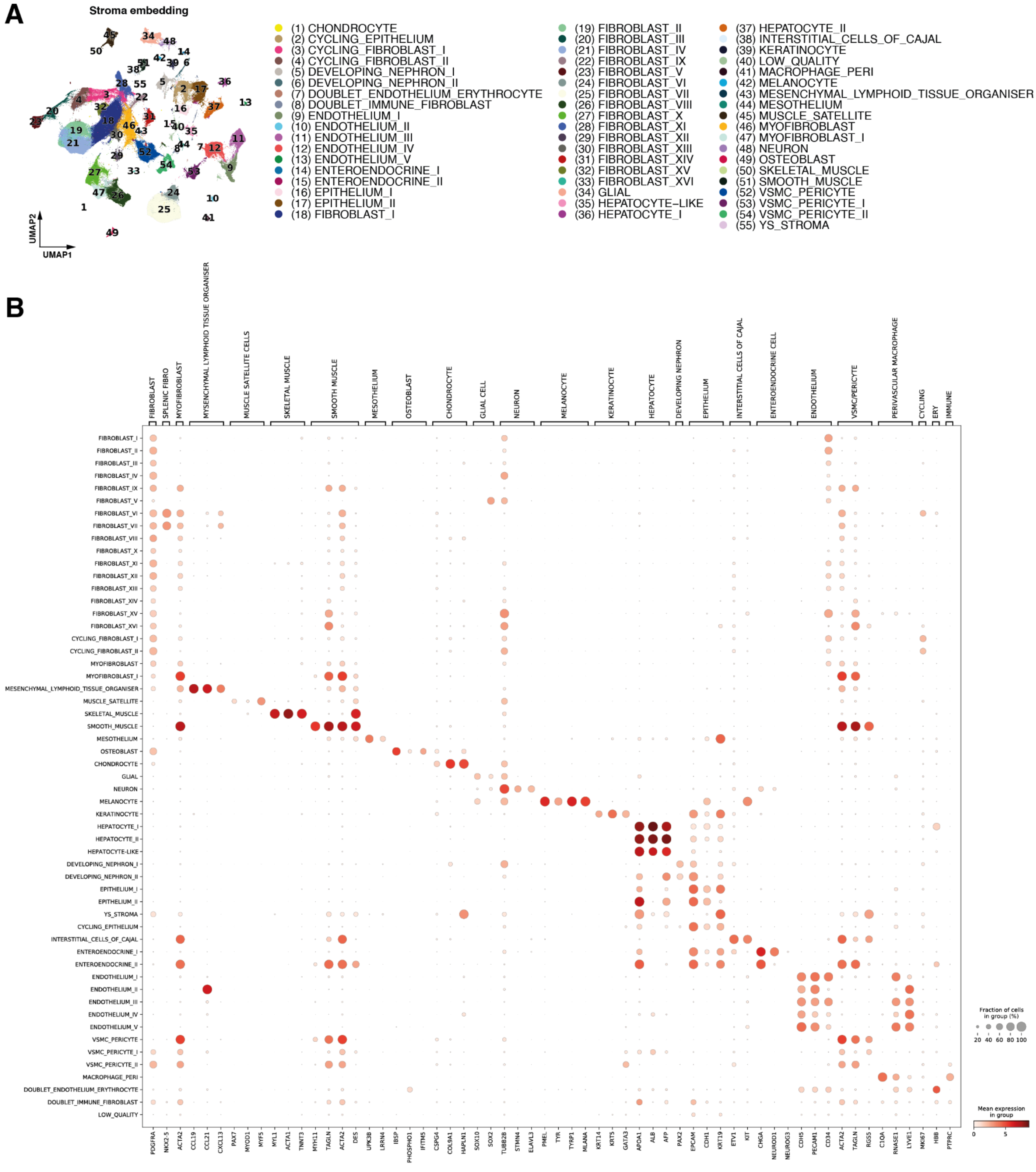

A
Stroma embedding
UMAP1
UMAP2
(1) CHONDROCYTE
(2) CYCLING_EPITHELIUM
(3) CYCLING_FIBROBLAST_I
(4) CYCLING_FIBROBLAST_II
(5) DEVELOPING_NEPHRON_I
(6) DEVELOPING_NEPHRON_II
(7) DOUBLET_ENDOTHELIUM_ERYTHROCYTE
(8) DOUBLET_IMMUNE_FIBROBLAST
(9) ENDOTHELIUM_I
(10) ENDOTHELIUM_II
(11) ENDOTHELIUM_III
(12) ENDOTHELIUM_IV
(13) ENDOTHELIUM_V
(14) ENTEROENDOCRINE_I
(15) ENTEROENDOCRINE_II
(16) EPITHELIUM_I
(17) EPITHELIUM_II
(18) FIBROBLAST_I
(19) FIBROBLAST_II
(20) FIBROBLAST_III
(21) FIBROBLAST_IV
(22) FIBROBLAST_IX
(23) FIBROBLAST_V
(24) FIBROBLAST_VI
(25) FIBROBLAST_VII
(26) FIBROBLAST_VIII
(27) FIBROBLAST_X
(28) FIBROBLAST_XI
(29) FIBROBLAST_XII
(30) FIBROBLAST_XIII
(31) FIBROBLAST_XIV
(32) FIBROBLAST_XV
(33) FIBROBLAST_XVI
(34) GLIAL
(35) HEPATOCYTE−LIKE
(36) HEPATOCYTE_I
(37) HEPATOCYTE_II
(38) INTERSTITIAL_CELLS_OF_CAJAL
(39) KERATINOCYTE
(40) LOW_QUALITY
(41) MACROPHAGE_PERI
(42) MELANOCYTE
(43) MESENCHYMAL_LYMPHOID_TISSUE_ORGANISER
(44) MESOTHELIUM
(45) MUSCLE_SATELLITE
(46) MYOFIBROBLAST
(47) MYOFIBROBLAST_I
(48) NEURON
(49) OSTEOBLAST
(50) SKELETAL_MUSCLE
(51) SMOOTH_MUSCLE
(52) VSMC_PERICYTE
(53) VSMC_PERICYTE_I
(54) VSMC_PERICYTE_II
(55) YS_STROMA
B
FIBROBLAST
SPLENIC FIBRO
MYOFIBROBLAST
MYSENCHYMAL LYMPHOID TISSUE ORGANISER
MUSCLE SATELLITE CELLS
SKELETAL MUSCLE
SMOOTH MUSCLE
MESOTHELIUM
OSTEOBLAST
CHONDROCYTE
GLIAL CELL
NEURON
MELANOCYTE
KERATINOCYTE
HEPATOCYTE
DEVELOPING NEPHRON
EPITHELIUM
INTERSTITIAL CELLS OF CAJAL
ENTEROENDOCRINE CELL
ENDOTHELIUM
VSMC/PERICYTE
PERIVASCULAR MACROPHAGE
CYCLING
ERY
IMMUNE
FIBROBLAST_I
FIBROBLAST_II
FIBROBLAST_III
FIBROBLAST_IV
FIBROBLAST_IX
FIBROBLAST_V
FIBROBLAST_VI
FIBROBLAST_VII
FIBROBLAST_VIII
FIBROBLAST_X
FIBROBLAST_XI
FIBROBLAST_XII
FIBROBLAST_XIII
FIBROBLAST_XIV
FIBROBLAST_XV
FIBROBLAST_XVI
CYCLING_FIBROBLAST_I
CYCLING_FIBROBLAST_II
MYOFIBROBLAST
MYOFIBROBLAST_I
MESENCHYMAL_LYMPHOID_TISSUE_ORGANISER
MUSCLE_SATELLITE
SKELETAL_MUSCLE
SMOOTH_MUSCLE
MESOTHELIUM
OSTEOBLAST
CHONDROCYTE
GLIAL
NEURON
MELANOCYTE
KERATINOCYTE
HEPATOCYTE_I
HEPATOCYTE_II
HEPATOCYTE-LIKE
DEVELOPING_NEPHRON_I
DEVELOPING_NEPHRON_II
EPITHELIUM_I
EPITHELIUM_II
YS_STROMA
CYCLING_EPITHELIUM
INTERSTITIAL_CELLS_OF_CAJAL
ENTEROENDOCRINE_I
ENTEROENDOCRINE_II
ENDOTHELIUM_I
ENDOTHELIUM_II
ENDOTHELIUM_III
ENDOTHELIUM_IV
ENDOTHELIUM_V
VSMC_PERICYTE
VSMC_PERICYTE_I
VSMC_PERICYTE_II
MACROPHAGE_PERI
DOUBLET_ENDOTHELIUM_ERYTHROCYTE
DOUBLET_IMMUNE_FIBROBLAST
LOW_QUALITY
PDGFRA
NKX2-5
ACTA2
CCL19
CCL21
CXCL13
PAX7
MYOD1
MYF5
MYL1
ACTA1
TNNT3
MYH11
TAGLN
ACTA2
DES
UPK3B
LRRN4
IBSP
PHOSPHO1
IFITM5
CSPG4
COL9A1
HAPLN1
SOX10
SOX2
TUBB2B
STMN4
ELAVL3
PMEL
TYR
TYRP1
MLANA
KRT14
KRT5
GATA3
APOA1
ALB
AFP
PAX2
EPCAM
CDH1
KRT19
ETV1
KIT
CHGA
NEUROD1
NEUROG3
CDH5
PECAM1
CD34
ACTA2
TAGLN
RGS5
C1QA
RNASE1
LYVE1
MKI67
HBB
PTPRC
Fraction of cells in group (%)
20 40 60 80 100
Mean expression in group
0 5

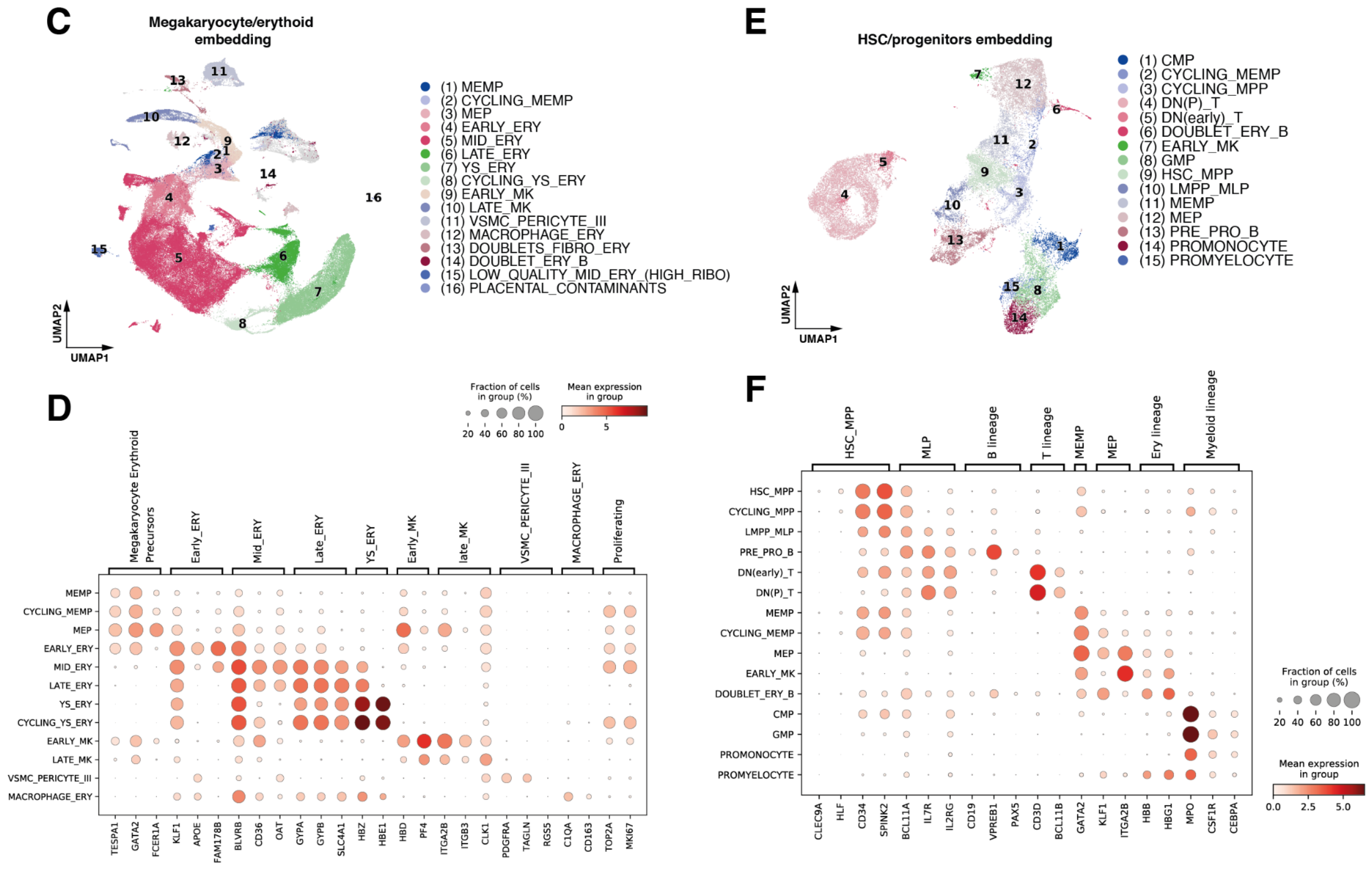
C
Megakaryocyte/erythoid embedding
(1) MEMP
(2) CYCLING_MEMP
(3) MEP
(4) EARLY_ERY
(5) MID_ERY
(6) LATE_ERY
(7) YS_ERY
(8) CYCLING_YS_ERY
(9) EARLY_MK
(10) LATE_MK
(11) VSMC_PERICYTE_III
(12) MACROPHAGE_ERY
(13) DOUBLETS_FIBRO_ERY
(14) DOUBLET_ERY_B
(15) LOW_QUALITY_MID_ERY_(HIGH_RIBO)
(16) PLACENTAL_CONTAMINANTS
UMAP1
UMAP2
E
HSC/progenitors embedding
(1) CMP
(2) CYCLING_MEMP
(3) CYCLING_MPP
(4) DN(P)_T
(5) DN(early)_T
(6) DOUBLET_ERY_B
(7) EARLY_MK
(8) GMP
(9) HSC_MPP
(10) LMPP_MLP
(11) MEMP
(12) MEP
(13) PRE_PRO_B
(14) PROMONOCYTE
(15) PROMYELOCYTE
D
Fraction of cells in group (%)
20 40 60 80 100
Mean expression in group
0 5
Megakaryocyte Erythroid Precursors
Early_ERY
Mid_ERY
Late_ERY
YS_ERY
Early_MK
late_MK
VSMC_PERICYTE_III
MACROPHAGE_ERY
Proliferating
MEMP
CYCLING_MEMP
MEP
EARLY_ERY
MID_ERY
LATE_ERY
YS_ERY
CYCLING_YS_ERY
EARLY_MK
LATE_MK
VSMC_PERICYTE_III
MACROPHAGE_ERY
TESPA1 GATA2 FCER1A KLF1 APOE FAM178B BLVRB CD36 OAT GYPA GYPB SLC4A1 HBZ HBE1 HBD PF4 ITGA2B ITGB3 CLK1 PDGFRA TAGLN RGS5 C1QA CD163 TOP2A MKI67
F
HSC_MPP
MLP
B lineage
T lineage
MEMP
MEP
Ery lineage
Myeloid lineage
HSC_MPP
CYCLING_MPP
LMPP_MLP
PRE_PRO_B
DN(early)_T
DN(P)_T
MEMP
CYCLING_MEMP
MEP
EARLY_MK
DOUBLET_ERY_B
CMP
GMP
PROMONOCYTE
PROMYELOCYTE
CLEC9A HLF CD34 SPINK2 BCL11A IL7R IL2RG CD19 VPREB1 PAX5 CD3D BCL11B GATA2 KLF1 ITGA2B HBB HBG1 MPO CSF1R CEBPA
Fraction of cells in group (%)
20 40 60 80 100
Mean expression in group
0.0 2.5 5.0

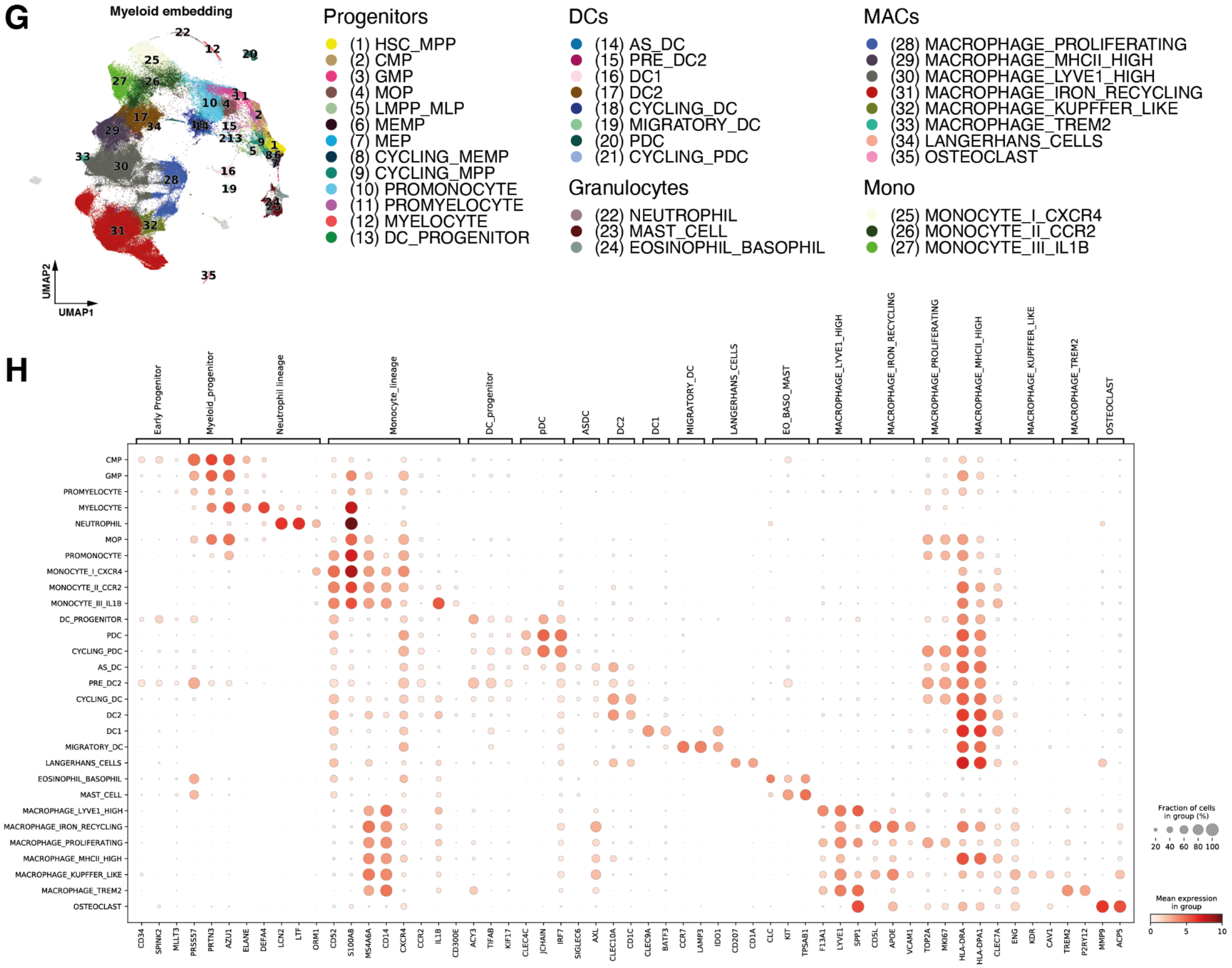
G
Myeloid embedding
UMAP1
UMAP2
Progenitors
(1) HSC_MPP
(2) CMP
(3) GMP
(4) MOP
(5) LMPP_MLP
(6) MEMP
(7) MEP
(8) CYCLING_MEMP
(9) CYCLING_MPP
(10) PROMONOCYTE
(11) PROMYELOCYTE
(12) MYELOCYTE
(13) DC_PROGENITOR
DCs
(14) AS_DC
(15) PRE_DC2
(16) DC1
(17) DC2
(18) CYCLING_DC
(19) MIGRATORY_DC
(20) PDC
(21) CYCLING_PDC
Granulocytes
(22) NEUTROPHIL
(23) MAST_CELL
(24) EOSINOPHIL_BASOPHIL
MACs
(28) MACROPHAGE_PROLIFERATING
(29) MACROPHAGE_MHCII_HIGH
(30) MACROPHAGE_LYVE1_HIGH
(31) MACROPHAGE_IRON_RECYCLING
(32) MACROPHAGE_KUPFFER_LIKE
(33) MACROPHAGE_TREM2
(34) LANGERHANS_CELLS
(35) OSTEOCLAST
Mono
(25) MONOCYTE_I_CXCR4
(26) MONOCYTE_II_CCR2
(27) MONOCYTE_III_IL1B
H
Early Progenitor
Myeloid_progenitor
Neutrophil lineage
Monocyte_lineage
DC_progenitor
pDC
ASDC
DC2
DC1
MIGRATORY_DC
LANGERHANS_CELLS
EO_BASO_MAST
MACROPHAGE_LYVE1_HIGH
MACROPHAGE_IRON_RECYCLING
MACROPHAGE_PROLIFERATING
MACROPHAGE_MHCII_HIGH
MACROPHAGE_KUPFFER_LIKE
MACROPHAGE_TREM2
OSTEOCLAST
CMP
GMP
PROMYELOCYTE
MYELOCYTE
NEUTROPHIL
MOP
PROMONOCYTE
MONOCYTE_I_CXCR4
MONOCYTE_II_CCR2
MONOCYTE_III_IL1B
DC_PROGENITOR
PDC
CYCLING_PDC
AS_DC
PRE_DC2
CYCLING_DC
DC2
DC1
MIGRATORY_DC
LANGERHANS_CELLS
EOSINOPHIL_BASOPHIL
MAST_CELL
MACROPHAGE_LYVE1_HIGH
MACROPHAGE_IRON_RECYCLING
MACROPHAGE_PROLIFERATING
MACROPHAGE_MHCII_HIGH
MACROPHAGE_KUPFFER_LIKE
MACROPHAGE_TREM2
OSTEOCLAST
CD34 SPINK2 MLLT3 PRSS57 PRTN3 AZU1 ELANE DEFA4 LCN2 LTF ORM1 CD52 S100A8 MS4A6A CD14 CXCR4 CCR2 IL1B CD300E ACY3 TIFAB KIF17 CLEC4C JCHAIN IRF7 SIGLEC6 AXL CLEC10A CD1C CLEC9A BATF3 CCR7 LAMP3 IDO1 CD207 CD1A CLC KIT TPSAB1 F13A1 LYVE1 SPP1 CD5L APOE VCAM1 TOP2A MKI67 HLA-DRA HLA-DPA1 CLEC7A ENG KDR CAV1 TREM2 P2RY12 MMP9 ACP5
Fraction of cells in group (%)
20 40 60 80 100
Mean expression in group
0 5 10

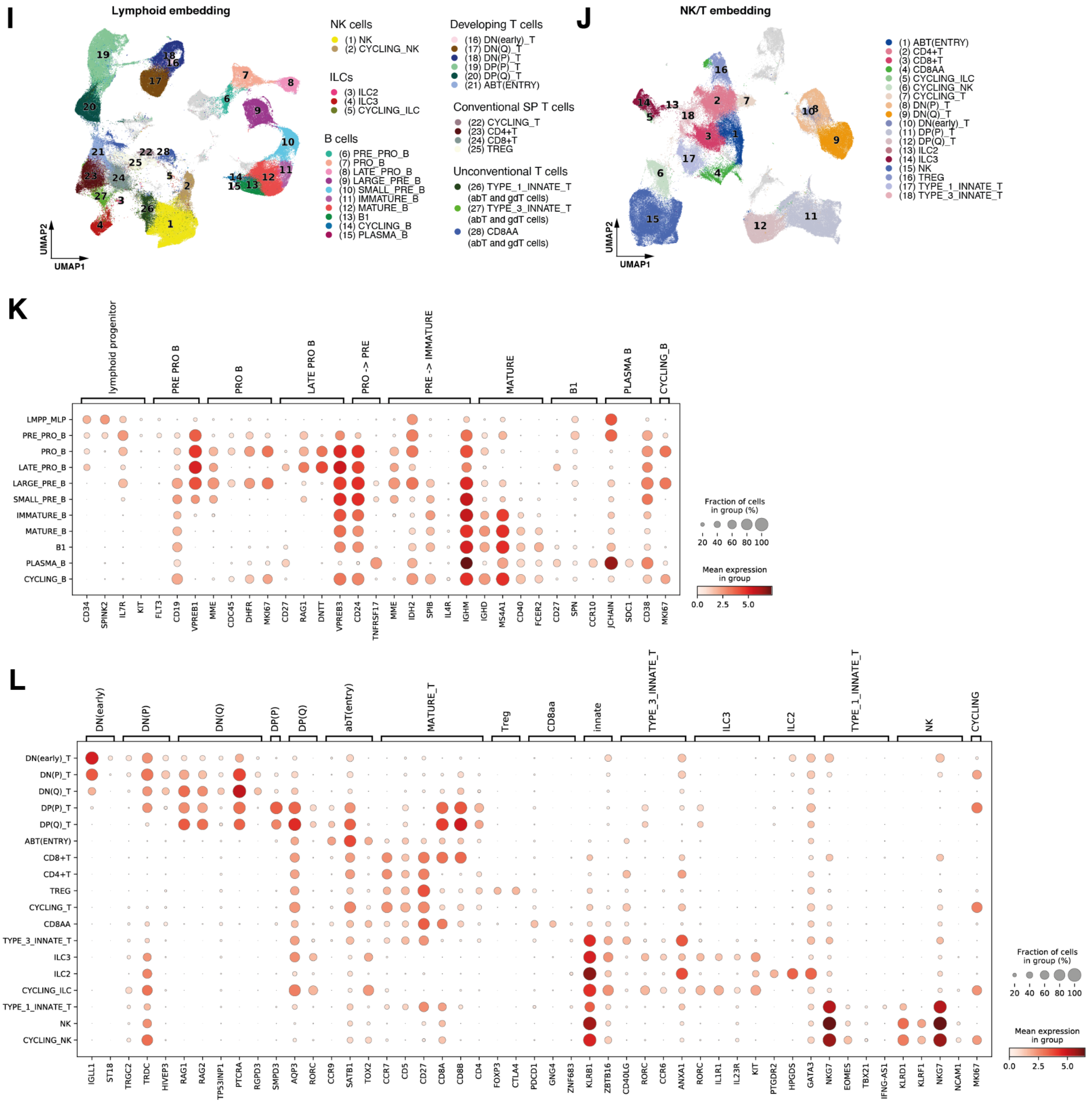


**fig. S4: Cross-tissue annotation of hierarchical subsets of scRNA-seq integrated dataset.** For each subset embedding generated through scVI, we show UMAP embeddings of cells coloured by annotated cell populations and dot plots of mean expression (log-normalised counts, dot colour) and fraction of expressing cells (dot size) of marker genes (columns) used for cell population annotation (rows). (**A-B**) Annotation of stromal cells. (**C-D**) Annotation of megakaryocyte and erythroid cells (cells in grey are progenitors annotated through embedding shown in (E)). (**E-F**) Annotation of haematopoietic and immune cell progenitors. (**G-H**) Annotation of myeloid cells (cells in grey are progenitors annotated through embedding shown in (E) or low-quality clusters) (**I-L**) Annotation of lymphoid cells (cells in grey are progenitors annotated through embedding shown in (E) or low-quality clusters). The embedding of all lymphoid cells is shown in (I), the embedding used for annotation of NK/T cells is visualised in (L). The dot plot for annotation of B cells is shown in (M), the dot plot for annotation of T cells is shown in (N).

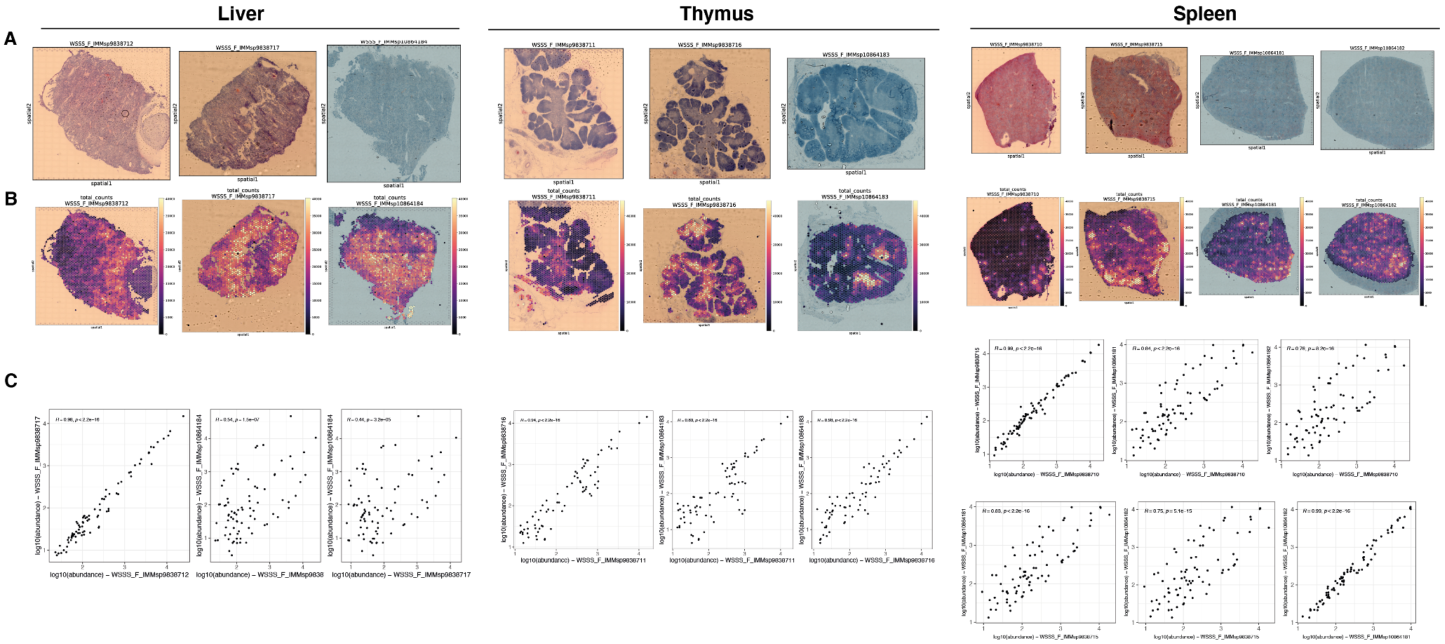


**fig. S5: Quality control metrics for Visium 10x data and cell type mapping with cell2location** (**A**) H&E staining of tissue slides processed for spatial transcriptomics with Visium 10x protocol. (**B**) Total RNA counts in analysed tissue spots. (**C**) Analysis of robustness of cell type mapping with cell2location: for mapping on each organ we correlate the total abundance (in $\log_{10}$ scale) of each cell type (points) in different tissue slides from the same organ (biological replicates). The Pearson's correlation coefficient and p-value for permutation test are reported.

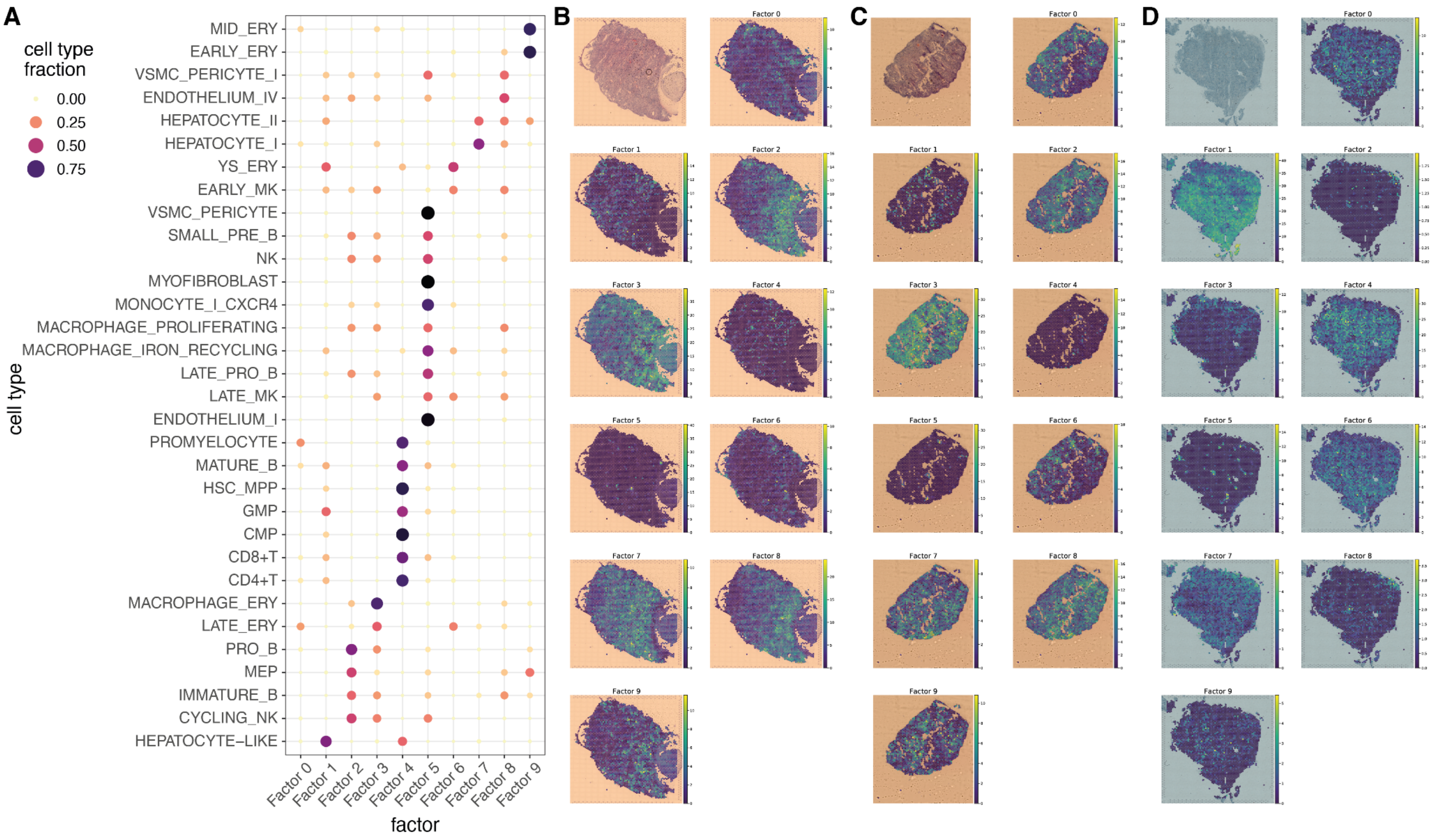


**fig. S6: Cell type spatial microenvironments in the fetal liver detected by non-negative matrix factorization on spatial cell type abundances.** (**A**) Dot plot of cell type contributions to latent factors (microenvironments) identified with non-negative matrix factorisation (NMF) of spatial cell type abundances estimated with cell2location. The colour and the size of the dots represent the relative fraction of the cell population assigned to the factor. We exclude cell types where the value for the 99% quantile of cell abundance in all the slides from the same organ is always below the detection threshold of 0.15. (**B-D**) Spatial locations of microenvironments on liver slides, with the colour representing the weighted contribution of each microenvironment to each spot. The image of the H&E staining for the slide is shown for reference.

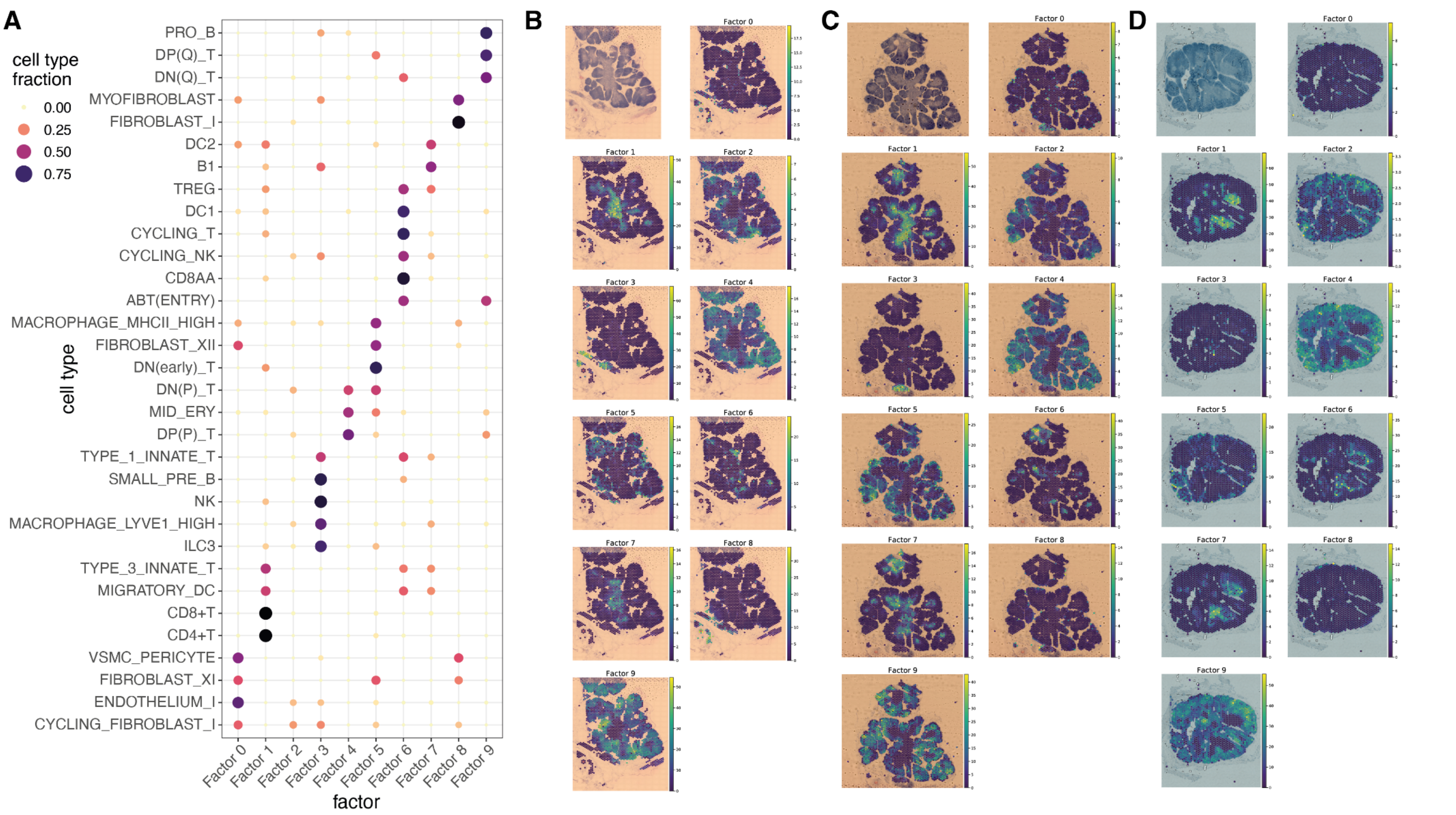


**fig. S7: Cell type spatial microenvironments in the fetal thymus detected by non-negative matrix factorization on spatial abundances.** (**A**) Dot plot of cell type contributions to latent factors (microenvironments) identified with non-negative matrix factorisation (NMF) of spatial cell type abundances estimated with cell2location. The colour and the size of the dots represent the relative fraction of the cell population assigned to the factor. We exclude cell types where the value for the 99% quantile of cell abundance in all the slides from the same organ is always below the detection threshold of 0.15. (**B-D**) Spatial locations of microenvironments on thymus slides, with the colour representing the weighted contribution of each microenvironment to each spot. The image of the H&E staining for the slide is shown for reference.

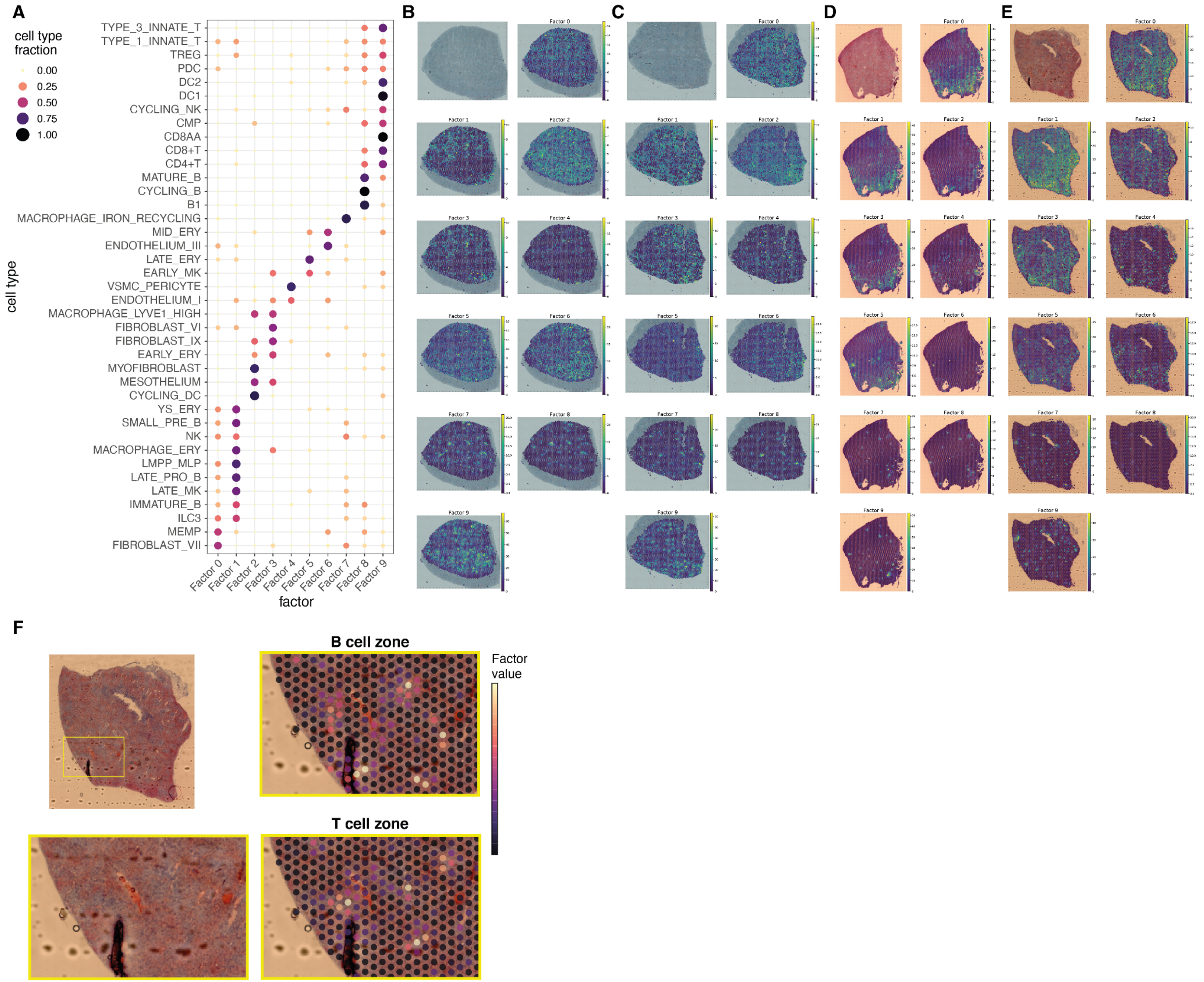


**fig. S8: Cell type spatial microenvironments in the fetal spleen detected by non-negative matrix factorization on spatial abundances.** (**A**) Dot plot of cell type contributions to latent factors (microenvironments) identified with non-negative matrix factorisation (NMF) of spatial cell type abundances estimated with cell2location. The colour and the size of the dots represent the relative fraction of the cell population assigned to the factor. We exclude cell types where the value for the 99% quantile of cell abundance in all the slides from the same organ is always below the detection threshold of 0.15. (**B-E**) Spatial locations of microenvironments on spleen slides, with the colour representing the weighted contribution of each microenvironment to each spot. The image of the H&E staining for the slide is shown for reference. (**F**) Zoomed-in views of fetal spleen tissue slide shown in Fig. 1D, showing weighted microenvironment contribution (factor values) of lymphoid aggregates B cell zone microenvironment (Factor 8) and T cell zone microenvironment (Factor 9). These exemplify how the B and T cell zones were proximal to each other but not completely overlapping.

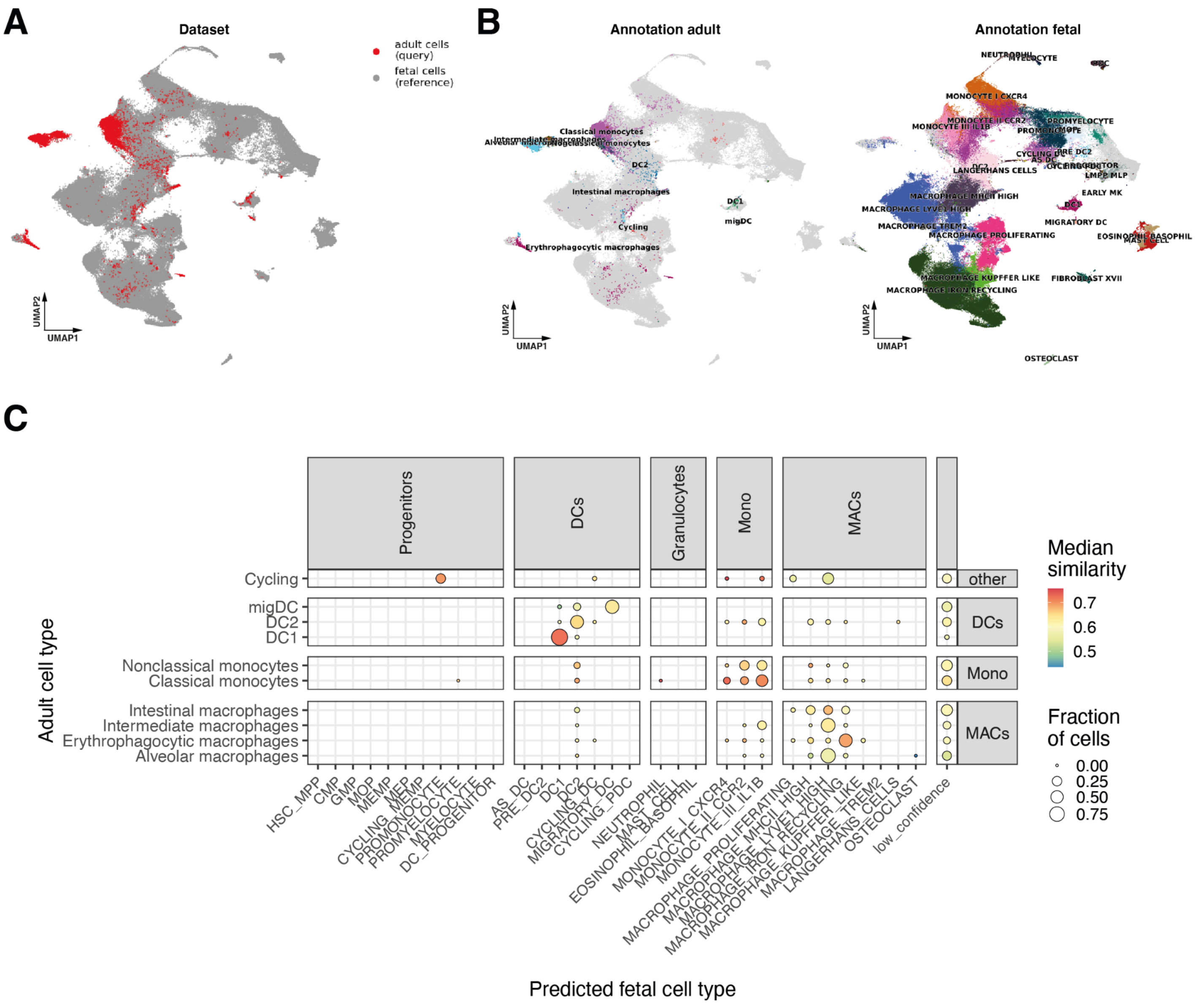


**fig. S9: Mapping of adult myeloid cells to prenatal reference with transfer learning**. (**A**) UMAP embeddings of mapping of adult myeloid cells (13,062) to developmental myeloid reference (218,758 cells) using scArches on scVI model. Points are coloured by the dataset of origin. (**B**) UMAP embedding as in A, points are coloured by cell population annotation label, for adult cells (left) and developmental cells (right). (**C**) Correspondence between developmental and adult myeloid transcriptional phenotypes estimated by label transfer after mapping with scArches. The dot size is proportional to the fraction of cells in the adult population (y-axis) with a given predicted prenatal cell population label (x-axis). The dot colour denotes the median similarity of the adult cells to prenatal cells in the common embedding (see Methods). Adult cells where less than 50% of prenatal neighbours have a uniform annotation are labelled as 'low confidence'.

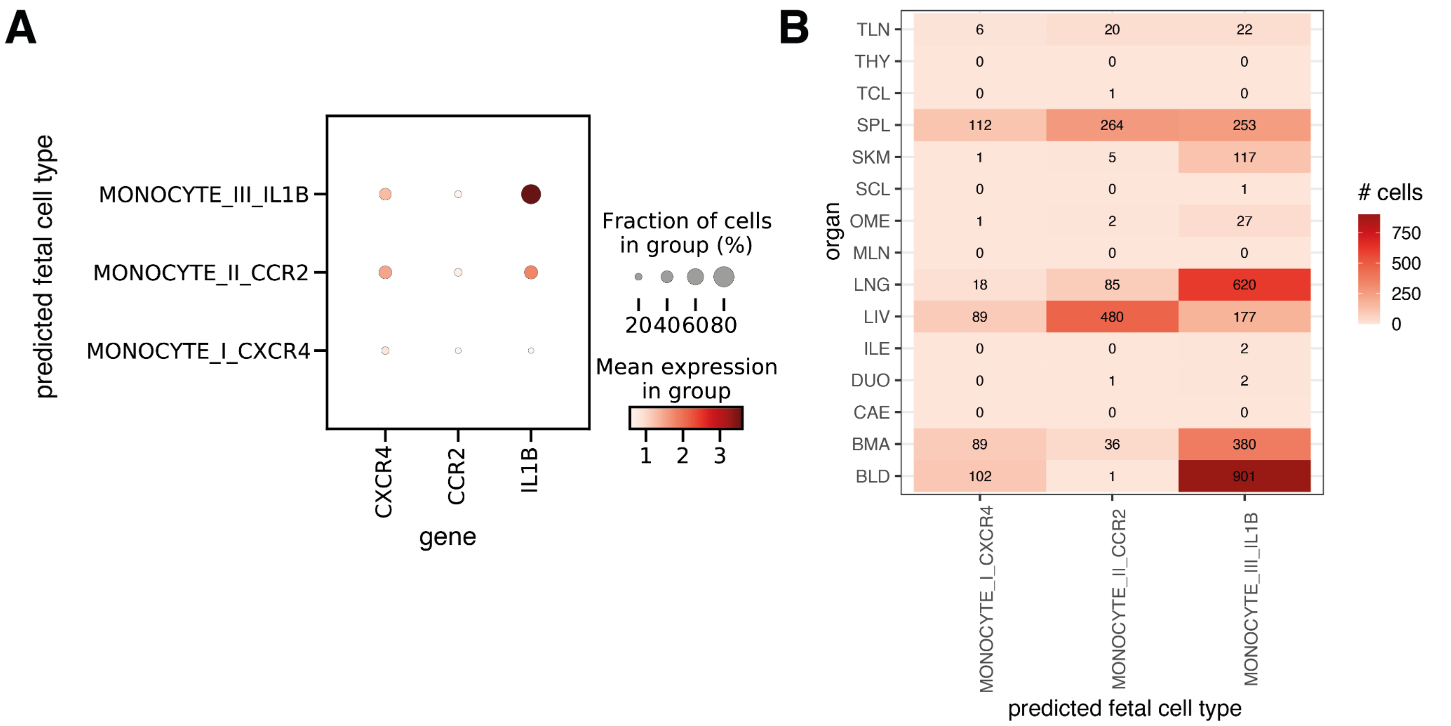


**fig. S10: Prenatal-adult comparison in monocytes.** (**A**) Dot plot of expression of monocyte subtype markers in adult cells aligned to prenatal monocyte subtypes. (**B**) Heat map of distribution across organs of adult cells aligned to prenatal monocyte subtypes.

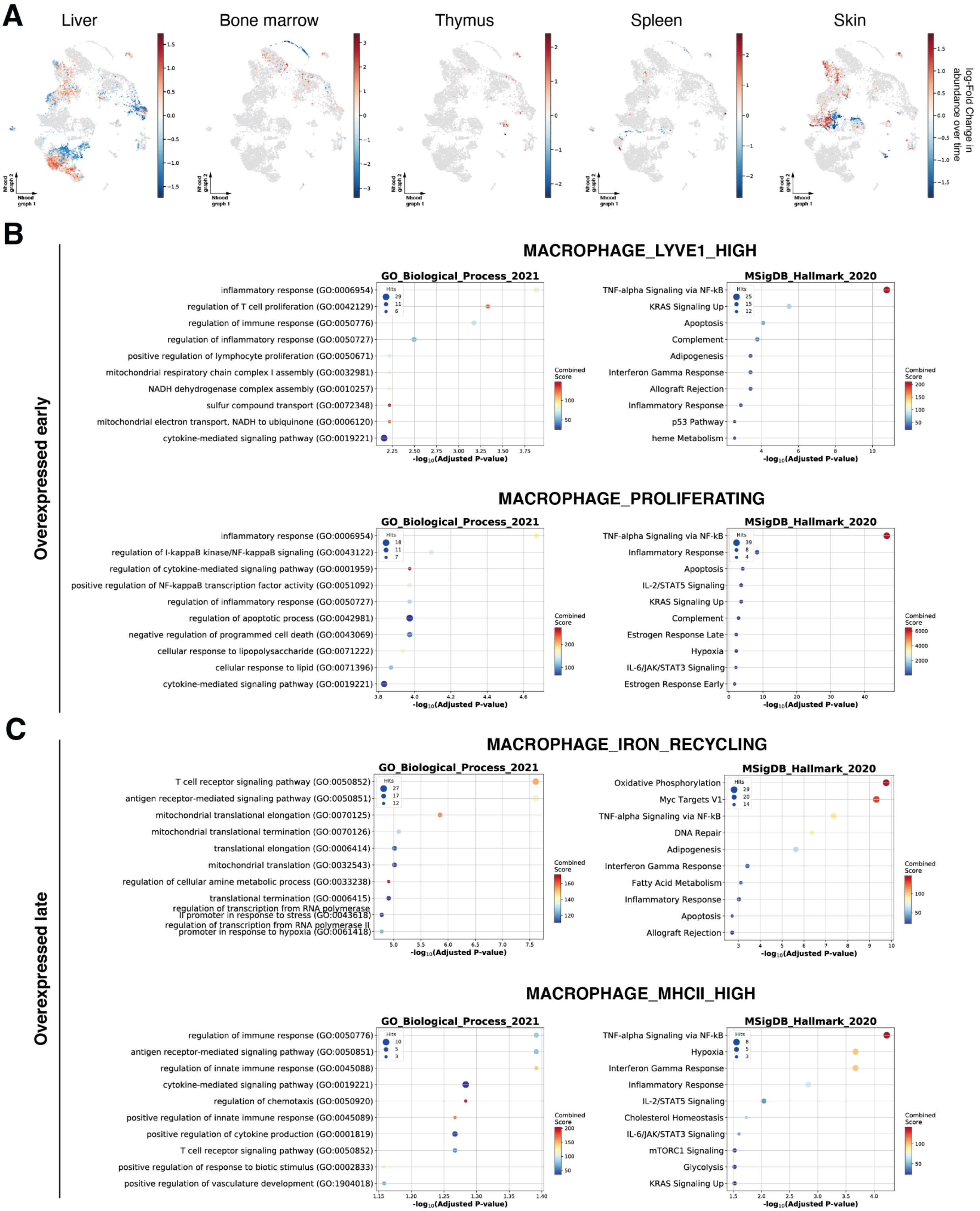
A
Liver
Bone marrow
Thymus
Spleen
Skin
Nhood graph 1
Nhood graph 2
log-Fold Change in abundance over time
B
Overexpressed early
MACROPHAGE_LYVE1_HIGH
GO_Biological_Process_2021
inflammatory response (GO:0006954)
regulation of T cell proliferation (GO:0042129)
regulation of immune response (GO:0050776)
regulation of inflammatory response (GO:0050727)
positive regulation of lymphocyte proliferation (GO:0050671)
mitochondrial respiratory chain complex I assembly (GO:0032981)
NADH dehydrogenase complex assembly (GO:0010257)
sulfur compound transport (GO:0072348)
mitochondrial electron transport, NADH to ubiquinone (GO:0006120)
cytokine-mediated signaling pathway (GO:0019221)
MSigDB_Hallmark_2020
TNF-alpha Signaling via NF-kB
KRAS Signaling Up
Apoptosis
Complement
Adipogenesis
Interferon Gamma Response
Allograft Rejection
Inflammatory Response
p53 Pathway
heme Metabolism
Hits
Combined Score
-log10(Adjusted P-value)
MACROPHAGE_PROLIFERATING
GO_Biological_Process_2021
inflammatory response (GO:0006954)
regulation of I-kappaB kinase/NF-kappaB signaling (GO:0043122)
regulation of cytokine-mediated signaling pathway (GO:0001959)
positive regulation of NF-kappaB transcription factor activity (GO:0051092)
regulation of inflammatory response (GO:0050727)
regulation of apoptotic process (GO:0042981)
negative regulation of programmed cell death (GO:0043069)
cellular response to lipopolysaccharide (GO:0071222)
cellular response to lipid (GO:0071396)
cytokine-mediated signaling pathway (GO:0019221)
MSigDB_Hallmark_2020
TNF-alpha Signaling via NF-kB
Inflammatory Response
Apoptosis
IL-2/STAT5 Signaling
KRAS Signaling Up
Complement
Estrogen Response Late
Hypoxia
IL-6/JAK/STAT3 Signaling
Estrogen Response Early
C
Overexpressed late
MACROPHAGE_IRON_RECYCLING
GO_Biological_Process_2021
T cell receptor signaling pathway (GO:0050852)
antigen receptor-mediated signaling pathway (GO:0050851)
mitochondrial translational elongation (GO:0070125)
mitochondrial translational termination (GO:0070126)
translational elongation (GO:0006414)
mitochondrial translation (GO:0032543)
regulation of cellular amine metabolic process (GO:0033238)
translational termination (GO:0006415)
regulation of transcription from RNA polymerase II promoter in response to stress (GO:0043618)
regulation of transcription from RNA polymerase II promoter in response to hypoxia (GO:0061418)
MSigDB_Hallmark_2020
Oxidative Phosphorylation
Myc Targets V1
TNF-alpha Signaling via NF-kB
DNA Repair
Adipogenesis
Interferon Gamma Response
Fatty Acid Metabolism
Inflammatory Response
Apoptosis
Allograft Rejection
MACROPHAGE_MHCII_HIGH
GO_Biological_Process_2021
regulation of immune response (GO:0050776)
antigen receptor-mediated signaling pathway (GO:0050851)
regulation of innate immune response (GO:0045088)
cytokine-mediated signaling pathway (GO:0019221)
regulation of chemotaxis (GO:0050920)
positive regulation of innate immune response (GO:0045089)
positive regulation of cytokine production (GO:0001819)
T cell receptor signaling pathway (GO:0050852)
positive regulation of response to biotic stimulus (GO:0002833)
positive regulation of vasculature development (GO:1904018)
MSigDB_Hallmark_2020
TNF-alpha Signaling via NF-kB
Hypoxia
Interferon Gamma Response
Inflammatory Response
IL-2/STAT5 Signaling
Cholesterol Homeostasis
IL-6/JAK/STAT3 Signaling
mTORC1 Signaling
Glycolysis
KRAS Signaling Up

**fig. S11: Differential abundance across gestation in myeloid cell populations.** (**A**) Milo neighbourhood embedding of myeloid cells showing differential abundance across gestation. Each point represents a neighbourhood, the layout of points is determined by the position of the neighbourhood index cell in the UMAP in fig. S4G, the size of points is proportional to the number of cells in the neighbourhood. Neighbourhoods are coloured by their log fold change (logFC) in abundance over time, where logFC > 0 indicates significant enrichment in early cells and logFC > 0 indicates significant enrichment in late cells. Only neighbourhoods showing significant differential abundance (SpatialFDR < 10%) are coloured. (**B-C**) Gene set enrichment analysis results for differentially expressed genes in gestation stage-specific neighbourhoods of macrophages. Each plot shows the top 10 significant hits for the gene list. The x-axis shows the negative $\log_{10}$ of the p-value adjusted for multiple testing (Benjamini-Hochberg correction). The size of the dots is proportional to the number of genes associated with the gene set. The colour represents the combined enrichr score calculated with gseapy. Results using the Gene Ontology Biological Process and the MSigDB Hallmark 2020 databases are shown. (B) Gene set enrichment analysis for genes overexpressed in early-specific neighbourhoods of LYVE1$^{hi}$ macrophages and proliferating macrophages. (C) Gene set enrichment analysis for genes overexpressed in late-specific neighbourhoods of iron-recycling macrophages and MHCII$^{hi}$ macrophages.

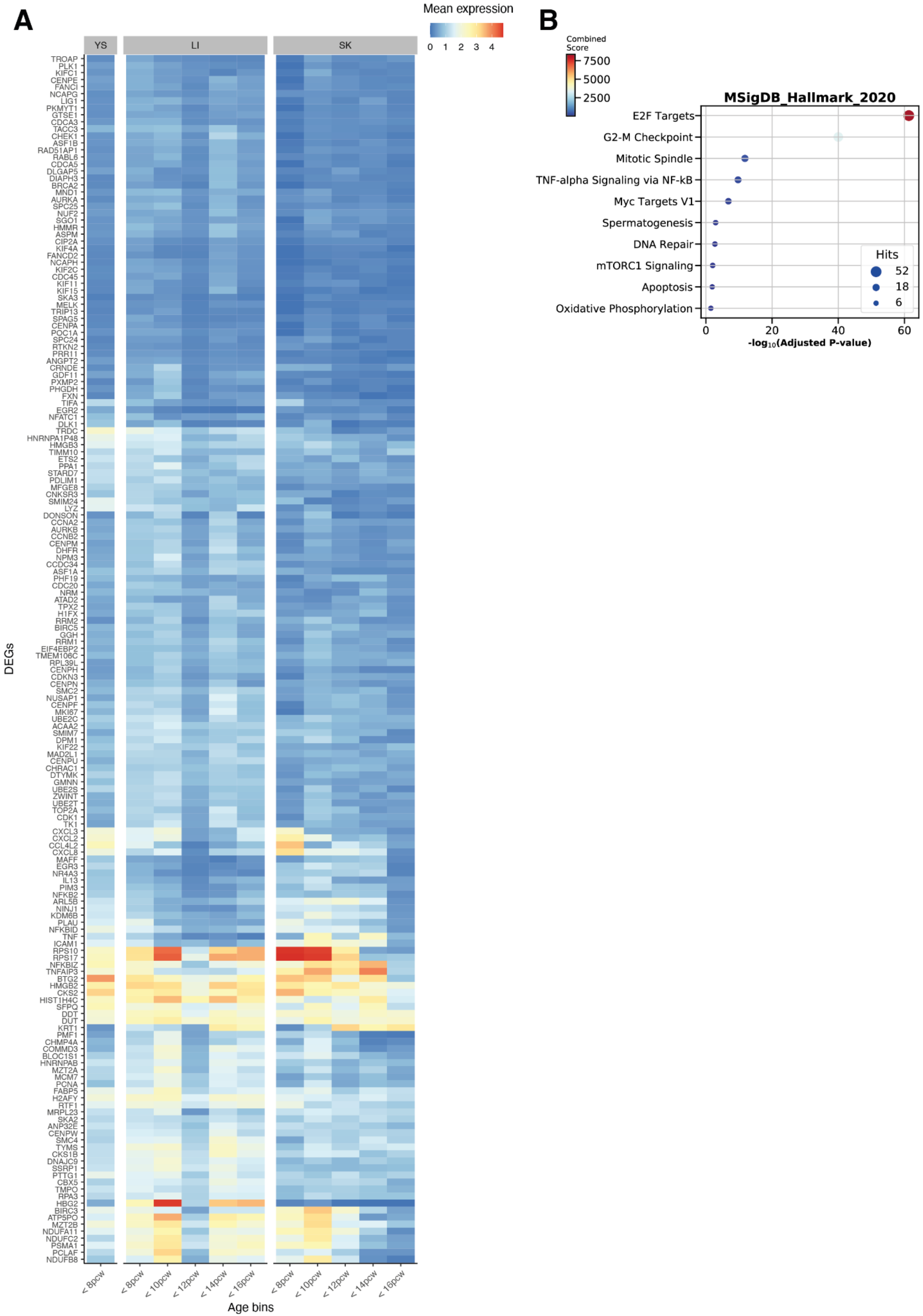

A
B
Mean expression
0 1 2 3 4
YS
LI
SK
DEGs
TROAP
PLK1
KIFC1
CENPE
FANCI
NCAPG
LIG1
PKMYT1
GTSE1
CDCA3
TACC3
CHEK1
ASF1B
RAD51AP1
RABL6
CDCA5
DLGAP5
DIAPH3
BRCA2
MND1
AURKA
SPC25
NUF2
SGO1
HMMR
ASPM
CIP2A
KIF4A
FANCD2
NCAPH
KIF2C
CDC45
KIF11
KIF15
SKA3
MELK
TRIP13
SPAG5
CENPA
POC1A
SPC24
RTKN2
PRR11
ANGPT2
CRNDE
GDF11
PXMP2
PHGDH
FXN
TIFA
EGR2
NFATC1
DLK1
TRDC
HNRNPA1P48
HMGB3
TIMM10
ETS2
PPA1
STARD7
PDLIM1
MFGE8
CNKSR3
SMIM24
LYZ
DONSON
CCNA2
AURKB
CCNB2
CENPM
DHFR
NPM3
CCDC34
ASF1A
PHF19
CDC20
NRM
ATAD2
TPX2
H1FX
RRM2
BIRC5
GGH
RRM1
EIF4EBP2
TMEM106C
RPL39L
CENPH
CDKN3
CENPN
SMC2
NUSAP1
CENPF
MKI67
UBE2C
ACAA2
SMIM7
DPM1
KIF22
MAD2L1
CENPU
CHRAC1
DTYMK
GMNN
UBE2S
ZWINT
UBE2T
TOP2A
CDK1
TK1
CXCL3
CXCL2
CCL4L2
CXCL8
MAFF
EGR3
NR4A3
IL13
PIM3
NFKB2
ARL5B
NINJ1
KDM6B
PLAU
NFKBID
TNF
ICAM1
RPS10
RPS17
NFKBIZ
TNFAIP3
BTG2
HMGB2
CKS2
HIST1H4C
SFPQ
DDT
DUT
KRT1
PMF1
CHMP4A
COMMD3
BLOC1S1
HNRNPAB
MZT2A
MCM7
PCNA
FABP5
H2AFY
RTF1
MRPL23
SKA2
ANP32E
CENPW
SMC4
TYMS
CKS1B
DNAJC9
SSRP1
PTTG1
CBX5
TMPO
RPA3
HBG2
BIRC3
ATP5PO
MZT2B
NDUFA11
NDUFC2
PSMA1
PCLAF
NDUFB8
< 8pcw
< 8pcw
< 10pcw
< 12pcw
< 14pcw
< 16pcw
< 8pcw
< 10pcw
< 12pcw
< 14pcw
< 16pcw
Age bins
Combined Score
7500
5000
2500
MSigDB_Hallmark_2020
E2F Targets
G2-M Checkpoint
Mitotic Spindle
TNF-alpha Signaling via NF-kB
Myc Targets V1
Spermatogenesis
DNA Repair
mTORC1 Signaling
Apoptosis
Oxidative Phosphorylation
Hits
52
18
6
0
20
40
60
-log10(Adjusted P-value)

**fig. S12: Differential expression analysis on early-specific neighbourhoods of mast cells.** (**A**) Average expression by time point of 185 genes over-expressed in early-specific neighbourhoods of mast cells. (**B**) Gene set enrichment analysis results using the MSigDB Hallmark 2020 database. The x-axis shows the negative $\log_{10}$ of the p-value adjusted for multiple testing (Benjamini-Hochberg correction). The size of the dots is proportional to the number of genes associated with the gene set. The colour represents the combined enrichr score calculated with gseapy.

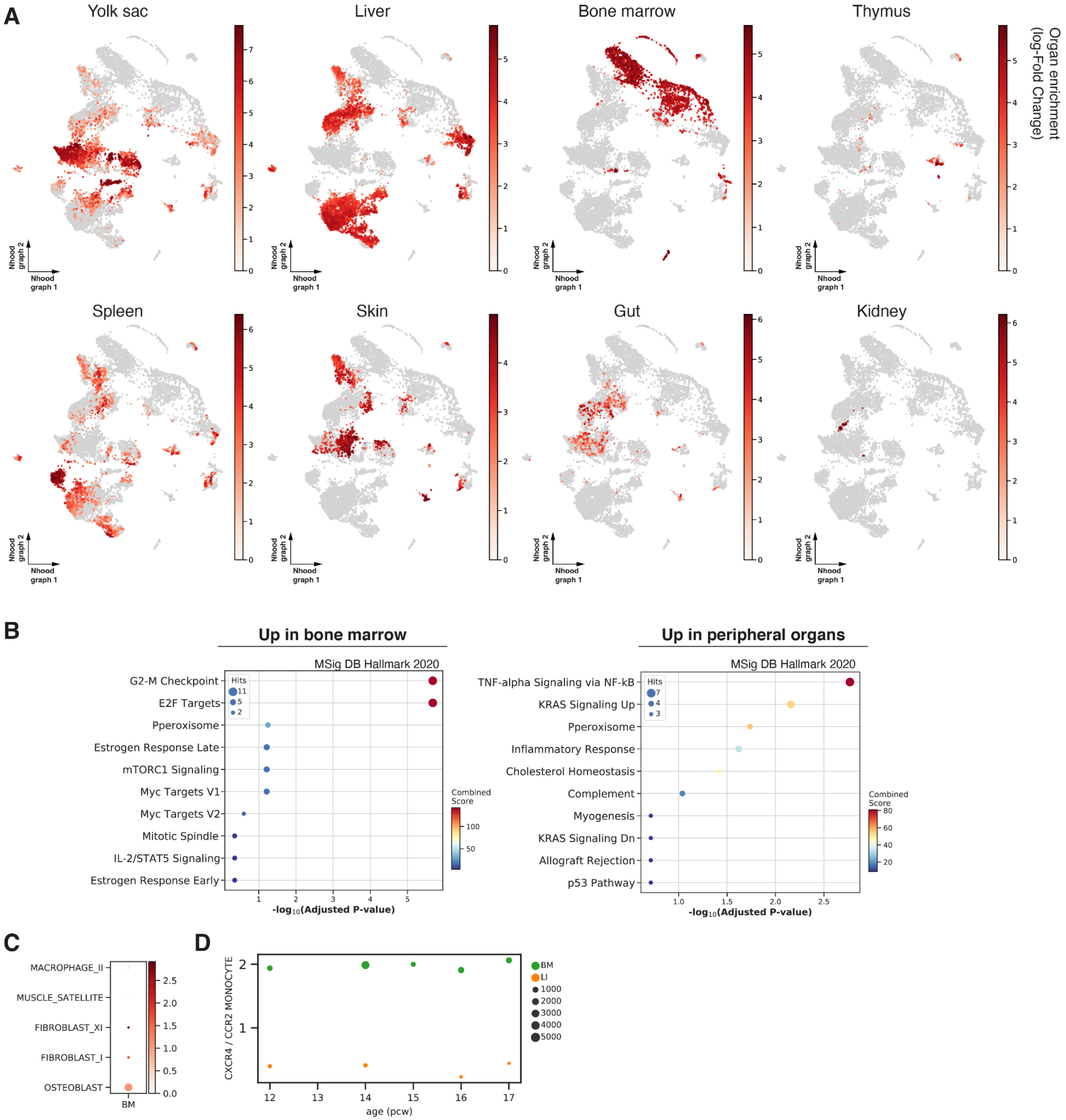
A
Yolk sac
Liver
Bone marrow
Thymus
Spleen
Skin
Gut
Kidney
Organ enrichment (log-Fold Change)
Nhood graph 1
Nhood graph 2
B
Up in bone marrow
MSig DB Hallmark 2020
G2-M Checkpoint
E2F Targets
Pperoxisome
Estrogen Response Late
mTORC1 Signaling
Myc Targets V1
Myc Targets V2
Mitotic Spindle
IL-2/STAT5 Signaling
Estrogen Response Early
Hits
Combined Score
-log10(Adjusted P-value)
Up in peripheral organs
MSig DB Hallmark 2020
TNF-alpha Signaling via NF-kB
KRAS Signaling Up
Pperoxisome
Inflammatory Response
Cholesterol Homeostasis
Complement
Myogenesis
KRAS Signaling Dn
Allograft Rejection
p53 Pathway
C
MACROPHAGE_II
MUSCLE_SATELLITE
FIBROBLAST_XI
FIBROBLAST_I
OSTEOBLAST
BM
D
CXCR4 / CCR2 MONOCYTE
age (pcw)
BM
LI

**fig. S13: Differential abundance across organs in myeloid cell populations.** (**A**) Milo neighbourhood embedding of myeloid cells. Each point represents a neighbourhood, the layout of points is determined by the position of the neighbourhood index cell in the UMAP in fig. S4G, the size of points is proportional to the number of cells in the neighbourhood. Neighbourhoods are coloured by their log fold change in abundance between the specified organ and all other organs. Only neighbourhoods showing significant enrichment (SpatialFDR < 10% and logFC ≥ 2) are coloured. (**B**) Dot plot of enrichment analysis results on genes upregulated (left) and downregulated (right) in bone marrow CCR2$^{hi}$ monocytes compared to other organs. The x-axis represents the negative $\log_{10}$ of p-values adjusted for multiple testing (Benjamini-Hochberg correction) The y-axis shows the top 10 enriched gene sets (using the MSigDB Hallmark 2020 database). The size of the dots is proportional to the number of genes associated with the gene set. The colour represents the combined enrichr score calculated with gseapy. (**C**) Dot plot of *CXCL12* expression in cell populations in the bone marrow. The colour represents the average expression level (normalised and log-transformed counts) and the size represents the cell count of each cell type within bone marrow. Only cell populations with average expression > 1, and cell count > 10 are shown. (**D**) Ratio of abundance of CXCR4$^{hi}$ monocytes over CCR2$^{hi}$ monocytes in bone marrow (BM) and liver (LI) over gestational age. Dot size is proportional to the total number of cells at the corresponding time point.

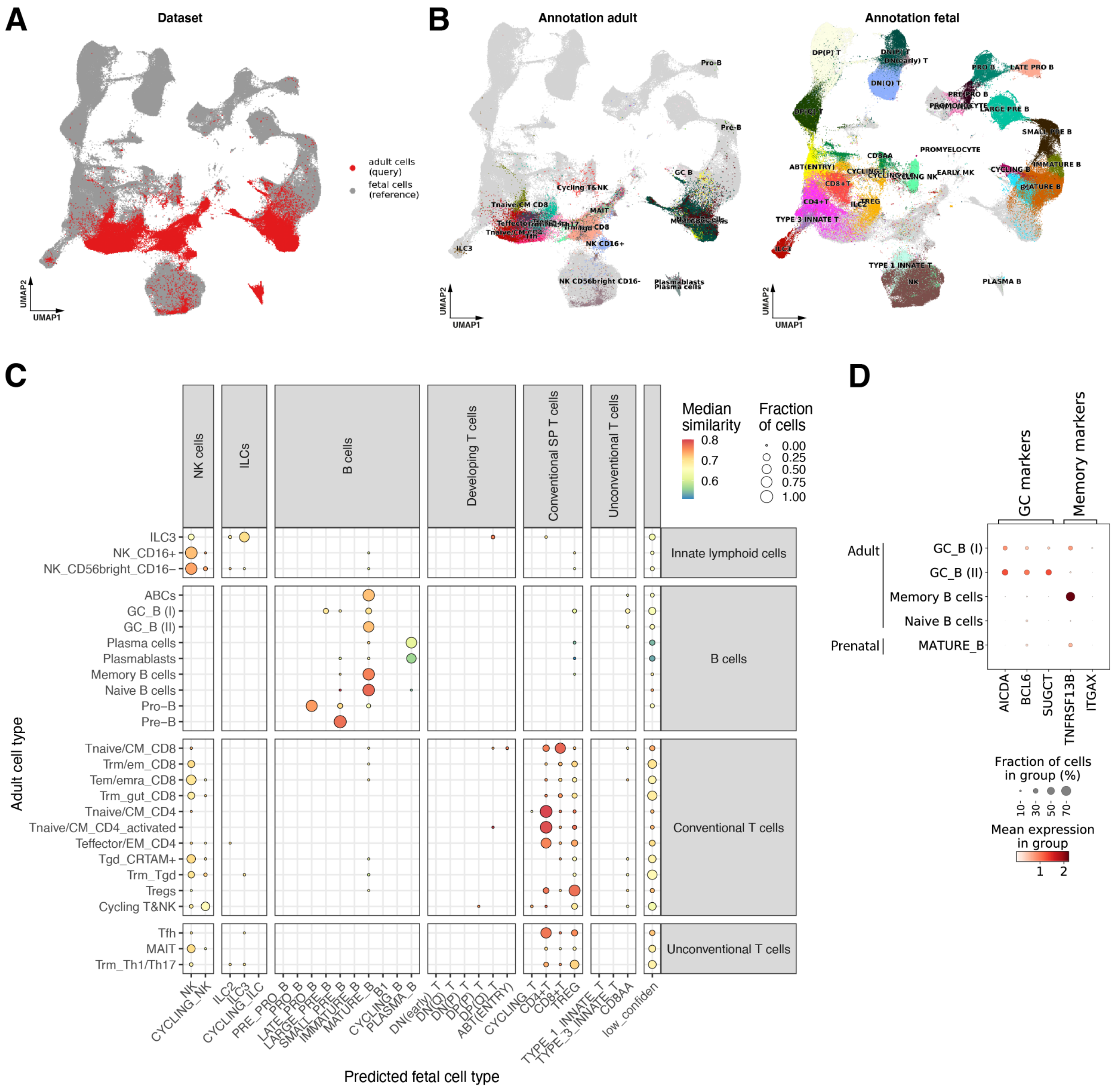


**fig. S14: Mapping of adult lymphoid cells to prenatal reference with transfer learning**. (**A**) UMAP embeddings of mapping of adult lymphoid cells (13,062) to prenatal lymphoid reference (218,758 cells) using scArches on scVI model. Points are coloured by the dataset of origin. (**B**) UMAP embedding as in A, points are coloured by cell population annotation label, for adult cells (left) and prenatal cells (right). (**C**) Correspondence between prenatal and adult myeloid transcriptional phenotypes estimated by label transfer after mapping with scArches. The dot size is proportional to the fraction of cells in the adult population (y-axis) with a given predicted prenatal cell population label (x-axis). The dot colour denotes the median similarity of the adult cells to prenatal cells in the common embedding (see Methods). Adult cells where less than 50% of prenatal neighbours have a uniform annotation are labelled as 'low confidence'. (**D**) Dot plot of expression of germinal centre (GC) and memory B cell marker genes in Adult and prenatal mature B cell subsets.

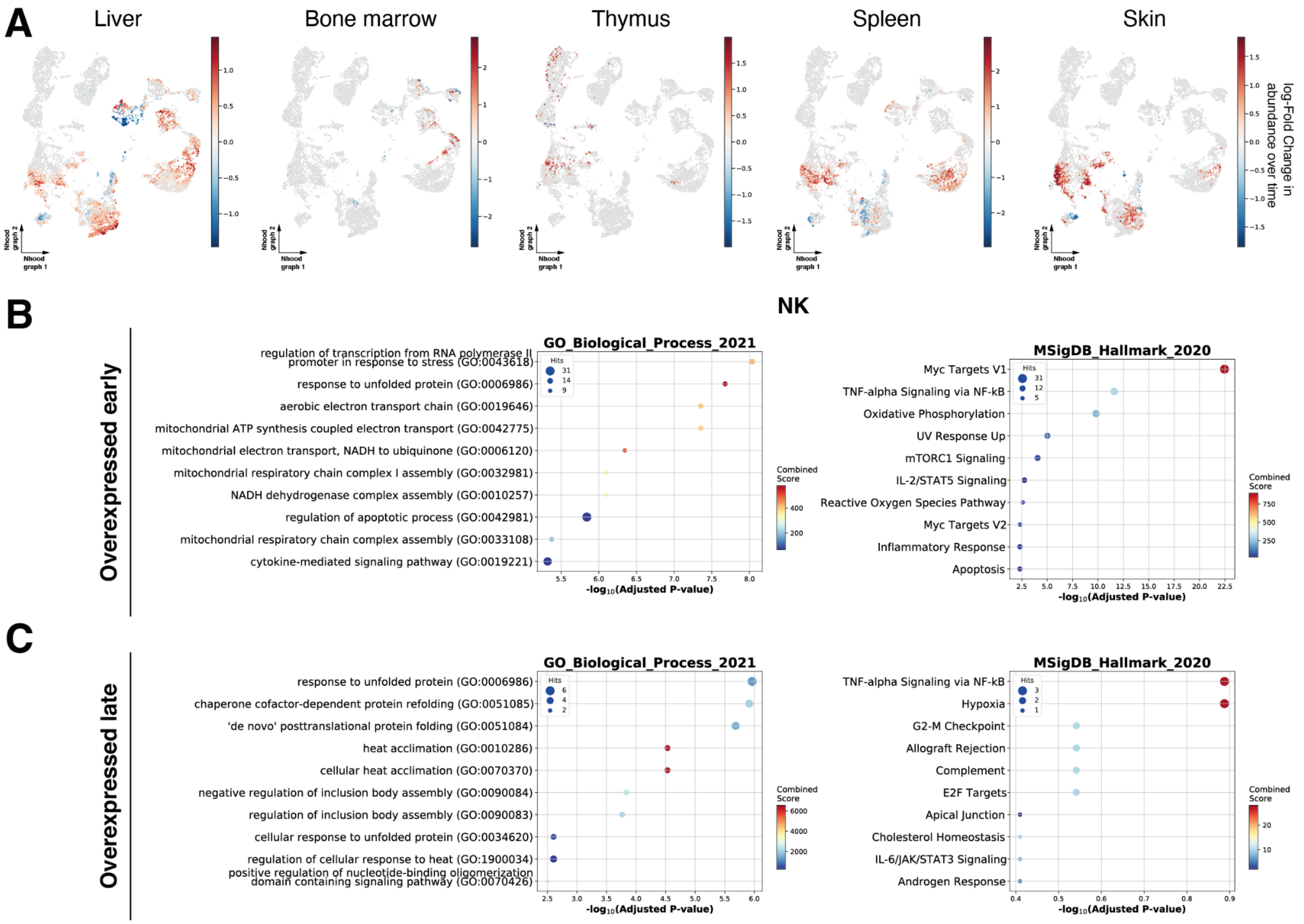


**fig. S15: Differential abundance across gestation in lymphoid cell populations.** (**A**) Milo neighbourhood embedding of lymphoid cells showing differential abundance across gestation. Each point represents a neighbourhood, the layout of points is determined by the position of the neighbourhood index cell in the UMAP in fig. S5I, the size of points is proportional to the number of cells in the neighbourhood. Neighbourhoods are coloured by their log fold change (logFC) in abundance over time, where logFC > 0 indicates significant enrichment in early cells and logFC > 0 indicates significant enrichment in late cells. Only neighbourhoods showing significant differential abundance (SpatialFDR < 10%) are coloured. (**B-C**) Gene set enrichment analysis results for differentially expressed genes in early-specific neighbourhoods (B) and late specific neighbourhoods (C) of NK cells. Each plot shows the top 10 significant hits for the gene set. The x-axis shows the negative $\log_{10}$ of the p-value adjusted for multiple testing (Benjamini-Hochberg correction). The size of the dots is proportional to the number of genes associated with the gene set. The colour represents the combined enrichr score calculated with gseapy. Results using the Gene Ontology Biological Process and the MSigDB Hallmark 2020 databases are shown.

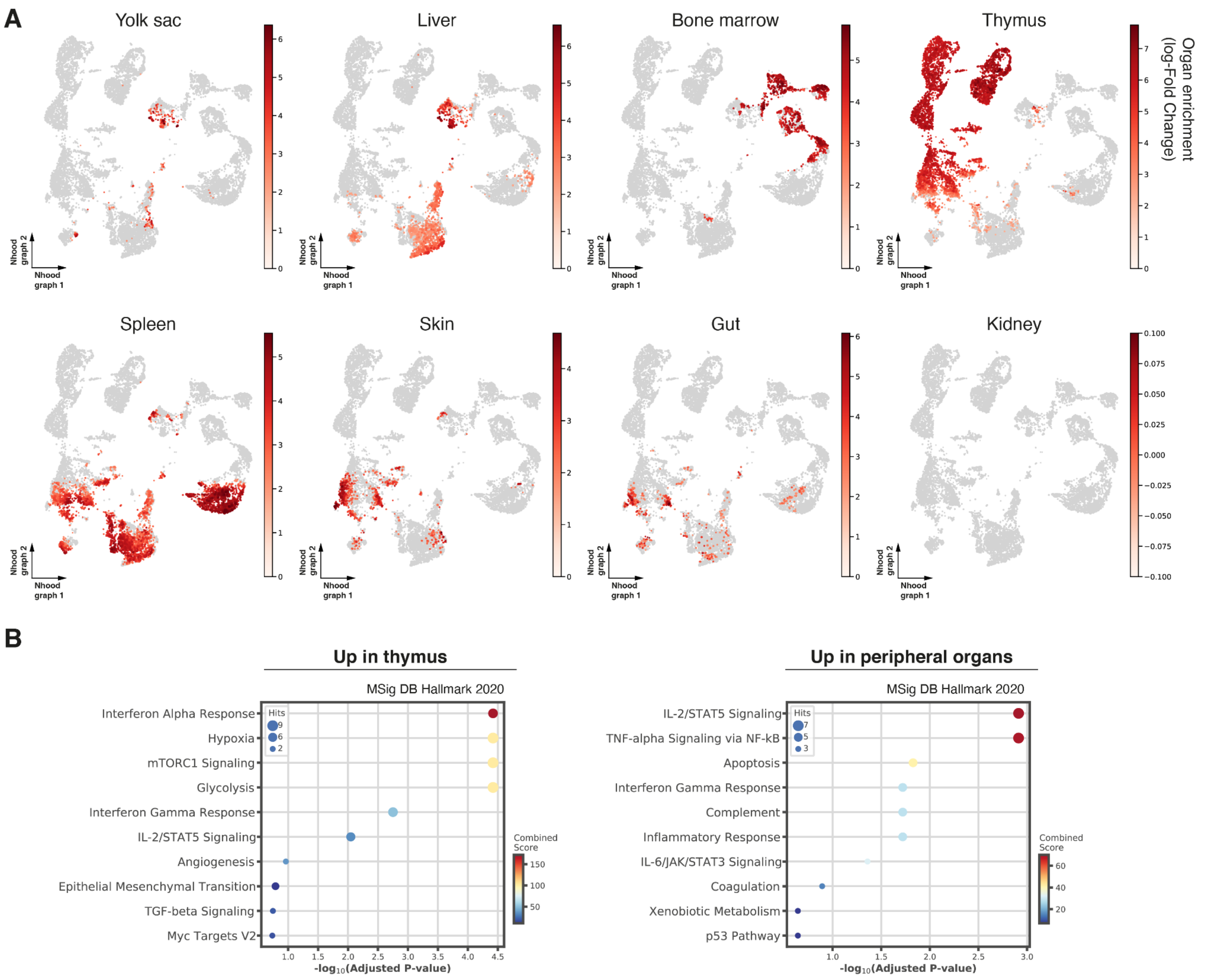


**fig. S16: Differential abundance across organs in lymphoid compartment.** (**A**) Milo neighbourhood embedding of lymphoid cells showing differential abundance between organs. Each point represents a neighbourhood, the layout of points is determined by the position of the neighbourhood index cell in the UMAP in Fig. S4I, the size of points is proportional to the number of cells in the neighbourhood. Neighbourhoods are coloured by their log fold change in abundance between the specified organ and all other organs. Only neighbourhoods showing significant enrichment (SpatialFDR < 10% and log-fold change > 2) are coloured. (**B**) Dot plot of enrichment analysis results on genes upregulated (left) and downregulated (right) in thymic mature T cells compared to other organs. The x-axis represents the negative $\log_{10}$ of p-values adjusted for multiple testing (Benjamini-Hochberg correction) The y-axis shows the top 10 enriched gene sets (using the MSigDB Hallmark 2020 database). The size of the dots is proportional to the number of genes associated to the gene set. The colour represents the combined enrichr score calculated with gseapy.

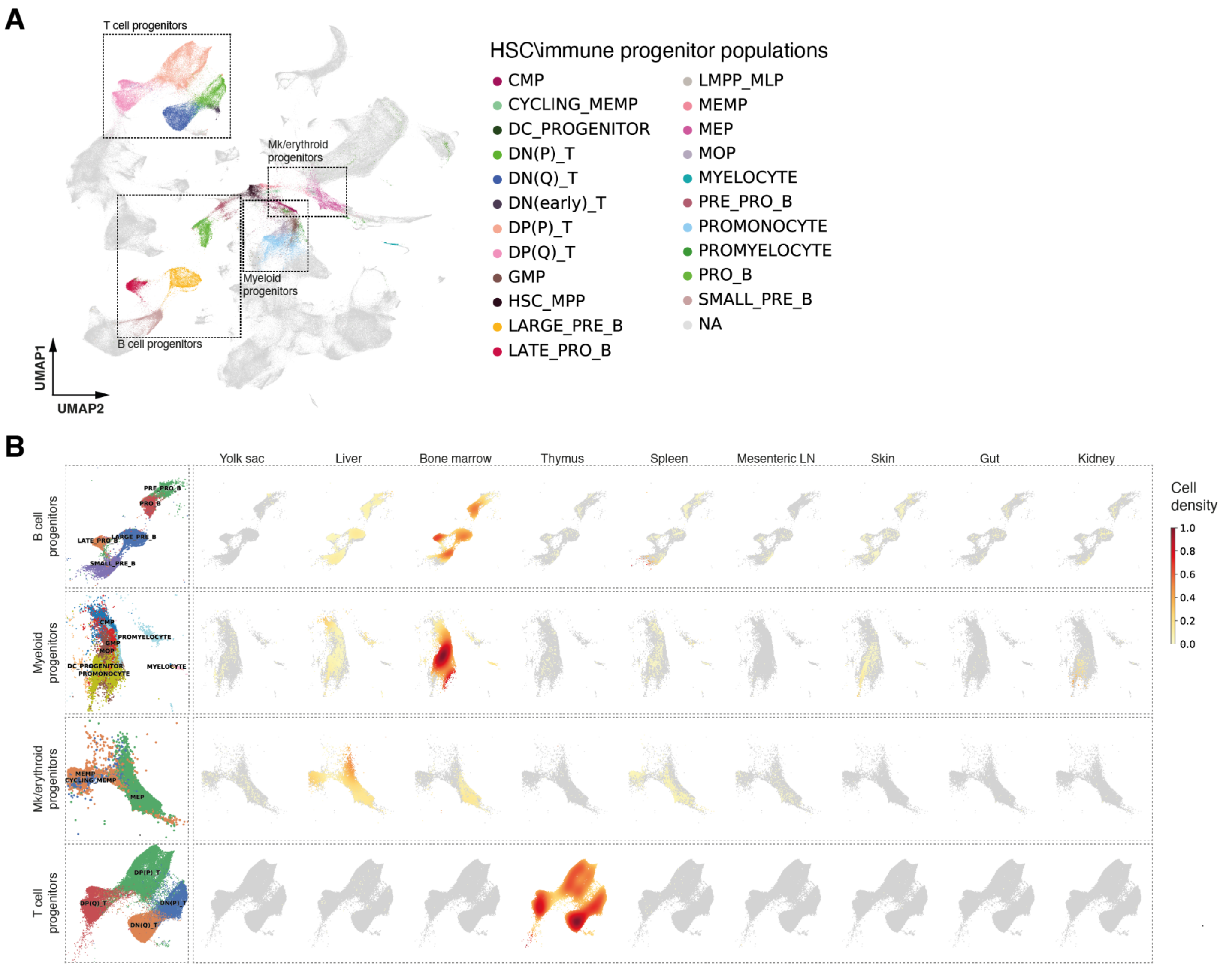


**fig. S17. Full spectrum of haematopoietic progenitors in peripheral organs.** (**A**) UMAP embedding of all immune and blood cells, highlighting the progenitor cell populations. Dashed boxes highlight lineage populations shown in B. (**B**) Density plot of cells from each organ on a subset of UMAP embedding for different haematopoietic lineages. Density is calculated over all immune cells within each organ.

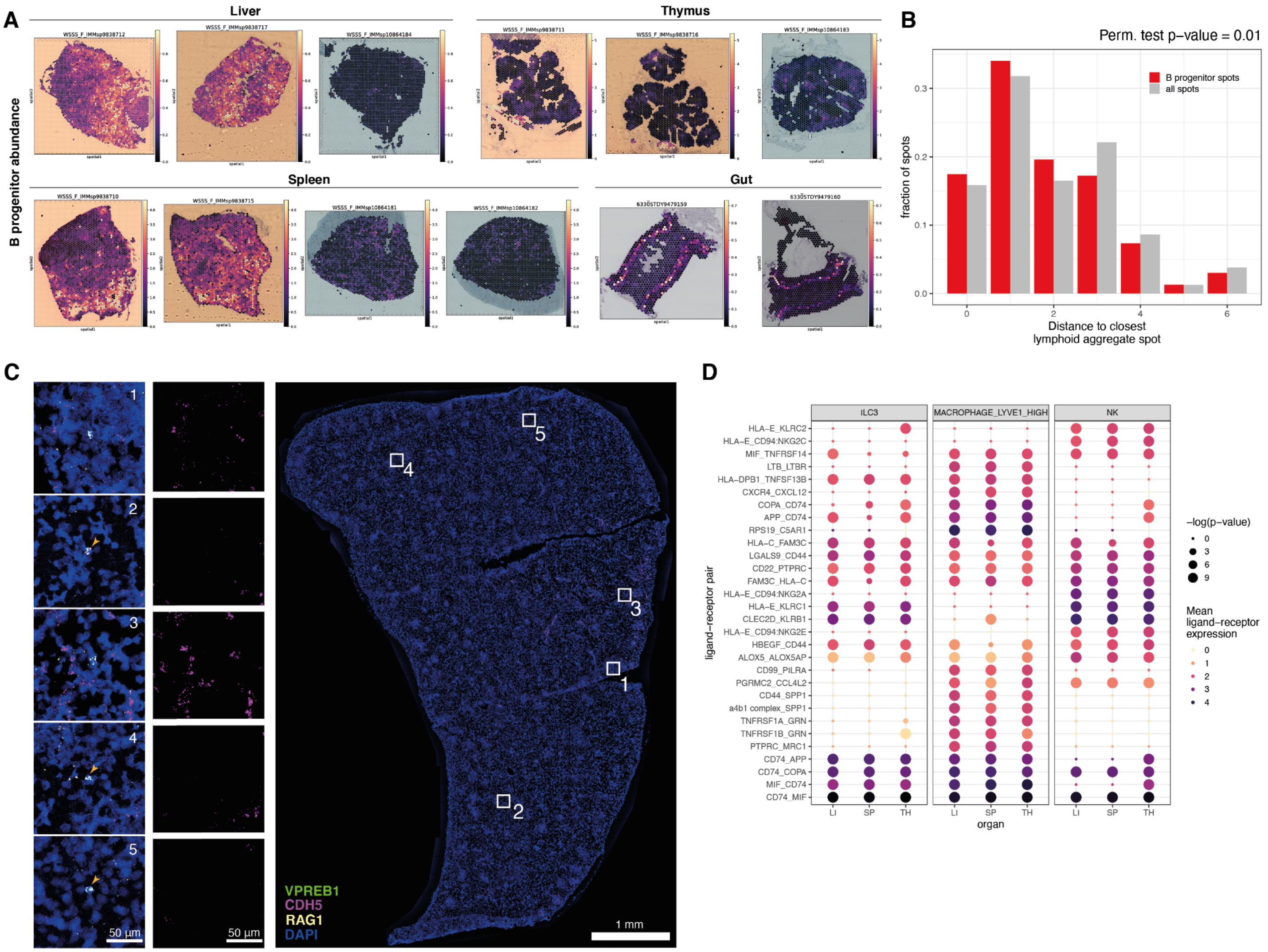


**fig. S18. System-wide B lymphopoiesis.** (**A**) Sum of abundances of B progenitor cell populations in spatial transcriptomics slides estimated with cell2location. (**B**) Distribution of distance to the closest spot assigned as splenic lymphoid aggregate microenvironment from spots containing B cell progenitors (red) and from all analysed spots (grey). Distance is measured as Euclidean distance of spots in spatial coordinates. We test if the distance to lymphoid aggregates is significantly smaller in B progenitor spots compared to other spots with a permutation test (5000 samples). Spots were assigned to the lymphoid aggregate microenvironment if the NMF factor value for the microenvironment (see Fig. 1D, 'B cell zone') was above the 95% quantile for that slide. We considered spots to contain B cell progenitors if the sum of abundances of the B cell progenitors was above the 95% quantile for that slide. (**C**) Multiplex smFISH staining of DAPI, *CDH5* for endothelial cells, and *VPREB1*, *RAG1* for B progenitors in the human fetal spleen at 14 pcw. Left: 5 cells highlighted from corresponding regions in the right panel overview matched by numbers. Middle: *CDH5* channel depicting endothelial cells. Right: full section view with the areas of interest boxed in white. The orange arrows point to B progenitors away from blood vessels. (**D**) Predicted cell-cell interactions between B progenitors and co-localising cell types (ILC3, LYVE1$^{hi}$ Macrophage and NK cells) from CellPhoneDB across liver (LI), spleen (SP) and thymus (TH) (see Methods). The first gene in each ligand-receptor pair is from B progenitors and the second from the interacting cell type. The colour represents the average expression values of the ligand and receptor within their corresponding cell types, and the size represents -log(p-value).

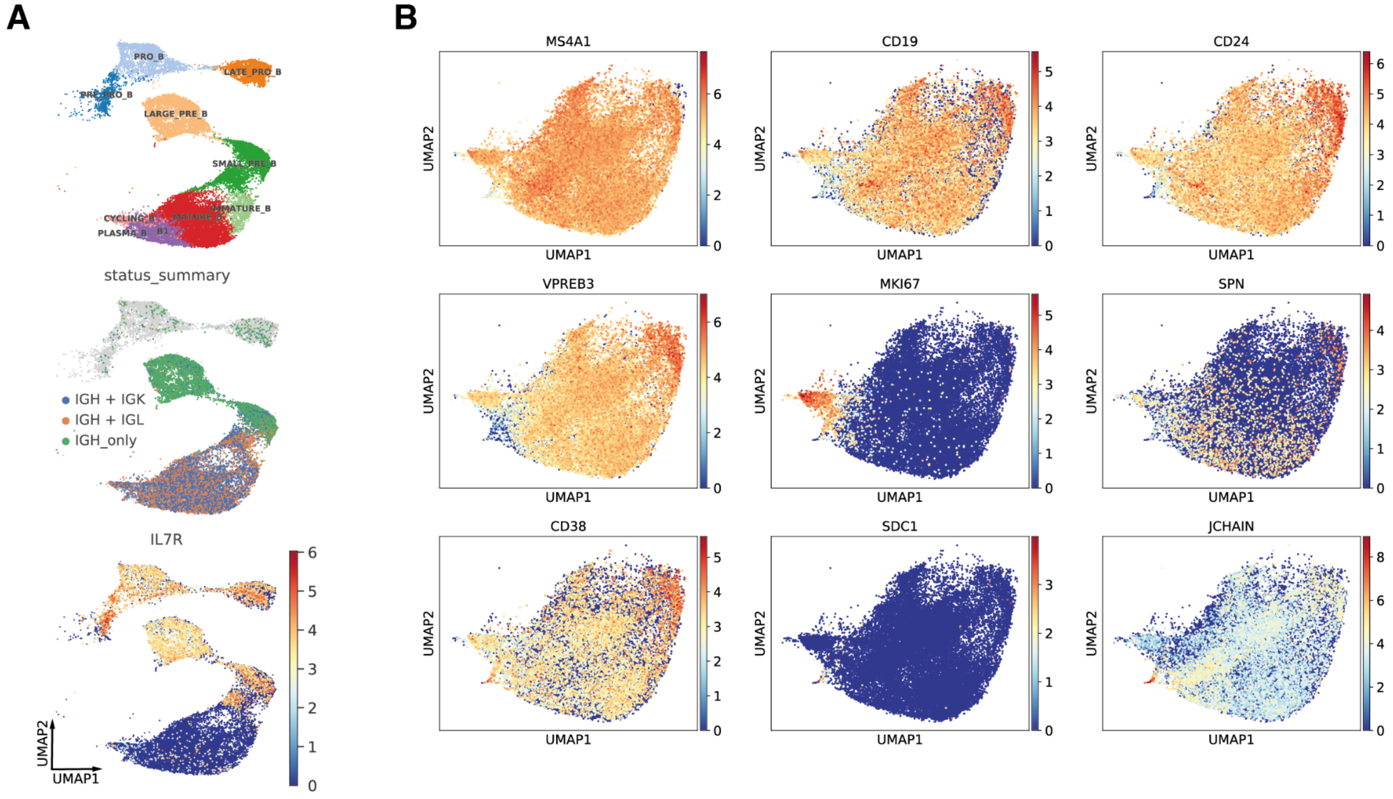


**fig. S19. Characterisation of B cells.** (**A**) Close-up view of B cell populations on the UMAP embedding of all lymphoid cells (as shown in fig. S5I), coloured by annotated cell population identity (top), status summary of cells expressing productive heavy (IGH) and/or light chains (IGK or IGL) of BCR from single cell BCR sequencing (middle), and coloured by *IL7R* expression pattern (bottom). (**B**) Marker gene expression patterns overlaid onto the same UMAP plot in Fig. 4A.

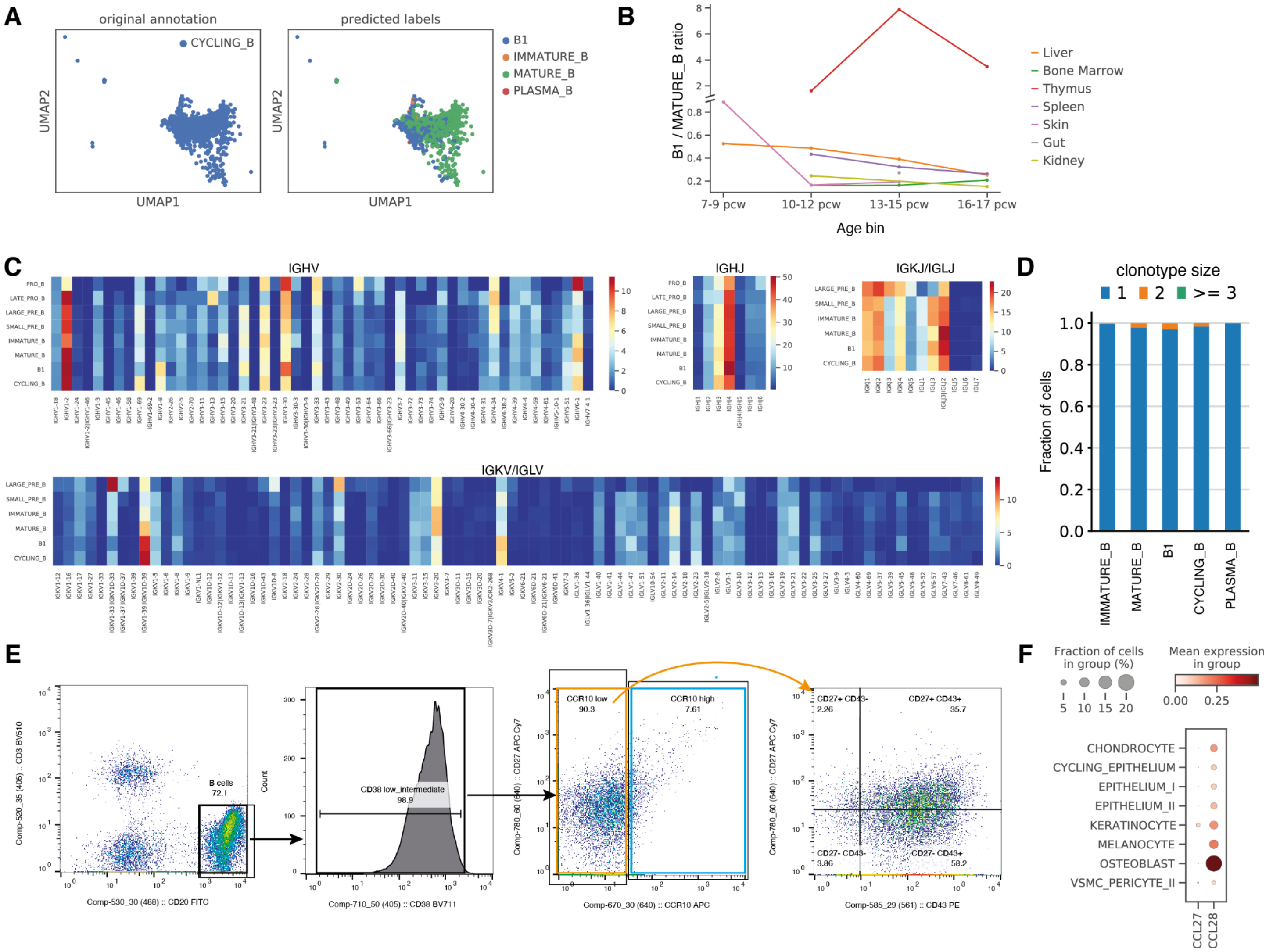


**fig. S20. Characterisation of putative B1 cells.** (**A**) Close-up view of Cycling B cell population on UMAP embedding of all lymphoid cells (as shown in fig. S5I), coloured by original cell population annotation (left) and annotations predicted by logistic regression trained on all mature B cell subsets (right). (**B**) Ratio of B1 cell number over Mature B cell number in different organs across different gestational age bins. (**C**) Heat map showing the percentage of each BCR heavy (IGHV, IGHJ) and light chain (IGKV/IGLV, IGKJ/IGLJ) V and J gene segments present in different B cell subtypes. (**D**) Barplot of cell fractions with different clonotype size across different mature B cell subtypes. (**E**) Representative flow cytometry plots showing the sorting strategy for the ELISpot experiment shown in Fig. 4E. The splenic B cells were gated from live single cells which were $CD3^{-}CD20^{+}$, excluding the top 1% of cells expressing the highest level of CD38 to avoid plasma cells (which should also be CD20 low and therefore not gated in), and split the rest into four fractions - $CCR10^{hi}$, $CCR10^{lo}CD27^{+}CD43^{+}$, $CCR10^{lo}CD27^{-}CD43^{+}$, $CCR10^{lo}CD27^{-}CD43^{-}$. We then performed an ELISpot experiment on all four fractions without any stimulation. (**F**) Dot plot showing gene expressions of *CCL27* and *CCL28* within the stromal cell populations. Only cell types with log normalised expression of *CCL27* or that of *CCL28* above 0.05 are shown here.

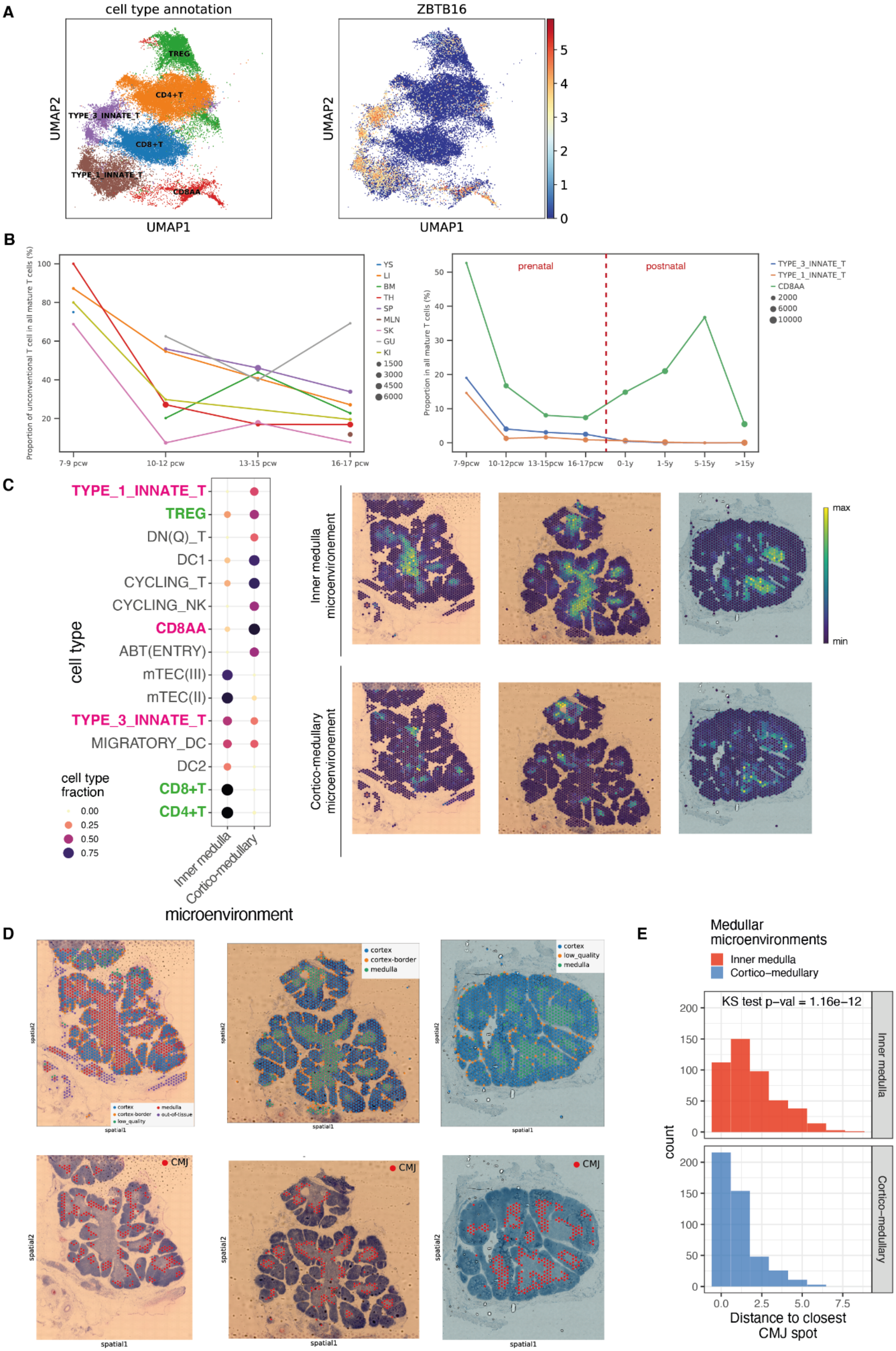
A
cell type annotation
ZBTB16
UMAP2
UMAP1
TREG
CD4+T
TYPE_3_INNATE_T
CD8+T
TYPE_1_INNATE_T
CD8AA
B
Proportion of unconventional T cell in all mature T cells (%)
YS
LI
BM
TH
SP
MLN
SK
GU
KI
7-9 pcw
10-12 pcw
13-15 pcw
16-17 pcw
prenatal
postnatal
Proportion in all mature T cells (%)
TYPE_3_INNATE_T
TYPE_1_INNATE_T
CD8AA
7-9pcw
10-12pcw
13-15pcw
16-17pcw
0-1y
1-5y
5-15y
>15y
C
TYPE_1_INNATE_T
TREG
DN(Q)_T
DC1
CYCLING_T
CYCLING_NK
CD8AA
ABT(ENTRY)
mTEC(III)
mTEC(II)
TYPE_3_INNATE_T
MIGRATORY_DC
DC2
CD8+T
CD4+T
cell type
cell type fraction
Inner medulla
Cortico-medullary
microenvironment
Inner medulla microenvironement
Cortico-medullary microenvironement
max
min
D
cortex
cortex-border
low_quality
medulla
out-of-tissue
spatial1
spatial2
CMJ
E
Medullar microenvironments
Inner medulla
Cortico–medullary
KS test p–val = 1.16e–12
count
Distance to closest CMJ spot

**fig. S21. Distribution of unconventional T cells across gestation and in thymic tissue.** (**A**) Left: Close-up view of mature T cells on UMAP embedding of all NK/T cells (as shown in fig. S4L). Type 1 Innate T, 3 Innate T and CD8AA contain both αβT cells and γδT cells. Right: *ZBTB16* expression pattern overlaid onto the same UMAP plot. (**B**) Left: proportion of unconventional T cells in all mature T cells in different organs across different gestational age bins. Point size represents the number of mature T cells in a given organ within that age bin. Lines and points are colour-coded by organs (YS: yolk sac; LI: liver; BM: bone marrow; TH: thymus; SP: spleen; MLN: mesenteric lymph node; SK: skin; GU: gut; KI: kidney). Right: proportion of each unconventional T cell subtype in all mature T cells in thymus across different age groups. Point size represents the number of mature T cells in thymus within that age bin. (**C**) Left: cell type contributions to medullary microenvironments containing mature T cells in thymus, identified with non-negative matrix factorisation of spatial cell type abundances estimated with cell2location. The colour and the size of the dots represent the relative fraction of the cell population assigned to the microenvironment. Unconventional T cell types are highlighted in magenta. Conventional T cell types are highlighted in green. Right: spatial locations of medullary microenvironments on different thymic slides, with the colour representing the weighted contribution of each microenvironment to each spot. (**D**) Top: annotation of tissue regions on Visium spots inferred by clustering of H&E image features. Bottom: location of interface region between cortex and medulla region, which we consider as the cortico-medullary junction (CMJ). (**E**) Histogram of Euclidean distance to the nearest CMJ spot for spots assigned to inner-medulla microenvironment (red) or cortico-medullary microenvironment (blue). Spots were assigned to a microenvironment if the NMF factor value for the microenvironment (see C) was above the 90% quantile. The p-value for the Kolmogorov–Smirnov test comparing the two distributions is reported.

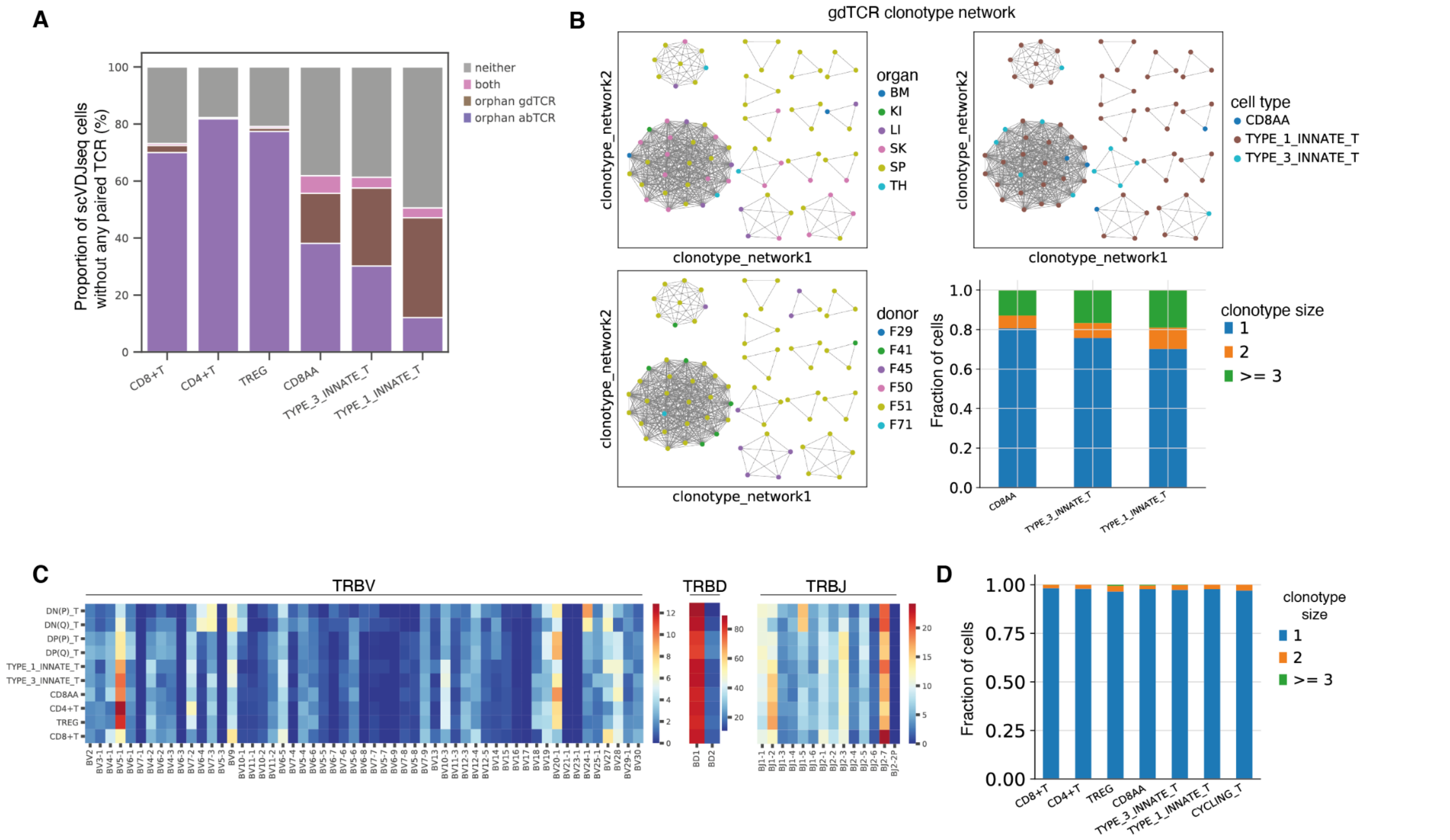


**fig. S22. T cell receptor (TCR) sequence analysis.** (**A**) Proportions of cells without any paired TCR expressing orphan γδTCR, i.e. one of γ or δ chain, or orphan αβTCR, i.e. one of α or β chain, or both or neither. (**B**) Top left: clonotype network graph of γδTCR. Each fully connected subnetwork represents a clonotype cluster, with each dot representing a cell. The dots are colour-coded by organs (YS: yolk sac; LI: liver; BM: bone marrow; TH: thymus; SP: spleen; MLN: mesenteric lymph node; SK: skin; GU: gut; KI: kidney). Top right: γδTCR clonotype network graph colour-coded by cell types. Bottom left: γδTCR clonotype network graph colour-coded by donors. Bottom right: barplot of cell fractions with different γδTCR clonotype size across different unconventional T cell subtypes. (**C**) Heat map showing the percentage of each TRBV, TRBD and TRBJ gene segment present in different T cell subtypes. (**D**) Barplot of cell fractions with different clonotype size across different mature T cell subtypes.

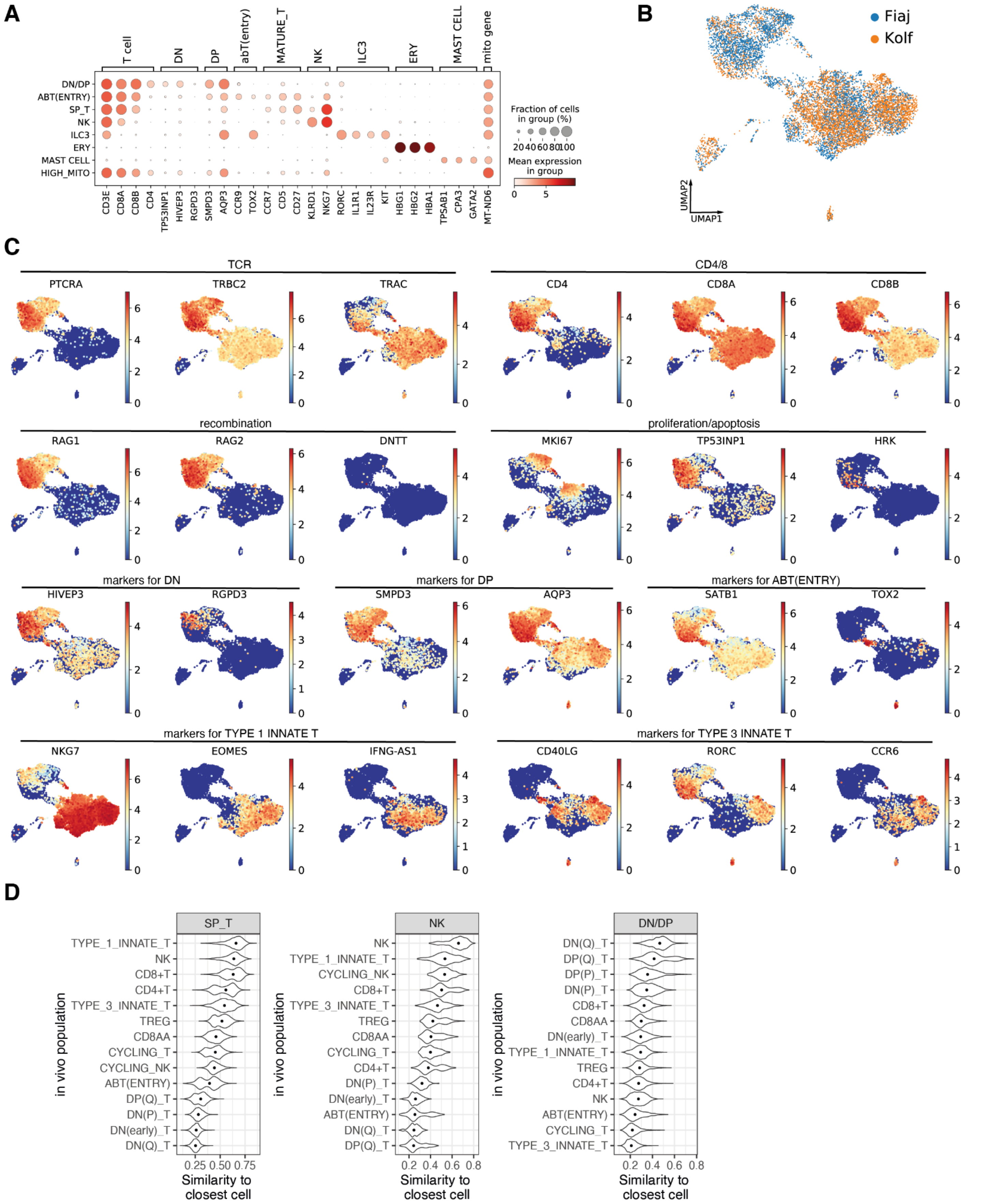
A
T cell
DN
DP
abT(entry)
MATURE_T
NK
ILC3
ERY
MAST CELL
mito gene
DN/DP
ABT(ENTRY)
SP_T
NK
ILC3
ERY
MAST CELL
HIGH_MITO
CD3E
CD8A
CD8B
CD4
TP53INP1
HIVEP3
RGPD3
SMPD3
AQP3
CCR9
TOX2
CCR7
CD5
CD27
KLRD1
NKG7
RORC
IL1R1
IL23R
KIT
HBG1
HBG2
HBA1
TPSAB1
CPA3
GATA2
MT-ND6
Fraction of cells in group (%)
20 40 60 80 100
Mean expression in group
0 5
B
Fiaj
Kolf
UMAP2
UMAP1
C
TCR
PTCRA
TRBC2
TRAC
CD4/8
CD4
CD8A
CD8B
recombination
RAG1
RAG2
DNTT
proliferation/apoptosis
MKI67
TP53INP1
HRK
markers for DN
HIVEP3
RGPD3
markers for DP
SMPD3
AQP3
markers for ABT(ENTRY)
SATB1
TOX2
markers for TYPE 1 INNATE T
NKG7
EOMES
IFNG-AS1
markers for TYPE 3 INNATE T
CD40LG
RORC
CCR6
D
SP_T
TYPE_1_INNATE_T
NK
CD8+T
CD4+T
TYPE_3_INNATE_T
TREG
CD8AA
CYCLING_T
CYCLING_NK
ABT(ENTRY)
DP(Q)_T
DN(P)_T
DN(early)_T
DN(Q)_T
0.25 0.50 0.75
NK
NK
TYPE_1_INNATE_T
CYCLING_NK
CD8+T
TYPE_3_INNATE_T
TREG
CD8AA
CYCLING_T
CD4+T
DN(P)_T
DN(early)_T
ABT(ENTRY)
DN(Q)_T
DP(Q)_T
0.2 0.4 0.6 0.8
DN/DP
DN(Q)_T
DP(Q)_T
DP(P)_T
DN(P)_T
CD8+T
CD8AA
DN(early)_T
TYPE_1_INNATE_T
TREG
CD4+T
NK
ABT(ENTRY)
CYCLING_T
TYPE_3_INNATE_T
0.2 0.4 0.6 0.8
in vivo population
Similarity to closest cell

**fig. S23. Analysis of Artificial Thymic Organoids scRNA-seq data.** (**A**) Dot plot of marker genes for ATO cell populations. (**B**) Cells coloured by the starting iPSC lines in ATO overlaid on UMAP embedding shown in Fig. 5F. (**C**) Expression of T cell marker genes in ATO overlaid on UMAP embedding shown in Fig. 5F. (**D**) Violin plots of similarity to closest *in vivo* cell (x-axis) for each cell type in the *in vivo* dataset (y-axis) for *in vitro* single-positive T cells (SP_T), NK cells and developing T cells (DN/DP). Similarities are calculated in the scVI latent space for lymphoid cells after mapping *in vitro* cells with scArches.

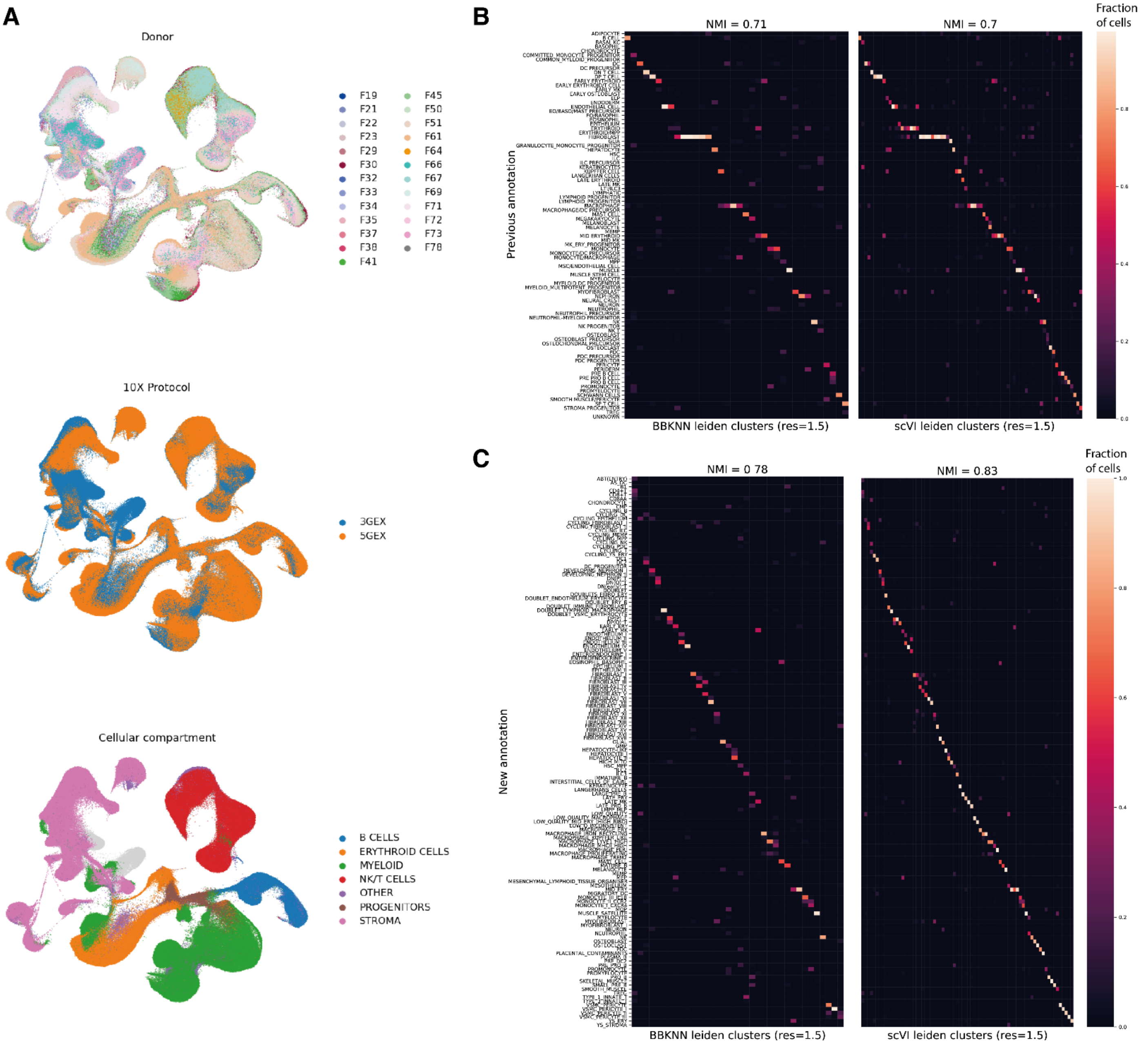


**fig. S24: Comparison across data integration methods.** (**A**) UMAP embeddings of scRNA-seq profiles after data integration with BBKNN coloured by (top to bottom): 10x library prep protocol, donor ID, cellular compartment. (**B**) heat map of confusion matrices between Leiden clusters and previously annotated cell type labels, with clustering on BBKNN integration (left, 37 clusters) or scVI integration (right, 75 clusters) of the full dataset (clustering resolution = 1.5). (**C**) heat map of confusion matrices between Leiden clusters and newly annotated cell type labels, with clustering on BBKNN integration (left, 37 clusters) or scVI integration (right, 75 clusters) of the full dataset (clustering resolution = 1.5). For each confusion matrix, the Normalized Mutual Information (NMI) score between cluster labels and annotation labels is shown.

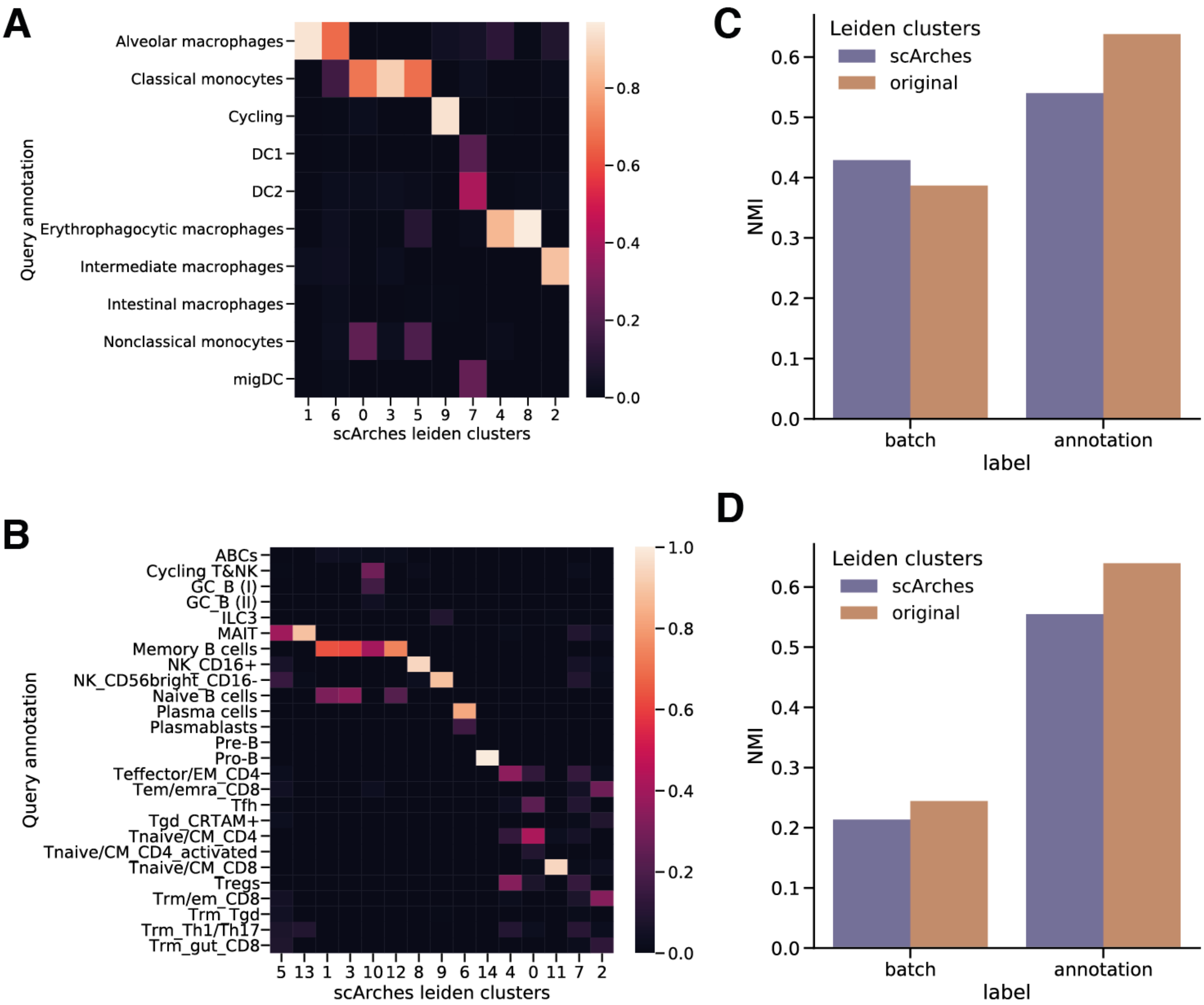


**fig. S25: Validation of biological conservation after scArches mapping of adult cells to prenatal reference.** (**A, B**) Heat maps of confusion matrix between adult immune cell annotations from Conde et al. (2020) and Leiden clusters obtained with latent dimensions after mapping adult cells on the prenatal reference with scArches, for myeloid cells (A) and lymphoid cells (B). (**C, D**) Barplots of Normalized Mutual Information (NMI) between technical batch or cell type annotation and Leiden clusters from scArches mapping or the clusters in the BBKNN embedding from Conde et al.

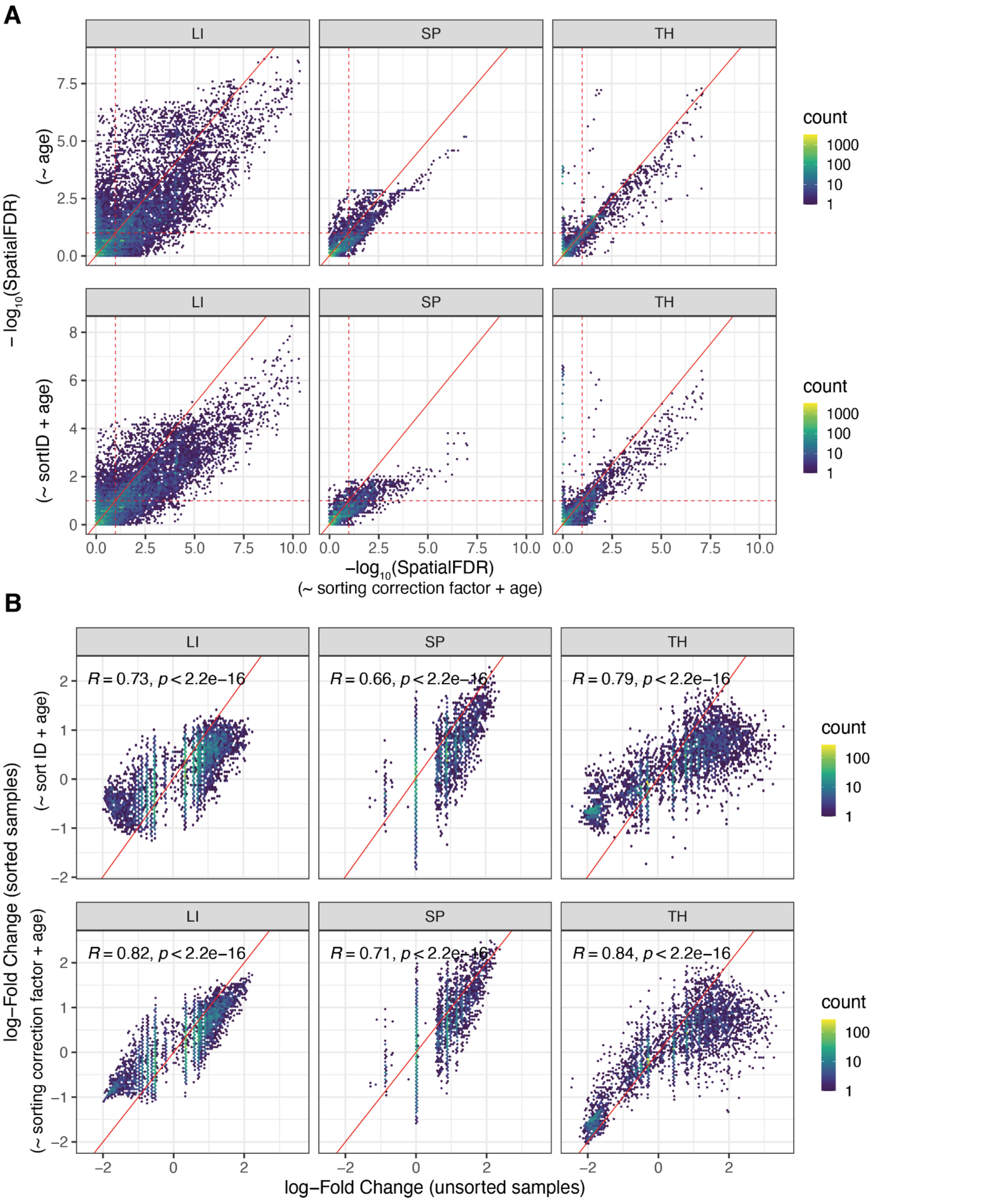

A
LI
SP
TH
$-\log_{10}$(SpatialFDR)
(~ age)
(~ sortID + age)
count
1000
100
10
1
$-\log_{10}$(SpatialFDR)
(~ sorting correction factor + age)
B
$R = 0.73$, $p < 2.2e-16$
$R = 0.66$, $p < 2.2e-16$
$R = 0.79$, $p < 2.2e-16$
$R = 0.82$, $p < 2.2e-16$
$R = 0.71$, $p < 2.2e-16$
$R = 0.84$, $p < 2.2e-16$
log−Fold Change (sorted samples)
(~ sort ID + age)
(~ sorting correction factor + age)
log−Fold Change (unsorted samples)

**fig. S26: Validation of quantification of FACS sorting effect on cell abundances in Milo neighbourhoods.** (**A**) Scatter plot of SpatialFDR (in - $\log_{10}$ scale) estimated in test for differential abundance across gestational age, regressing out continuous FACS sorting factor (x-axis), without regressing out FACS effect (y-axis, top) and regressing out FACS protocol label ($CD45^{+}$/$CD45^{-}$/unsorted) (y-axis, bottom). Results for the test on cells from liver (LI), spleen (SP) and thymus (TH) are shown. The dotted red lines indicate the significance threshold of 10% SpatialFDR. (**B**) Scatterplot of log-Fold change estimated in test for differential abundance across gestational age testing on the subset of unsorted samples (x-axis) and on FACS sorted samples (y-axis), regressing out the FACS protocol label (top) or the FACS sorting factor (bottom).

### Supplementary Tables

**table S1: Table_S1.csv (separate file)**
Differential expression analysis results for test on gestation-stage specific macrophage and mast cells neighbourhoods

**table S2: Table_S2.csv (separate file)**
Differential expression analysis results for comparison of bone marrow and peripheral CCR2hi monocytes

**table S3: Table_S3.csv (separate file)**
Differential expression analysis results for test on gestation-stage specific NK cells neighbourhoods

**table S4: Table_S4.csv (separate file)**
Differential expression analysis results for comparison of thymic and peripheral mature T cells
